# Eyespots originated multiple times independently across the Lepidoptera

**DOI:** 10.1101/2024.02.07.579046

**Authors:** Brian Hanotte, Beatriz Willink, Antónia Monteiro

**Affiliations:** Department of Biological Sciences, National University of Singapore, Singapore; Department of Zoology, Stockholm University, Sweden; Department of Entomology, Cornell University, Ithaca, NY 14853, USA

## Abstract

Eyespot colour patterns are an effective defence against predators and are found in many animal lineages. In nymphalid butterflies, eyespots have a single evolutionary origin close to the base of this clade, but eyespots are also present in many other lepidopteran lineages and may have multiple independent origins. Here we use phylogenetic comparative methods to investigate the evolution of eyespots across a multi-superfamily phylogeny of Lepidoptera, and to pinpoint lineages in which eyespots likely originated independently. We find a total of 32 separate origins of Discal eyespots (in the discal wing region) and 21 separate origins of Marginal eyespots (in the marginal wing region). In two instances, eyespots were preserved in most extant representatives of a subsequent species radiation. In one such instance, in the Nymphalidae, ancestral state reconstructions suggest that an ancestor with Marginal eyespots predated the origin of one or more lineages with Discal eyespots, while in the Saturniidae we observed the opposite. We conclude that eyespots do not appear to be homologous across the Lepidoptera. Our phylogenetic inference provides a roadmap for future developmental and functional studies that can address whether Discal and Marginal eyespots share the same gene-regulatory network. This study, therefore, has implications for our understanding of the evolution of serial homologues and of convergent evolution of visual signals in insects.

## Introduction

Adaptive traits in animals such as protective colour patterns have long been a source of interest and research (Tylor, 1886; Poulton, 1890; Beddard, 1895). Stripes, spots, and blotches can all function as protective colour patterns (Stevens, 2005; Caro et al. 2016; Mizuno et al. 2024), but the ocellus, or eyespot, which is characterized by the presence of one or more concentric rings of contrasting colours, is among one of the most researched protective patterns. Eyespots are found across a wide variety of vertebrate and invertebrate species (Edmunds, 1976; Meadows, 1993; Loyau et al. 2007; He et al. 2010; Wong & Marek, 2019; Hernández-Palma et al. 2023) and represent a textbook example of evolutionary convergence in response to selection by visual predators (Stevens et al. 2008; Oliver et al. 2012; Cunha et al. 2023).

In Lepidoptera (butterflies and moths), eyespots occur in a variety of locations on the body of larvae and/or the wings of adults and have been implicated in multiple ecological contexts. In both larvae and adults, eyespots are used in predator deterrence and in attack deflection, but in adults they can also function in courtship and mate selection (Breuker & Brakefield 2002; Robertson and Monteiro 2005; Vallin et al. 2007; Kodandaramaiah, 2011; Merilaita et al. 2011; Prudic et al. 2011; Kodandaramaiah et al. 2013; Hossie et al. 2014; Prudic et al. 2015; Mukherjee & Kodandaramaiah, 2015; Huq et al. 2019; Halali et al. 2019; Crees et al. 2021).

In adult wings, eyespots can be found on different regions and will be named, hereafter, as Marginal and Discal eyespots. Marginal eyespots develop close to the wing margin in between veins (Fig. 1A-B), whereas Discal eyespots are found more centrally located on the wing, either straddling the cross vein in the discal cell region or within the discal cell (Fig. 1 C-D; Otaki et al. 2020). Lepidopterans can have one or both types of eyespots on their wings. Marginal eyespots are often found in more than one unit on the wing margins and are considered serial homologs, whereas Discal eyespots are often found as single units. It is unclear whether Discal eyespots are serial homologs to Marginal eyespots, i.e., whether these two eyespot types are using the same gene-regulatory network (GRN). Some nymphalid Marginal eyespot genes, such as *Distal-less* and *engrailed* are expressed in saturniid Discal eyespots (Monteiro et al. 2006). Additionally, *Distal-less* knockouts in *Junonia coenia* changed both Marginal and Discal eyespots (Zhang and Reed, 2016). These data, however, are still too scant to support GRN homology.

The evolutionary history of Discal and Marginal eyespots across the Lepidoptera is also still largely unknown, as most research on eyespot evolution has focused on Marginal eyespots of brush-footed butterflies. This clade of butterflies (Nymphalidae), which contains ∼6,400 described species, accounts for only ∼3.3% of lepidopteran diversity (∼180,000 total described Lepidoptera; Brakefield, 2003; Heikkilä et al. 2015; Monteiro, 2015; Bhardwaj et al. 2020; Chazot et al. 2021). Phylogenetic comparative methods estimated that Marginal eyespots evolved once in the Nymphalidae, shortly after the origin of the clade (Oliver et al. 2012; Oliver et al. 2014). Thus, all Marginal eyespots in extant nymphalids are thought to be homologous. Nonetheless, Marginal eyespots are found in taxa across the entire order (e.g., Semanturidae, Saturniidae, Cambridae), and thus it is likely that these colour patterns have evolved independently in multiple lineages of this diverse insect clade. Similarly, Discal eyespots are widespread in Lepidoptera and have not been studied in a phylogenetic context (Martin and Reed, 2010). Because these two wing regions express different genes during development (Banerjee et al. 2023), Marginal and Discal eyespots might have evolved largely independently of each other. If so, we should expect the two eyespot forms to occur primarily in different lineages (Fig. 1E). Alternatively, eyespots might have first evolved on one wing region and later gained a novel domain of expression on the other region, with the order of these regions potentially varying (Fig. 1F-G). Here, we aim to investigate the evolutionary history of lepidopteran eyespots beyond Nymphalidae, to understand the diversity of paths that resulted in the morphological convergence of this complex colour pattern.

**Figure 1.**
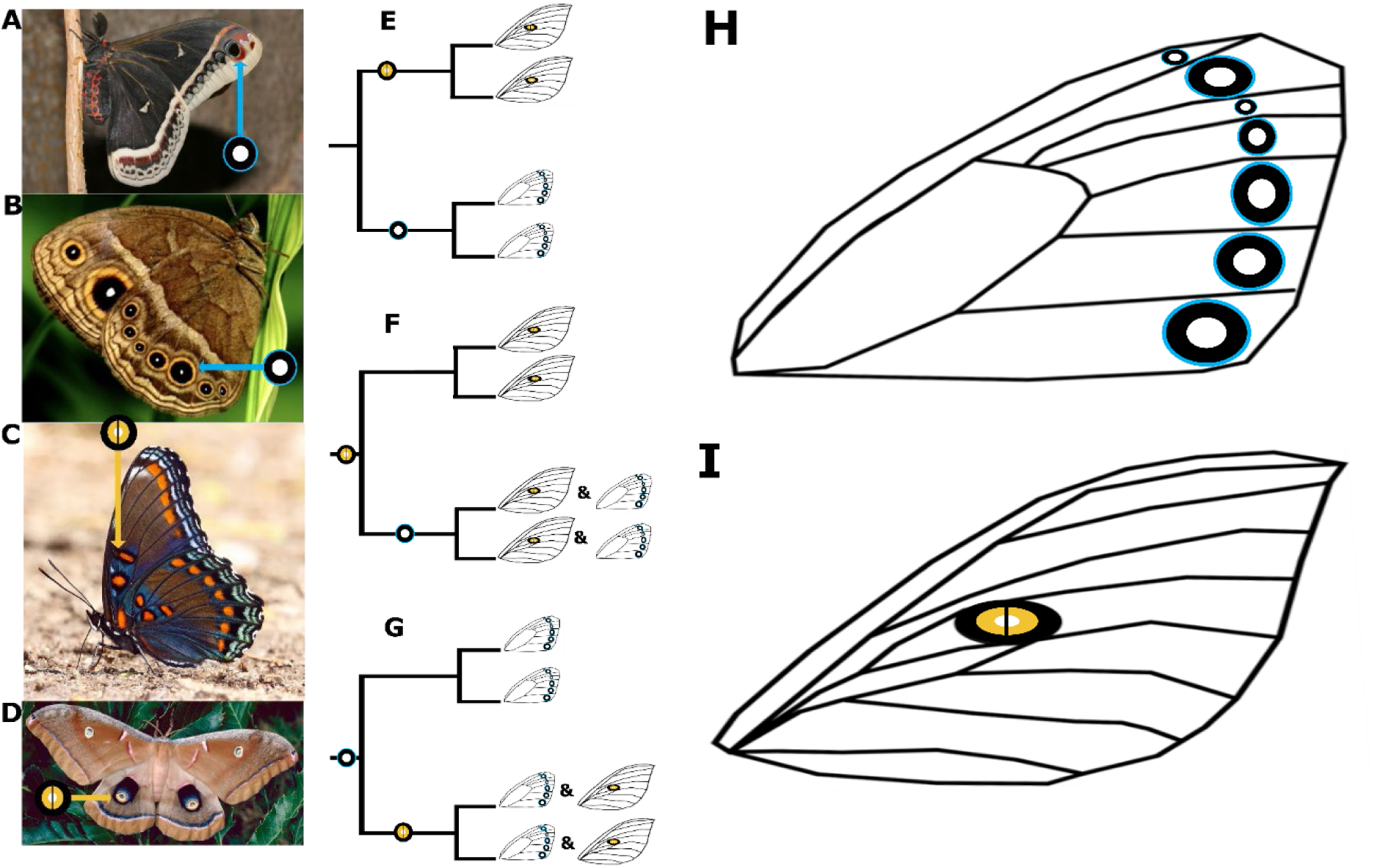
Lepidopteran eyespots have evolved in different wing regions: **A-B** the Marginal region, between veins and **C-D** the Discal region, straddling the cross vein in the discal cell or found within the cell. Eyespots could evolve in one of the following possible orders (E-G): **E** These two eyespot types may have evolved independently of each other and thus be primarily found in different clades (multiple origins). Alternatively, **F** Discal eyespots may evolve first followed by Marginal eyespots, or **G** Marginal eyespots may enable the subsequent evolution of Discal eyespots. **H** Marginal eyespots. **I** Discal eyespots. Photographs: **A** = *Eupakardia calleta* (Nicky Davis, WildUtah.US.), **B** = *Bicyclus anynana* (Wiliam Piel, National University of Singapore), **C** = *Limenitis arthemis* (Rawpixel, Rawpixel.com), **D** = *Antheraea polyphemus* (Lacy L. Hyche, Auburn University, Bugwood.org).

We used modern phylogenetic methods to investigate the evolutionary origins of Discal and Marginal eyespot across Lepidoptera. We first inferred a species-level phylogeny for Lepidoptera, sampling 5 to 27 molecular sequences per species across 715 taxa. Our phylogeny covers 65% of all superfamilies in the order Lepidoptera, allowing comparative inferences of eyespot evolution in moths and butterflies. We then modelled evolutionary origins and losses of eyespots and inferred ancestral states across the phylogeny. Additionally, for each subtree including multiple (two or more) eyespot-bearing taxa, we implemented a model comparison approach, based on marginal likelihood estimation, to quantify support for an ancestral origin of eyespots in the most recent common ancestor of the subtree. If an ancestral origin of eyespots was supported, we tentatively concluded that eyespots in all extant taxa of the subtree are likely homologous. These putative ancestral eyespot origins were then compared with the ancestral state estimates above, to evaluate the robustness of our results from tree-wide *versus* subtree approaches.

Our results support the hypothesis that eyespots in Lepidoptera have evolved multiple times independently. Our data also suggests that Discal and Marginal eyespots have evolved in different temporal sequences in the two main clades where they radiated (Nymphalidae and Saturniidae). We find that both Marginal and Discal eyespots have evolved in 11 superfamilies and numerous families across the lepidopteran tree. Finally, we find that eyespots are more commonly observed in the Discal wing region (32 independent origins) than the Marginal region (21 independent origins). Together, our results suggest that eyespot evolution is commonplace in Lepidoptera. Our study provides a phylogenetic roadmap for the future sampling of independent lineages to test whether all these eyespots share a common gene-regulatory network.

## Results

### Tree topology

While large-scale multi-superfamily phylogenetic trees of Lepidoptera have been inferred in several recent studies (Heikkilä et al. 2015; Mitter et al. 2017; Kawahara et al. 2019), these studies focused on resolving deep ancestral nodes within Lepidoptera and sampled taxa accordingly. We aimed, instead, to maximize taxonomic coverage across the entire clade and thus enable large-scale comparative analyses of eyespot evolution, by reconstructing a new phylogeny of Lepidoptera. We focused on Ditrysia, a natural clade that includes 99% of extant species (Heikkilä et al. 2015) and omitted more basal branching taxa due to unavailable wing images. We sampled 5 to 27 gene fragments per species across 715 taxa and conducted maximum likelihood phylogenetic inference using a generalized time reversible (GTR) substitution model with gamma-distributed site heterogeneity upon the concatenated alignment. Our inferred tree topology (Fig. 2) is largely congruent with previous phylogenies (Heikkilä et al. 2015; Mitter et al. 2017; Kawahara et al. 2019, Wright et al., 2026).

Topological differences observed between this study and previous ones involved species whose placement is phylogenetically uncertain (*incertae cedis*), or superfamilies previously shown to require reclassification (i.e. Tineoidea or the Cossioidea-Sesioidea complex, among others; Mutanen et al. 2010; Bazinet et al. 2013; Heikkilä et al. 2015 Reiger et al. 2015; Mitter et al. 2017). These differences in topology are unlikely to have a significant impact on the main findings of this work, as they are primarily nested within large clades entirely lacking eyespots. A summary and discussion of taxonomic relationships identified in this work can be found in our Supplementary Material (see *Tree Topology*).

**Figure 2.**
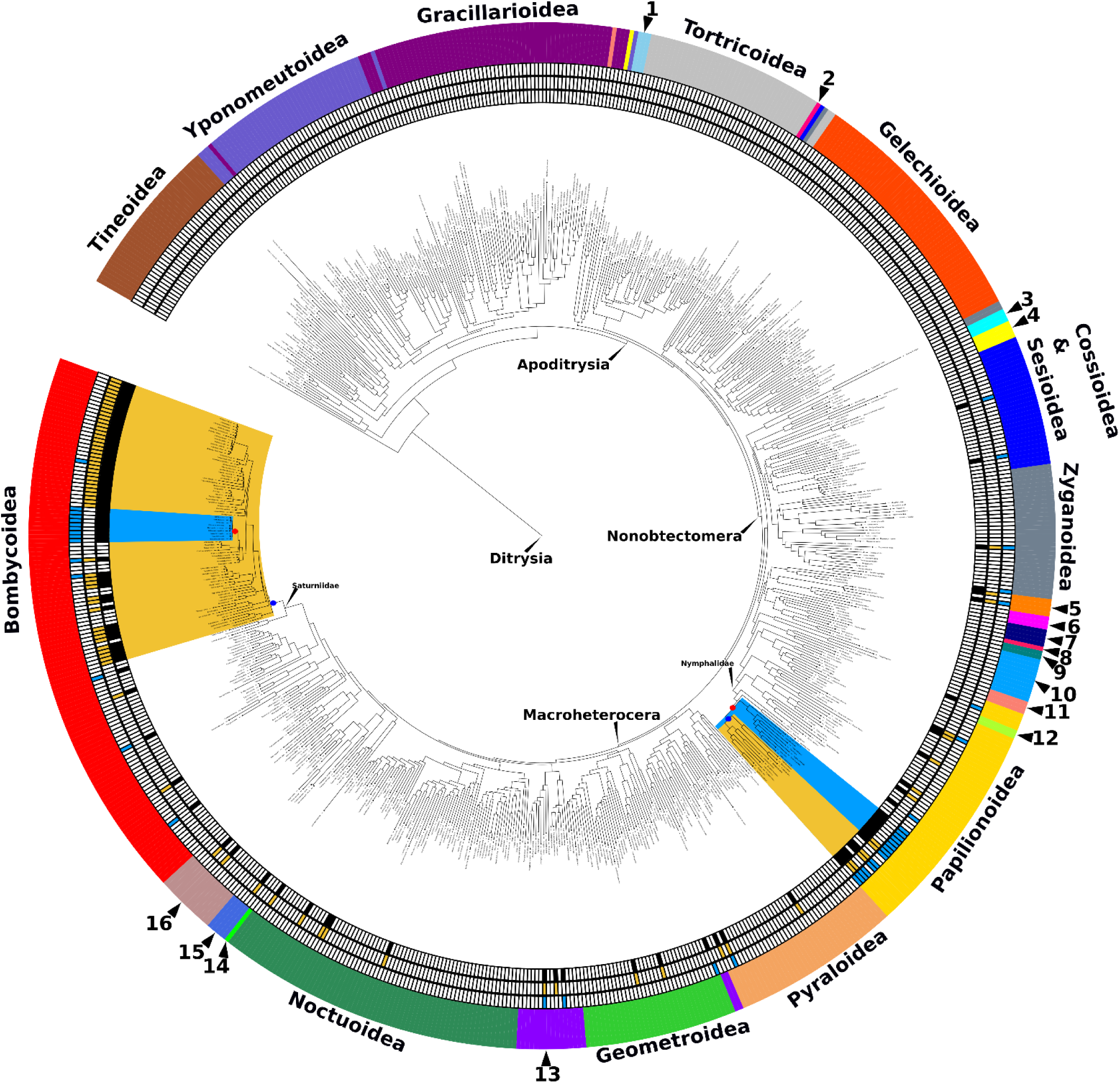
Phylogeny of Ditrysia comprising 715 species from 28 superfamilies. The superfamilies are outlined on the outer circumference of the tree. Superfamilies which are too small to be written are numbered as following: 1 = Urodoidea, 2 = Galacticoidea, 3 = Immoidea, 4 = Choreutoidea, 5 = Calliduloidea, 6= Epermenoidea, 7 = Carposinoidea, 8 = Copomorphoidea, 9 = Hyblaeoidea, 10 = Thyridoidea, 11 = Pterophoroidea, 12 = Hedyloidea, 13 = Drepanoidea, 14 = Cimelioidea, 15 = Mimallonoidea, 16 = Lasiocampoidea. The mostly likely origins of eyespots in Nymphalidae and Saturniidae, based on ancestral state reconstruction, are shown by red (Marginal) and blue (Discal) circles. Descendant clades are shaded in in blue (Marginal) and yellow (Discal). Colour bars plotted outside the phylogeny indicate eyespot presences on extant taxa: black = any kind of eyespot, blue = Marginal eyespots, and yellow = Discal eyespots.

**Figure 3.**
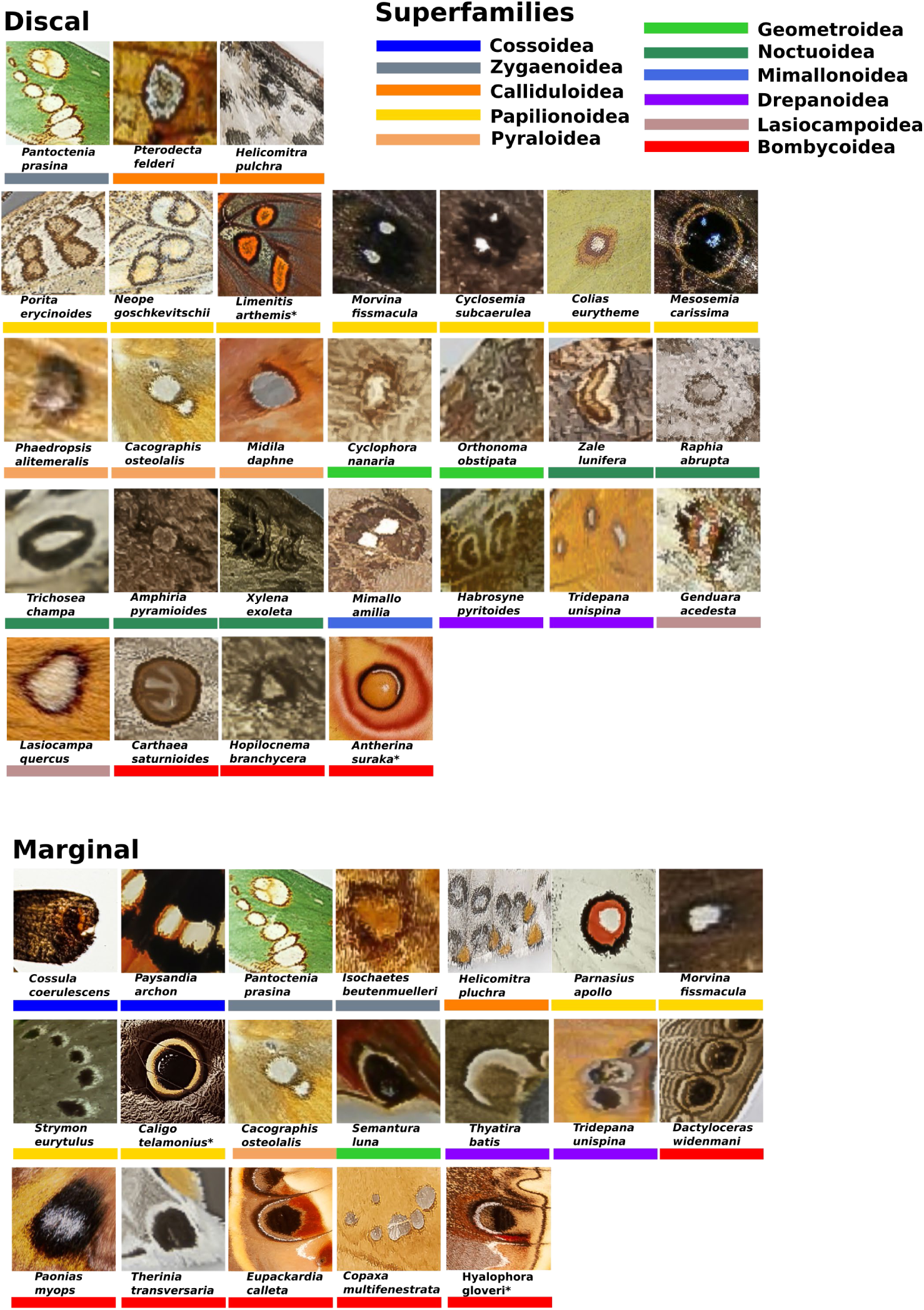
Examples of eyespot occurrences across Ditrysia superfamilies, illustrating the criterion used to define eyespot wing patterns in this study. Species marked with a * are representatives for two taxonomic groups (Bombycoidea:Saturniidae and Papilionoide:Nymphalidae), in which our analyses suggest both Discal and Marginal eyespots have evolved, albeit in distinctly orderded sequences (Fig. 1F,G, see Results). Colours in the key represent the corresponding superfamilies in Figure 2.

### Ancestral state reconstructions and evolutionary rates

To investigate the historical evolution of eyespots across Ditrysia, we obtained photographs of as many species as possible across the phylogeny (only 4 species had missing images) and scored presence or absence of Discal and/or Marginal eyespots anywhere on the wings of either sex (Fig. 2). Eyespots were defined as wing patterns containing a circular or oblong spot encircled by one or more concentric rings of a distinct colour that also contrast with the overall background colour of the wing (Fig. 3). We also scored eyespots using a ‘relaxed’ definition, where non-circular/oblong shapes (e.g., triangles, rectangles, teardrops, Figs. S1, S2) encircled by another ring of colour were also considered. Relaxing the eyespot criteria increased the number of taxa classified as displaying Discal eyespots but did not affect the scoring of Marginal eyespots. Here, we focus on the results from analyses under the ‘strict’ eyespot criteria, and report results of analyses based on the more inclusive eyespot definition in the Supporting Material.

We found that 41 species have Marginal eyespots and 82 species have Discal eyespots (100 species under the ‘relaxed’ eyespot definition). A total of 15 species have both eyespot patterns on their wings across our data set (25 species under the ‘relaxed’ eyespot definition). Eyespots were absent outside the Nonobtectomera (Tineoidea - Galacticoidea) and were not observed in the nonobtectomeran Gelechioidea (Fig. 2). We modelled eyespot evolution as a discrete trait with two alternative states, presence or absence, regardless of eyespot location on the wings, using RevBayes v1.3.0 (Höhna et al. 2016). We monitored joint conditional ancestral states in two independent MCMC simulations and used stochastic character mapping (Freyman and Höhna, 2019) to estimate the number of eyespot gains and losses across the Ditrysia tree. We then repeated these analyses for Marginal and Discal eyespots separately, to determine if these distinct eyespot types have radiated in different clades and in different temporal sequences.

Our three ancestral state reconstructions (for the two eyespot types combined, for Marginal and for Discal eyespots) indicated that both the most recent common ancestor (MRCA) of all Ditrysia as well as the MRCA of all Nonobtectomera (including and excluding the Gelechioidea) did not have eyespots (Figs. 4, S3 – S6). Eyespots, regardless of where they occur, most likely evolved multiple times independently. We see this through several potential eyespot origins across the Lepidoptera (Table 1). The data suggest that eyespots may not persist long evolutionarily, with the rate of losses exceeding the rate of origins in the analyses on Discal eyespots and the analyses on both eyespot types combined (Figs. 4, S3 – S6).

Eyespot-bearing species are distributed across 11 superfamilies, but in most cases, eyespots occur in only one genus or a few genera, within a superfamily. Because taxonomic sampling remains sparse (22.60% of genera on average across all superfamilies in this study), it is currently difficult to discern how phylogenetically widespread eyespots are in most of these superfamilies. There are, however, two major independent occurrences of Marginal and Discal eyespots. In both Nymphalidae and Saturniidae a single common eyespot ancestor is shared across multiple species in the family, and both eyespot types have been retained in multiple lineages after their origin (Figs. 4, S3-S6).

**Table 1.** Number of independent eyespot origins based on stochastic character maps from discrete-trait models of the evolution of any eyespot type (irrespective of location), Discal eyespots, and Marginal eyespots. For each model, we report the median number of origins across the posterior, and the 95% highest posterior density (HPD) interval.

| Eyespot type | Independent origins [95% HPD interval] |
| --- | --- |
| Discal | 32 [28-36] |
| Marginal | 21 [21-23] |
| Any eyespot type | 38 [34-42] |

**Figure 4.**
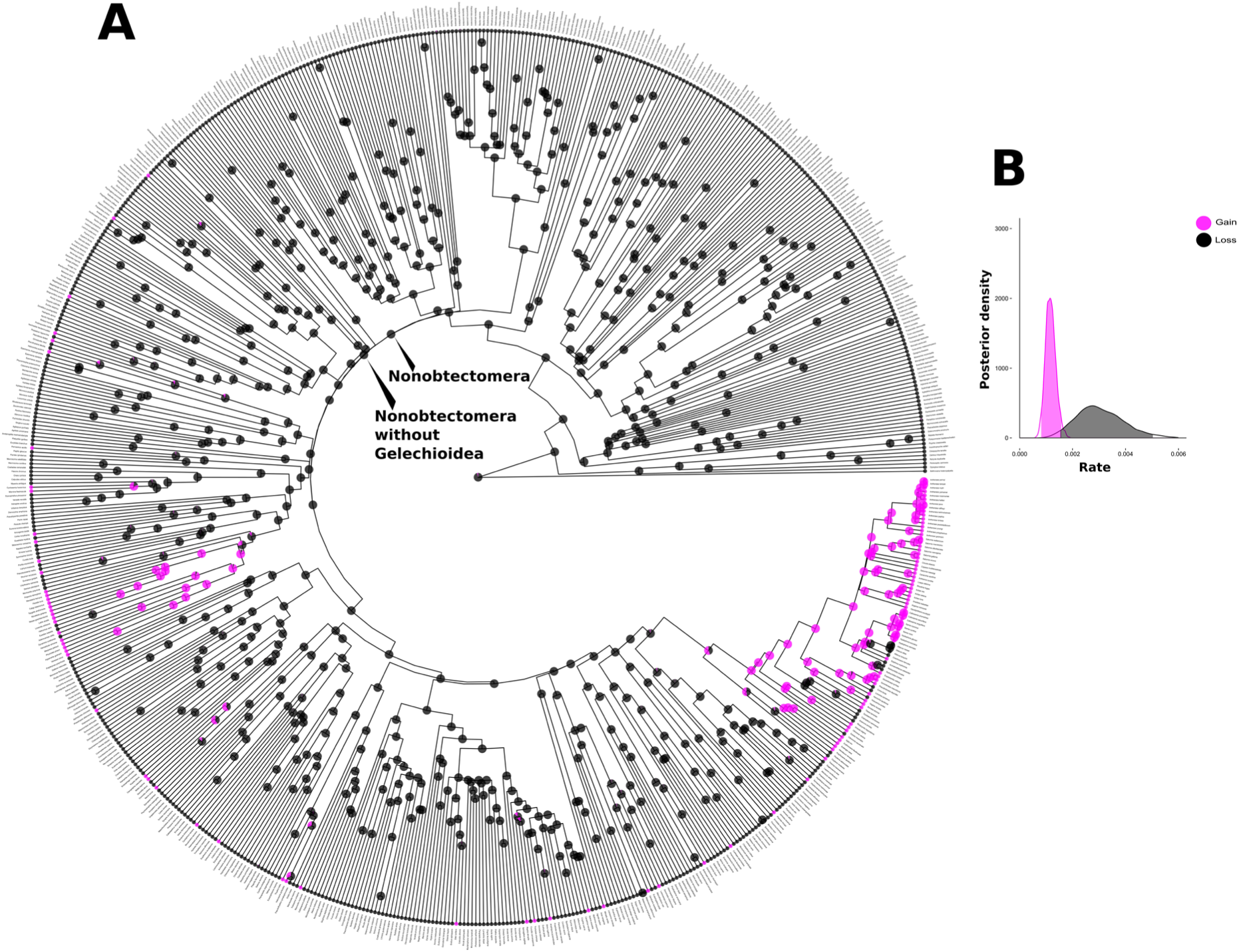
Eyespot ancestral state reconstruction and transition rates in Lepidoptera for Marginal and Discal eyespots combined. We modelled eyespot evolution as a discrete trait with two alternative states (presence or absence) regardless of marginal or discal location. **A)** Phylogeny of the Lepidoptera with ancestral state reconstructions. Pies indicate the posterior probability of eyespot presence for each internal node. The posterior probability of eyespot presence at the root of Lepidoptera was 0.038. The posterior probability of an eyespot at the base of the Nonobtectomera was 0, and finally the posterior probability of an eyespot within the Nonobtectomera excluding the Gelechioidea but including all present-day eyespot species was 0. **B)** Posterior distributions of rate estimates for eyespot gain and eyespot loss across Lepidoptera. Higher rate values (i.e. a more rightward distribution) indicate a shorter waiting time for an evolutionary transition (eyespot gain or loss). Separate analyses for Marginal and Discal eyespots and analyses under a ‘relaxed’ definition of eyespots are shown in the Supporting Material (Figs. S3-S6).

### Homology of eyespots with a single common ancestor – common ancestor model vs the multiple origins model

To validate ancestral state reconstructions pointing to independent eyespot origins, we conducted a separate test of eyespot homology for each subclade of our Ditrysia tree including at least two eyespot-bearing taxa (Fig. S7). These tests contrasted the hypothesis of a single ancestral origin of eyespots in the most recent common ancestor of the subclade (hereafter the ‘common ancestor’ model) against the hypothesis of two or more independent eyespot origins within the subclade (hereafter the ‘multiple origins’ model). Support for the ‘common ancestor’ model would be suggestive of eyespot homology across all extant species in the subclade, especially if congruent with the results of tree-wide ancestral state reconstructions (Fig. 4), which allowed for eyespot loss and regain.

The ‘common ancestor’ and ‘multiple origins’ models were parametrized as our eyespot evolution models used for ancestral state reconstructions above (Fig. 4, see also Methods), except for one key difference. Here, we specified alternative priors on the root state frequencies to enforce certainty of an eyespot presence at the root of each subclade in the ‘common ancestor’ model, and to enforce certainty of a common ancestor without eyespots in the ‘multiple origins’ model. For each subclade, we then estimated the marginal likelihoods of these two models and compared them using Bayes factors. Marginal likelihood estimates indicate the likelihood of a model, including all parameters and priors, given the data. By comparing alternative models that differ only by their prior beliefs on the character state at the root (presence or absence of eyespots), we thus tested if the assumption of eyespot homology across extant taxa is more likely, given the phylogeny and character data, than the assumption of independent eyespot origins.

We found support for Discal eyespot homology at the base of the Saturniini tribe of giant silkmoths (Saturniidae, Table 2). The same node was recovered as a shared ancestor bearing eyespots in the analysis where Discal and Marginal eyespots were combined (Table 2), consistent with an earlier origin of Discal eyespots, relative to Marginal eyespots, in Saturniini. In contrast, Marginal eyespots in silkmoths were only weakly supported in one ancestral node within the tribe Attacini, giving rise to the genera *Callosomia*, *Samia,* and *Hyalophora* (Table 2).

These results are consistent with our tree-wide ancestral state reconstructions (Fig. 4). However, our homology inference did not account for all current occurrences of Discal eyespots in Saturniidae, suggesting additional or more ancestral origins, which our analysis may not have been sufficiently powered to detect. Similarly, a single ancestral origin of Marginal eyespots within Attacini could not explain the current distribution of these eyespot patterns across Saturniidae, indicating that additional origins or repeated losses resulted in a few scattered descendants in the tribe Saturniini, and in the subfamily Oxytenninae (Fig. 2). Notably, when we relaxed the eyespot-defining criterion to include non-circular or oblong shapes, a more ancestral origin of Discal eyespots was inferred with strong support in the common ancestor of Saturniini + Attacini (Table 2).

In the Nymphalidae, Marginal eyespots are likely homologous (Fig. 4, Table 2, Fig. S4). Within Nymphalidae, the MRCA of the clade excluding Danainae and Libytheinae, was inferred as the most likely common ancestor of both Marginal eyespots and all eyespots, regardless of wing location (Figs. 4, S4, Table 2). In contrast, our model-comparison approach did not provide evidence for homologous Discal eyespots in brush-footed butterflies (Figs. 4, S4, Table 2). While Discal eyespots may be relatively common in Nymphalidae (six out of 20 species sampled), a robust inference of their evolutionary origins within the clade will likely require a more densely sampled phylogeny.

Together, our ancestral state reconstructions, and model-comparison approach point at two taxonomic groups containing eyespot-bearing lineages with contrasting evolutionary histories: the Saturniidae (either Saturniini or a more inclusive clade based on the ‘relaxed’ eyespot criterion), where Discal eyespots likely evolved first and were followed by a few origins of Marginal eyespots, and the Nymphalidae, where Marginal eyespots are more likely to have evolved first, followed by one or more origins of Discal eyespots (Figs. 2, S3 – S6).

**Table 2.** Giant silkmoth and brush-footed butterfly clades with highest support for an ancestral origin of eyespot patterns. For each clade, we report the statistical support (log_10_ BF) for the ‘common ancestor’ model of eyespot evolution, compared to the ‘multiple-origins’ model (see Methods). A conventional interpretation for log_10_ BF values is: 0-0.5 = evidence barely worth mentioning, 0.5-1 = substantial evidence, 1-2 = strong evidence, and > 2 = decisive evidence (Jeffreys, 1961). However, our simulation approach revealed that the absolute log_10_ BF values tend to decrease with tree size, regardless of the of the rate of false positives. For example, an incorrect log_10_ BF value in support of the ‘common ancestor’ model, occurred at rates well below 5% for trees with 10 extant taxa, even though the mean log_10_ BF value for the correct ‘multiple-origins’ model was only 0.2 for these small trees ( see *Inference of ancestral eyespot origins: validating the model-comparison approach* in the Supplementary Material ). We therefore cautiously report log_10_ BF values between 0.4-0.5 in a clade within Nymphalidae, including 18 species sampled in our phylogeny, and log_10_ BF values > 0.2 in clade within the tribe Attacini, which includes 9 species sampled in our phylogeny.

| Eyespot type | Family | Clade | $\log_{10}$ BF for common ancestor |
| --- | --- | --- | --- |
| All | Giant silkmoths (Saturniidae) | Saturniini (Saturniinae)<sup>1</sup> | 1.3 |
| All | Brush-footed butterflies (Nymphalidae) | All excluding Danainae and Libytheinae | 0.4 |
| Discal | Giant silkmoths (Saturniidae) | Tribe Saturniini (Saturniinae:Saturniinae)<sup>2</sup> | 1.4 |
| Discal | Brush-footed butterflies (Nymphalidae) | NA<sup>3</sup> | NA |
| Marginal | Giant silkmoths (Saturniidae) | *Callosomia*, *Samia*, *Hyalophora* (Saturniidae:Attacini) | 0.2 |
| Marginal | Brush-footed butterflies (Nymphalidae) | All excluding Danainae and Libytheinae | 0.4 |

<sup>1</sup>The tribe Saturniini here includes the genus *Rhodinia*, which follows Rougerie et al., (2022).
<sup>2</sup>Under the 'relaxed' eyespot criterion, a more inclusive clade comprising both Saturniini and Attacini is strongly supported with log<sub>10</sub> BF = 1.9.
<sup>3</sup>Even though six species out of 20 in Nymphalidae display Discal eyespots, no common ancestor was supported with log<sub>10</sub> BF value > 0.2.

### Correlated evolution of Marginal and Discal eyespots

Our finding of two subclades, Nymphalidae and Saturniidae, with both Marginal and Discal eyespots likely arising in different temporal orders (Fig. 2) led to the hypothesis that eyespots may initially evolve on a single wing domain (Marginal or Discal), but once the genetic machinery for eyespot development becomes established, eyespot evolution on a different wing domain is facilitated.

Critically, this novel-domain hypothesis is compatible with either wing region hosting the most ancestral eyespot, but it requires evidence that the ancestral origin facilitated eyespot evolution in novel domains, as would be expected if the eyespot gene-regulatory network is redeployed in a different group of cells as opposed to evolving anew.

We addressed this hypothesis using a correlated evolution model, wherein the rate of eyespot evolution on each wing region depends on whether eyespots are already present on the alternative region. To test for correlated evolution of eyespots between wing regions (Marginal and Discal), we modelled a reversible jump mixture distribution, sampling the posterior probabilities of correlated vs independent eyespot evolution models and assuming equal prior probabilities for both scenarios. If correlated evolution is supported, the origin of novel Marginal and Discal eyespots should be accelerated when eyespots already exist in the alternative wing domain, whereas no such acceleration should be present if eyespot gains are independent. We tested this hypothesis using pruned phylogenies for the superfamilies Papilionoidea (containing Nymphalidae) and Bombycoidea (containing Saturniidae), as both clades encompass multiple putative origins of both Discal and Marginal eyespots (Figs. S7 – S9) and therefore enable estimation of evolutionary rates on each eyespot domain.

Our results do not strongly support correlated evolution between Marginal and Discal eyespots. In Bombycoidea, where we expected the origin of Marginal eyespots to be contingent on the pre-existence of Discal eyespots, our results were inconclusive; the correlated and independent evolution models were each supported in about 50% of the posterior (Fig. S8, S9). This ambiguity may have arisen from the uncertain character state at the base of the giant silkmoth tribe Attacini, which currently includes several species with only Marginal eyespots (Fig. 5A). In our correlated evolution model, the MRCA of Attacini was reconstructed as ambiguous between having no eyespots or Discal eyespots only (Fig. 5A).

In Papilionoidea, the correlated evolution model, wherein the origin of Discal eyespots depends on the pre-existence of Marginal eyespots, was weakly supported, accounting for about 71% of the posterior and a log_10_BF = 0.39 (Fig. S10). Consistent with our hypothesis, Discal eyespots were inferred to arise at a higher rate in lineages bearing Marginal eyespots (Fig S11). Specifically, we estimated, through stochastic character mapping, a median of 4 (95% HPD interval = 2-7) *de novo* origins of Discal eyespots in Papilionoidea. In contrast, a median of 18 (95% HPD interval = 3-24) origins of Discal eyespots were estimated in lineages within Papilionoidea, already bearing Marginal eyespots. Ancestral state reconstructions suggested, with modest posterior probability, that nymphalid species bearing both Marginal and Discal eyespots may have shared an ancestor with only Marginal eyespots (Fig. 5B). We also note that modelling the joint evolution of both eyespot types resulted in inferred ancestral origins that were largely consistent with our previous tree-wide ancestral state reconstructions of individual eyespot domains (compare Fig. 5 and Figs. 4, S3-S6).

**Figure 5.**
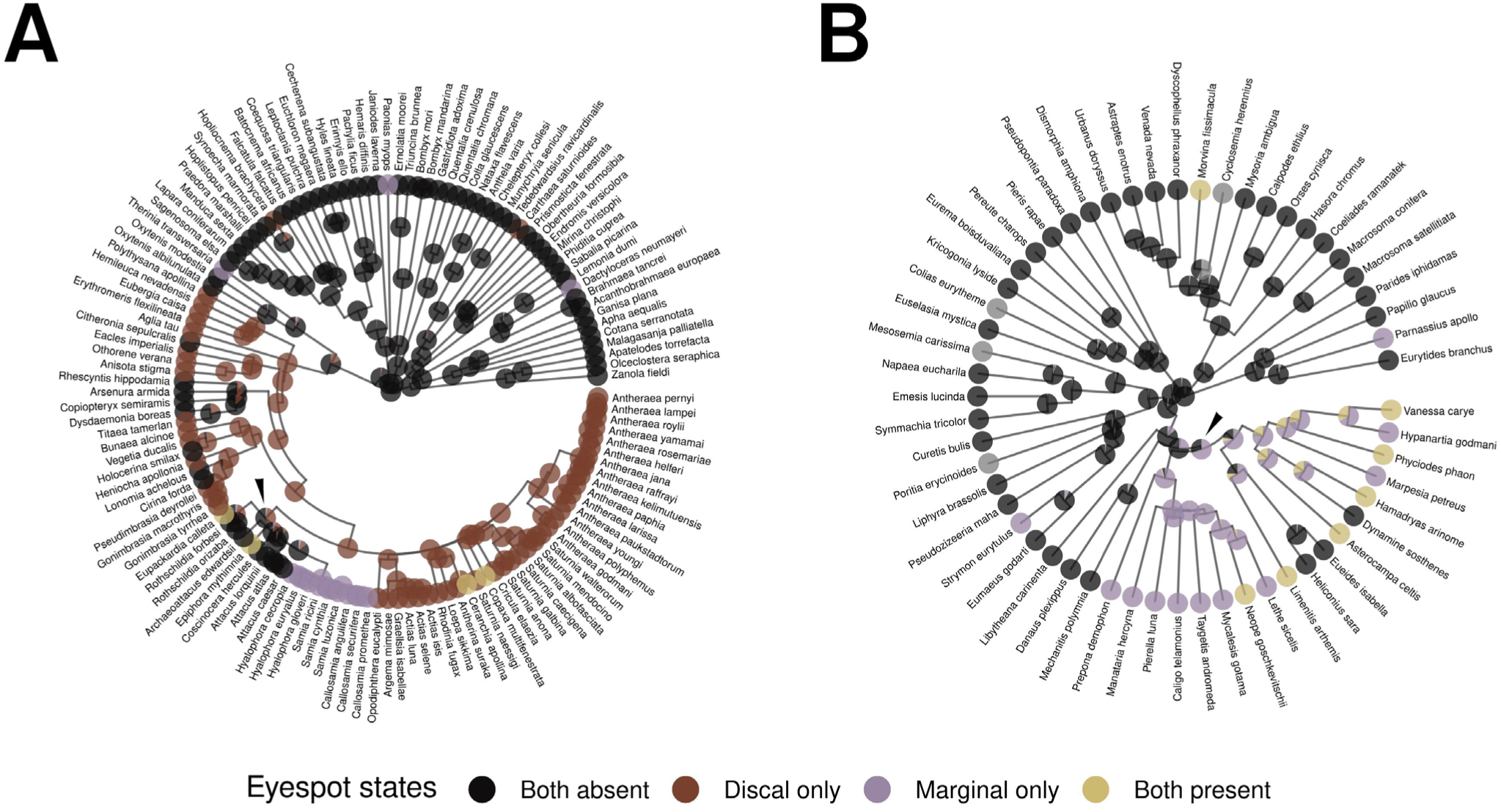
Joint evolution of Marginal and Discal eyespots. We modelled the evolution of Marginal and Discal eyespots eyespot while allowing character states (presence or absence) in one eyespot type to influence transitions in the other (the correlated evolution model), and inferred ancestral states for both eyespot types. Pies indicate the posterior probability of eyespot presence for each internal node **A)** Bombycoidea: the arrowhead indicates the common ancestor of the Saturniinae tribe Attacini. **B)** Papilionoidea: the arrowhead indicates a common ancestor likely displaying Marginal eyespots only and giving rise to species with both Marginal and Discal eyespots.

### Other smaller eyespot observations

Our stochastic character mapping analyses suggested that both Discal and Marginal eyespots have evolved multiple times in Ditrysia (Table 1), with only a few of these origins occurring in Nymphalidae and Saturniidae. The remaining origins of Discal and Marginal eyespots, occurred in sparsely sampled families (superfamilies), namely: Brahmaeidae (Bombycoidea), Sphingidae (Bombycoidea), Carthaeidae (Bombycoidea), Cossidae (Cossioidea-Sesioidea), Castniidae (Cossioidea-Sesioidea), Drepanidae (Drepanoidea), Geometridae (Geometroidea), Sematuridae (Geometroidea), Hesperiidae (Papilionoidea), Lycaenidae (Papilionoidea), Papilionidae (Papilionoidea), Pieridae (Papilionoidea), Riodinidae (Papilionoidea), Lasiocampidae (Lasiocampoidea), Mimallonidae (Mimallonoidea), Erebidae (Noctuoidea), Noctuidae (Noctuoidea), Crambidae (Pyraloidea) and Limacodidae (Zygaenoidea) (Figs. 2, S12, Table S1). These families represent potential subjects for future comparative studies when more complete phylogenies become available, and if eyespots are found in more than a handful of species.

To obtain a preliminary picture of whether eyespots are widespread within these clades, we searched image databases for close relatives of eyespot-bearing taxa within these families and recorded presence or absence of eyespots not included in our Ditrysia phylogeny. We looked at images available of up to three species within these families and found that 59% of the species with eyespots represented in Fig. 2 have congeners (species with which they share a genus) also with eyespots. Of the remaining species for which we did not find congeners with eyespots, 32% were part of a subfamily and 3% part of a family with eyespot-bearing species and 6% of species (three taxa) were not found to have a congener, subfamily, or family member with eyespots. This could be due to lack of image data (*Pterodecta felderi* and *Helicomitra pulchra*, in our phylogeny) or due to these clades being monotypic to the family level (*Carthaea saturnioides*). A visual summary of the species found on our tree with eyespots and up to three close relatives also supporting eyespots is provided in Fig. S12 and Table S1.

## Discussion

Our results show that eyespot evolution in Lepidoptera is diverse and complex. Most eyespot research up to now has been conducted in a single clade (Nymphalidae), but here we infer that eyespots have evolved multiple times independently across the Lepidoptera, including in many poorly known moth clades (Figs. 2-4). Our stochastic character mapping analysis inferred that the median number of Discal eyespot origins on our tree is 32 and the median number of Marginal eyespot origins is 21 (Table 1). Most of the clades with eyespots on our phylogeny have low taxonomic coverage, but we documented eyespots in close relatives of each of these lineages (Fig. S12, Table S1), suggesting the possibility of more ancestral origins than those inferred by our present analysis (Fig. 4). Future phylogenetic studies, with a denser taxonomic sampling, will be required to characterize the evolutionary history of eyespots in these lesser-known clades. Nonetheless, our study strongly suggests that eyespots are not homologous across the order Lepidoptera, i.e., the MRCA of all Ditrysia with eyespots did not have eyespots. We find that this result is robust to alternative analytical approaches as both ancestral state reconstructions (from tree-wide eyespot evolution models), and comparisons of different root-state assumptions within subclades, point at independent eyespot origins in the two main clades where these traits are known to be widely distributed, the brush-footed butterflies (Nymphalidae) and the giant silkmoths moths (Saturniidae).

Here we provide the first evidence that Discal eyespots are homologous, at least within the tribe Saturniini (Table 2) and may even predate the diversification of the clade (Figs. 2, S3, S6). Ancestral state reconstructions suggest that this homology might extend to the MRCA of Saturniidae (Figs. S3, S6). Our results highlight the need for future research with greater taxonomic resolution within the Saturniidae to more precisely infer the timing and origin of Discal eyespots (D’Abrera, 1995; 1998; 2012). Nonetheless, Discal eyespot homology is supported within Saturniini, despite extensive diversity in eyespot complexity, colour, and size (Fig. 5A, S3, S6, Table 2). In contrast, our model-comparison approach for homology inference of Discal eyespots in Nymphalidae was inconclusive (Table 2), but ancestral state reconstructions suggest two or more independent origins are plausible within the same clade in which Marginal eyespots are ancestral (Fig. S3, S6). Additional taxonomic sampling will be required to draw robust inferences on the possible ancestral origins of Discal eyespots in Nymphalidae, where the evolutionary history of Discal eyespots has not been previously investigated.

Our results are consistent with Marginal eyespots being homologous within the Nymphalidae but not so in the Saturniidae. The evolution of Marginal eyespots in the Nymphalidae has been studied before and we report similar findings to these previous studies. We report a single origin for these eyespots that aligns with the ‘late’ model proposed by Oliver et al. (2014). The ‘early’ and ‘late’ models (Oliver et al. 2012; 2014) differed in pinpointing the clade where eyespots first originated, due to a different way of scoring eyespots on the wing. In the ‘early’ model, eyespots were scored as being present or absent anywhere on the wing, whereas in the ‘late’ model eyespots were scored for specific wing sectors. In our analyses (Figs. 4, 5, Table 2) and in the ‘late’ model (Oliver et al. 2014), the origin of nymphalid eyespots is inferred to have occurred after the divergences between Danainae and most other nymphalid subfamilies. The current study, as well as these previous studies, agree on a single origin of Marginal eyespots within Nymphalidae. Marginal eyespots in the Saturniidae, however, are likely not all homologous, as ancestral state reconstructions suggested multiple independent origins of Marginal eyespots within this clade (Fig. S4). Marginal eyespots evolved and radiated within the Attacini tribe of the subfamily Saturniinae which includes species of the genus *Samia, Callosamia,* and *Hyalophora* (Table 2). Marginal eyespots likely evolved separately in species from the genus *Copaxa, Eupakardia* (monotypic), as well as *Epiphora* and *Therinia* (Fig. S1, S4, S9).

Despite the independent origin of eyespots in the Papilionoidea and Bombycoidea, two superfamilies currently understood to be ∼110 million years apart (Kawahara et al. 2019), it is still possible that these (and other) Lepidopteran eyespots share the same gene-regulatory network. Previous research by Murugesan et al. (2022) found that an appendage gene-regulatory network was co-opted to build Marginal eyespots in *Bicyclus anynana*, a nymphalid butterfly. It is possible that the same GRN was co-opted more than once across the Lepidoptera. In addition, if the appendage network is already deployed in one place in the wing, this may facilitate co-option to another place in the wing. Such idea was somewhat supported in our data as Marginal eyespots appear to have facilitated a more recent evolution of Discal eyespots in Nymphalidae (Fig. 5B, S10, S11). To test these co-option hypotheses, it will be important to molecularly characterize Marginal and Discal eyespot GRNs in moth lineages as well as Discal eyespots in nymphalid butterflies. Early immunochemistry work in two saturniid species detected the presence of two eyespot marker proteins previously identified in nymphalids, Distal-less and Engrailed (Monteiro et al. 2006).

Stronger evidence for the use of the same appendage GRN in these Discal moth eyespots may need to come from CRISPR knockouts of cis-regulatory elements belonging to common eyespot and appendage genes, showing that both traits are affected (Murugesan et al. 2022). Demonstrating that all eyespots use a single GRN, or, alternatively, that different “eyespot” GRNs can evolve using different genes and/or cis-regulatory elements is a highly interesting question for future research.

This line of research would help us understand molecular mechanisms of convergence of complex phenotypic traits.

The large number of independent origins for eyespots suggests that these traits can evolve readily, perhaps in a quantitative and gradual way. Previous genetic studies showed that eyespot number in *Bicyclus anynana* hindwings was regulated by a variety of quantitative trait loci (QTLs) around many different genes (Rivera-Colón et al. 2020). This suggests that eyespots differentiate following some threshold response to quantitative genetic variation. In addition, previous reaction-diffusion models also showed that eyespot centres could form on the wing by tweaks in a variety of different parameters (Nijhout, 1991; Connahs et al. 2019). It is possible that multiple genetic parameters facilitate the stable expression of *Distal-less* in the middle of the wing sector, an essential regulator of eyespots (Connahs et al. 2019; Matsuoka and Monteiro 2022). Given that the expression of *Distal-less* is linked to the co-option of the appendage GRN (Murugesan et al. 2022), a stable *Distal-less* expression may represent a successful co-option of the appendage GRN to produce an eyespot.

It is also interesting to note that eyespots evolved only in the larger members of Ditrysia, and this may represent a developmental constrain, where the origin of eyespots may be linked to the evolution of body and/or wing size. This hypothesis, which we termed the ‘eyespots in large wings hypothesis’ is further strengthened by the occurrence of eyespots in large non Ditrysian primitive Lepidoptera, such as *Zelotypia stacyi,* a Hepialidae, which was not part of this study. This species has large Discal eyespots, as well as a large body and wing size (Grehan and Mielke, 2018).

Mechanistically, it is possible that large wing sectors are required for the reaction-diffusion mechanism to take place to differentiate eyespot centres. A future study focusing on wing sector area, body size and eyespot occurrence can be undertaken to test this idea.

Finally, little is known about the ecological roles of eyespots in the large variety of taxa found in the order Lepidoptera. While some data suggest the location of eyespots is indicative of roles in predator deterrence, predator avoidance, or in sexual signalling, this study pooled the sexes and the wing surfaces due to the lack of sex- or surface-specific data for many of the species examined. Future studies are needed to connect changes in a species’ ecology with the evolutionary fate of eyespots.

## Conclusions

Our results revealed multiple independent origins of eyespot wing patterns across Ditrysia (Fig. 4, Table 1), the clade encompassing ∼99% of all Lepidoptera species. Both Discal and Marginal eyespots (Fig. 1) occur frequently in the giant silkmoths (Saturniidae) and brush-footed butterflies (Nymphalidae) (Fig. 2), yet eyespot evolution appears to have followed distinct trajectories in each family. In Nymphalidae, Marginal eyespots arose earlier and were followed by one or a few origins of Discal eyespots (Fig. 5B). In Saturniidae, Discal eyespots likely preceded Marginal eyespots, or these arose independently across different lineages (Fig. 5A, Table 2). These contrasting evolutionary histories call for comparative research on the gene-regulatory networks underlying eyespot development across clades and between eyespot types. Beyond these two well-studied families, eyespots occur widely across numerous lesser-known ditrysian lineages, underscoring the breadth of convergent evolution for eyespot patterns. Together, our findings highlight untapped opportunities to explore the ecological drivers and developmental mechanisms behind the repeated, independent origins of these complex and beautiful wing patterns across Lepidoptera.

## Materials and methods

### Molecular data

DNA sequence data for 645 species of Lepidoptera (moths and butterflies) and seven species of Trichoptera (caddisflies, outgroup) were kindly provided by Professor Emeritus Charles Mitter (University of Maryland, Table S2). Additional sequences for 70 species of Saturniidae (giant silkmoths) were downloaded from NCBI (GenBank). The full dataset was composed of protein-coding sequences from 27 genes (Table S3), with 5-24 fragments available across all species. Our taxonomic sampling includes ∼68% (90/133, Van Nieukerken et al. 2011) of all families and ∼65% (28/43, Van

Nieukerken et al. 2011) of all superfamilies in the order Lepidoptera. This study focuses on the suborder Ditrysia, which comprises ∼99% of currently described lepidopterans. Of these, ∼90% of families (90/100, Reiger et al. 2009) and ∼93-96% (28-30) of superfamilies (Van Nieukerken et al. 2011; Heikkilä et al. 2015; Mitter et al. 2017) are represented. For each superfamily, 1 – 128 species were sampled (median = 14). The full list of superfamilies as well as the number of species representatives for each superfamily are outlined in Table S2. We chose the Trichoptera for our outgroup because they are the closest extant relatives to the Lepidoptera (Mey et al. 2017), but distant enough to provide ingroup monophyly.

### Sequence alignment and phylogenetic inference

DNA sequences were aligned based on their amino acid translation using MACSE v 2.07 (Ranwez et al. 2011), and a maximum of 16 refinement iterations. The resulting alignment had a length of 22,608 bp and no premature stop codons or frameshift mutations.

Phylogenetic inference under maximum likelihood was conducted with IQ-TREE v 2.2.0 (Nguyen et al. 2015). We used ModelFinder (Kalyaanamoorthy et al. 2017) to select a codon-substitution model based on the concatenated gene matrix, followed by tree reconstruction using the best-fit model.

The best-fit model was the empirical codon model in Kosiol et al. (2007), with empirical codon frequencies and rate variation among sites following a gamma distribution containing 9 categories. Node support was estimated using ultra-fast bootstrap in UFBoot (Hoang et al. 2018), which converged under 200,000 iterations. UFBoot is an efficient approximation alternative to the traditional bootstrap method and it is particularly well-suited for large datasets such as ours.

After pruning the outgroup, we transformed the maximum likelihood tree for Ditrysia into a time tree, using the least square dating method in LSD2 v 2.4.4 (To et al. 2016). The LSD2 algorithm enables rapid divergence-time estimation, based on calibration points for ancestral and tip nodes. We lifted ancestral node calibrations from the most recent and complete phylogeny of Lepidoptera (Kawahara et al. 2019). This study used 16 fossils, 12 of which belong to taxa in Ditrysia, and were thus relevant for our analysis. We first identified the nodes in our maximum likelihood tree which corresponded to each calibrated node in Kawahara et al. (2019), excluding one fossil used to date a clade in Kawahara et al. (2019) that was not inferred as monophyletic in our tree. The ages of the remaining 11 fossils were then used as minimum ages for the LSD2 analysis on the rooted Ditrysia tree. The resulting time tree dated the common ancestor of Ditrysia at 152 Ma, consistent with previous estimates (Kawahara et al. 2019; Wahlberg et al. 2013).

### Image data collection

We scored images of each of the 715 species in our phylogeny for presence or absence of eyespot patterns (in all wing surfaces and in both sexes). Colour patterns were identified as eyespots if they met the following criterion: Patterns contained a circular or oblong spot encircled by one or more concentric rings of a distinct colour that also contrast with the overall background colour of the wing (Fig. 3). Eyespots were further classified into Marginal or Discal, based on their locations on the wing surface (Fig. 1). To explore the sensitivity of our results to the eyespot-defining criterion, we repeated our main comparative analyses under a more inclusive definition of eyespots, where non-circular/oblong shapes (e.g., triangles, rectangles, teardrops, Fig. S1) encircled by another ring of colour were also considered. This relaxed eyespot criterion resulted in an 22% increase in the prevalence of Discal eyespots (from 82 to 100 species), whereas the number of species with Marginal eyespots was unaffected by the eyespot definition (41 in both the strict and relaxed data sets).

We first queried a wide range of online databases for images of the type specimen of each of the sampled species (Table S4). If not available, images of other (non-type) specimens were collected from the same databases with a preference for museum specimens over other specimens. In cases where sequence data was assigned to a particular subspecies, we prioritized images of the same taxonomic rank when available. 85 species included in our dataset lacked publicly available and/or reliably identified images of any form. We photographed 81 of these species at the Mcguire Center for Lepidoptera (MGCL, Florida Museum of Natural History, 3215 Hull Rd, Gainesville, Fl 32611), using a Cannon D50 DSLR camera. We were unable to obtain images for 4 species (0.56%), which are treated as missing data in all comparative analyses. Wing surfaces were pooled together due to a lack of publicly available data for all wing surfaces in all species compromising the dataset.

### Modelling the origin and loss of eyespots in Lepidoptera

To model eyespot evolution across Lepidoptera, we used RevBayes v1.3.0 (Höhna et al. 2016). RevBayes is an open-source software designed for Bayesian phylogenetic inference. It allows users to build probabilistic graphical models using an interactive model-specification language. Eyespots were modelled as a discrete trait with two states (presence/absence). Eyespot states had unequal transition rates, drawn from identical exponential priors. The rate parameters of these priors were set to reflect an expectation of 10 events (10 eyespot gains and 10 eyespot losses) along the tree. Root state frequencies were in turn drawn from a Dirichlet prior, assuming equal probabilities of presence or absence of eyespots at the origin of the tree. Eyespot evolution was modelled separately using three datasets: eyespot presence/absence irrespective of eyespot location on the wing; presence/absence of Marginal eyespots; and presence/absence of Discal eyespots. For all three datasets, eyespots could be located on any wing surface (dorsal/ventral surfaces of fore or hindwings) and in either sex. For the Discal and all-eyespot datasets, we repeated the analyses under the ‘relaxed’ definition of eyespots. Our results are qualitatively equivalent for the two eyespot criteria, so we only report the results from analyses under the strict eyespot definition in the main text and provide inferences based on the relaxed eyespot definition in the SM (Figs. S3 - S6).

Each model was run for 100,000 iterations after an initial burn-in of 10,000, and tuning parameter proposals every 1,000 iterations on two independent chains. We evaluated the convergence and stationarity of the MCMC chains for each model using the R package *Convenience* v1.0.0 (Fanreti et al. 2013). Joint conditional ancestral states were sampled every 100 iterations and plotted using RevGadgets v1.1.1 (Tribble et al. 2023), *ggplot2* v3.4.3 (Wickham et al. 2016) and *ggtree* v3.9.0 (Yu et al. 2017; 2018, Yu, 2020; 2022; Xu et al. 2022). Stochastic character histories (Freyman and Höhna, 2019) were sampled every 100 iterations and summarized by computing the median number of transitions (eyespots gains and losses) and corresponding 95% highest posterior density (HPD) intervals. ITOL was used to enhance the aesthetics of phylogenetic figures (ITOL, 2023).

### Testing eyespot homology in selected clades

We next implemented a model-comparison approach to investigate eyespot homology among extant species of Lepidoptera. We identified all subclades including two or more taxa bearing eyespots, regardless of eyespot location. If the most recent common ancestor (MRCA) of these clades also displayed eyespots, it is likely that eyespots in extant taxa are homologous — i.e. inherited from their common ancestor. To test this hypothesis, we extracted all the above-mentioned subclades (Fig. S7 for schematic example) from the complete time tree, and for each subclade applied two alternative versions of the discrete-trait model described previously. In the first version (hereafter the ‘multiple origins’ model), we constrained the root state to eyespot absence, by assigning a fully informative prior to the root state parameter. In the second version (the ‘common ancestor’ model), we instead enforced a MRCA with eyespots by setting the root frequency of eyespot presence to one.

We subsequently computed the marginal likelihood of the two alternative models for each subclade and for our entire phylogeny, given the character data. A marginal likelihood indicates the fit of a model, averaged across its entire parameter space, to a data set (Höhna et al. 2016). Marginal likelihood estimation is routinely used in phylogenetic analysis, for instance, in the selection of appropriate nucleotide substitution models and for species delimitation applications (Oaks et al. 2019). Unlike other model comparison approaches, marginal likelihoods are weighted by the prior, as prior beliefs are bound to influence model fit (Xie et al. 2011). Here, the two models being compared (the ‘multiple origins’ model and the ‘common ancestor’ model) differ only by the prior controlling the character state at the root. Their direct comparison thus serves as a statistical test for the presence of eyespots in the most recent common ancestor (MRCA) of each subclade with eyespots. Marginal likelihood was estimated via the stepping stone algorithm (Fan et al. 2011; Xie et al. 2011). The power posterior analysis was split into 50 intervals between the prior and posterior and was run for 5,000 iterations with a burn-in of 5,000 generations. We repeated stepping-stone marginal-likelihood estimation to quantify estimation error across independent runs. Across clades, independent runs of the same analysis differed by a median log_10_BF of 0.030 (95th percentile = 0.106, max = 0.302).

Marginal likelihood estimates were then used to calculate the log10 of the raw Bayes Factor (log_10_BF) between the ‘multiple origins’ and ‘common ancestor’ models. Given convention (Jeffreys 1961) and the observed run-to-run variation in marginal-likelihood estimation, we interpret log_10_BF values between 0–0.5 as barely worth mentioning, values 0.5–1 as substantial evidence, values between 1–2 as strong evidence, and values > 2 as decisive evidence in favour of the ‘common ancestor’ model, if positive. Negative values were in turn interpreted as support for the ‘multiple origins’ model.

To validate this approach, we simulated trees and character histories, based on previous diversification rate estimates in Lepidoptera (Heikkilä et al. 2012), and the transition rate estimates from our main discrete-character model above (see *Modelling the origin and loss of eyespots in Lepidoptera*). On these simulated data, our model comparison approach had Type-I error rates < 0.025, indicating a low probability of incorrectly supporting the ‘common ancestor’ model. However, the expected log_10_BF values were also generally low, especially for small trees (100 terminal taxa: log_10_BF = 1.18; 50 terminal taxa: log_10_BF = 0.74; 10 terminal taxa: log_10_BF = 0.19), suggesting that the model comparison approach is conservative and even when eyespots are indeed ancestral, support for such inference will likely be modest (Fig S13). See *Inference of ancestral eyespot origins: validating the model-comparison approach* in the Supporting Material for detailed simulation results.

### Correlated evolution of Marginal and Discal eyespots

We then focused on two clades (Papilionoidea and Bombycoidea), in which both Marginal and Discal eyespots have evolved in distinctly ordered sequences (see Results). These findings suggested that an ancestral eyespot origin on one wing region (Marginal or Discal) could facilitate the subsequent evolution of eyespots on the alternative region, by redeploying the ancestral eyespot gene-regulatory network on a novel tissue domain. We tested this hypothesis using a correlated evolution analysis for each clade, where the probability of gaining Marginal or Discal eyespots can depend on whether the alternative eyespot type is already present. To do this, we modelled a reversible jump mixture distribution, which samples the posterior probabilities of parameter spaces for correlated and independent eyespot evolution models. Support for the correlated evolution model in the posterior distribution of the reversible jump parameter would indicate that the evolution of one eyespot type depends on the presence of the other. If the correlated evolution model is supported, estimates of transition rates would indicate whether the pre-existence of an alternative eyespot type facilitates or hinder the evolution of eyespots on a novel wing domain. Alternatively, support for the independent model would suggest that Discal and Marginal are similarly likely to evolve whether the alternative eyespot type is present or absent.

As in our previous analyses, we modelled all transition rates (eyespots gains and losses on each wing region) as drawn from identical exponential priors in Revbayes v1.3.0 (Höhna et al. 2016). The rate parameter of the exponential priors corresponded to a mean of 10 transitions across the tree. Root frequencies were drawn from a flat Dirichlet distribution. The models were run for 10,000,000 iterations, with a pre-burn-in of 25% and tuning parameter proposals every 5,000 iterations on two independent chains. Joint conditional ancestral states were sampled every 10,000 iterations and plotted using *RevGadgets* v1.1.1 (Tribble et al. 2023) in R. We evaluated the convergence and stationarity of the MCMC chains for each model using the R package *Convenience* V1.0.0 (Fanreti et al. 2013).

### Identifying potential eyespot pattern common ancestors

Our initial image data collection included only species with available sequence data, resulting in several taxa (19 families in 11 superfamilies) with a single eyespot (Discal and/or Marginal) bearing taxon. To address gaps arising from our initial image dataset, which included only taxa for which sequence data were available and consequently yielded several families (19 families in 11 superfamilies) represented by a single species bearing Discal and/or Marginal eyespots. We evaluated whether these taxa belong to broader lineages characterized by eyespot presence. Specifically, for each family represented in our phylogeny with Discal or Marginal eyespot occurrences, we surveyed all currently recognized congeneric and con-familial species with online available image resources. Our objective was to determine whether close relatives of the taxa included in our tree (Fig. 2, Table S2) also exhibit Discal and/or Marginal eyespots, thereby informing whether these lineages plausibly share a deeper, common origin of eyespot patterning. Given the substantial variation among the 19 families (across 11 superfamilies) in both species’ richness and frequency of eyespot expression, we provide a synthesized overview of these observations in tabular and figure form, illustrating up to three representative species per family (Figs. S12, Table S1).

## Acknowledgements

We thank Dr Mark Willmott and Prof Akito Kawahara (MGCL) for hosting B.H. at the MGCL and facilitating access to the collections; Prof emeritus Charles Mitter (University of Maryland) for graciously sending us molecular sequence data for most specimens reanalysed here; Monteiro lab members for lively discussion. B.H was supported by a National Research Foundation (NRF) Graduate Fellowship; B.W. was funded by an International Postdoc Grant from the Swedish Research Council (VR; grant no. 2019-06444); and research was supported by a NRF Investigatorship award (NRF-NRFI05-2019-0006) to A.M.

