## Supplementary tables, figures and text for "Eyespots originated multiple times independently across the Lepidoptera"

### Supporting Material for: Eyespots originated multiple times independently across the Lepidoptera

A

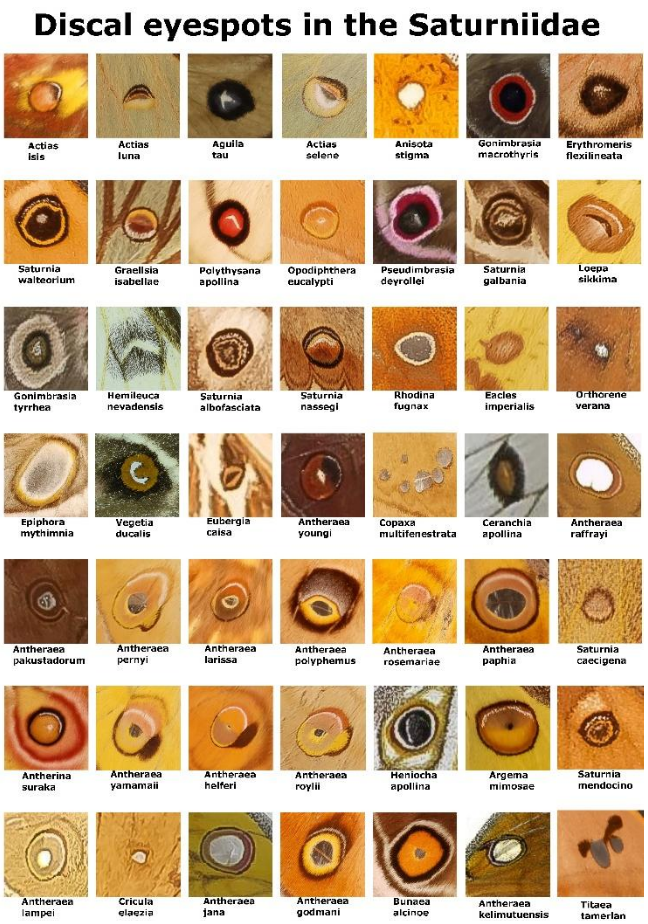


B

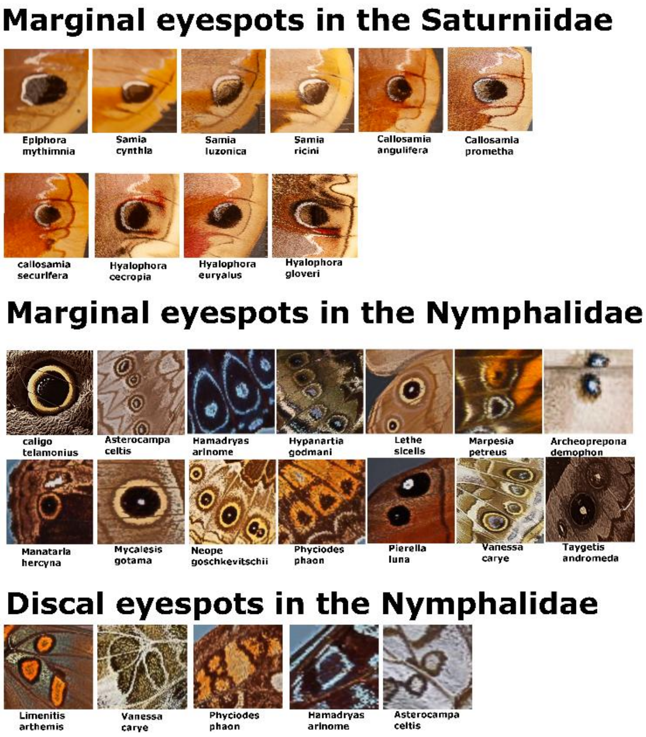


C

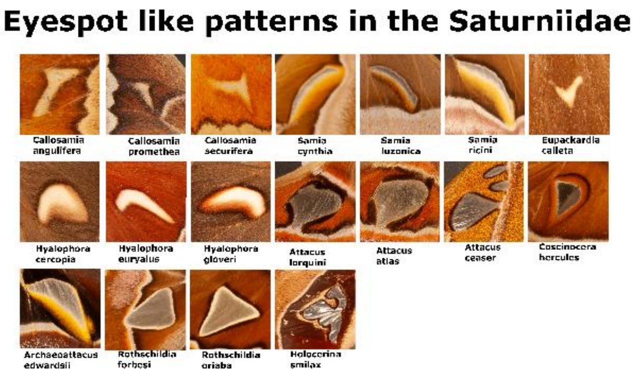


**Figure S1. Photographs of individual eyespot occurrences in species included in our phylogenetic analysis.** A) Discal eyespot patterns of all the Saturniidae included in our phylogeny only. B) Marginal eyespot patterns of all Saturniidae and Nymphalidae; Discal eyespots of all Nymphalidae included in our phylogeny only. C) Eyespot-like patterns (Observed in the Saturniidae only) included in the analysis when the circular criterion for eyespot shape was relaxed to include non-circular/oblong shapes (Fig. S2).

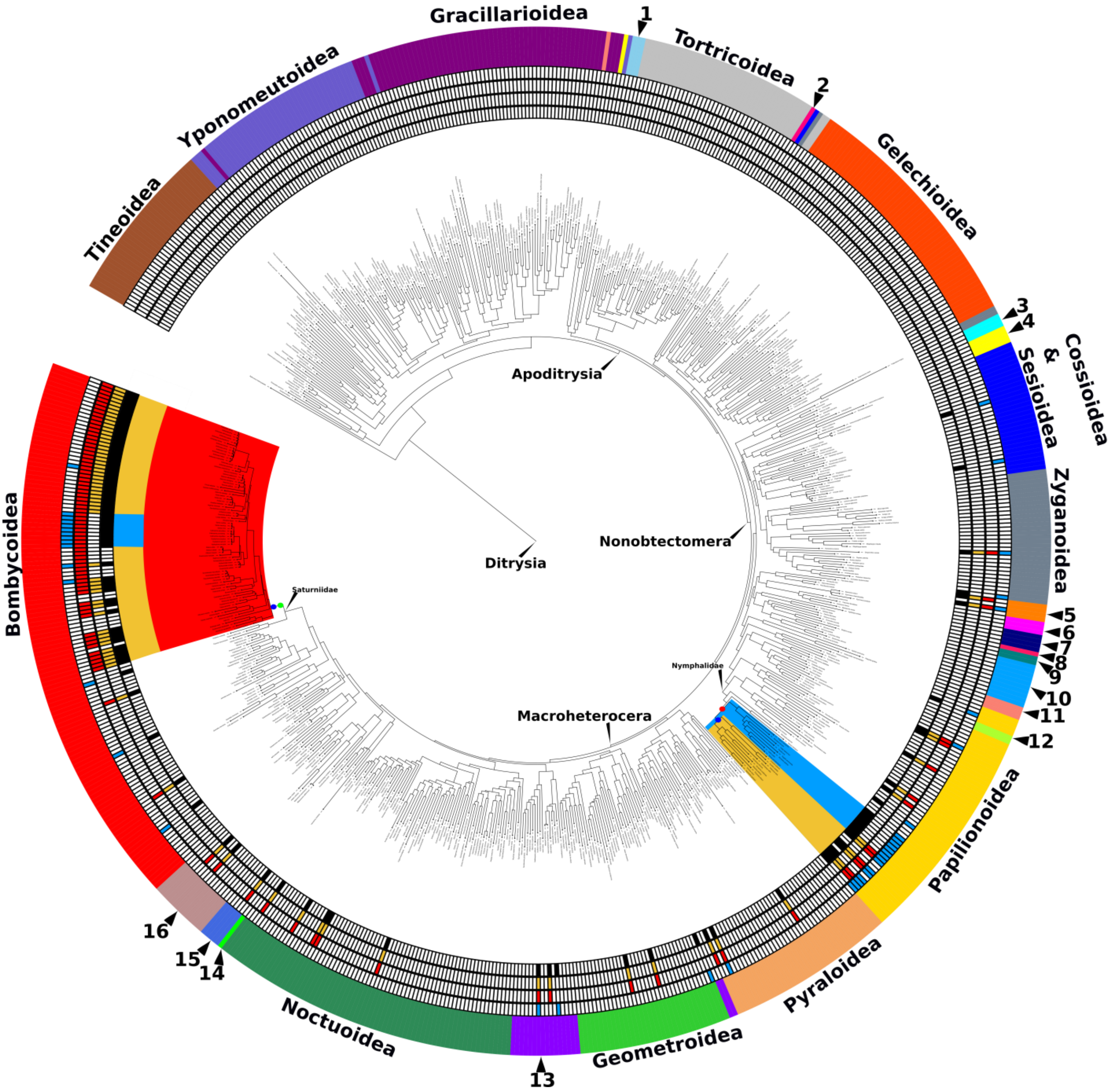


**Figure S2. Phylogeny of the Lepidoptera compromising 715 species from 28 superfamilies.** The superfamilies are outlined on the outer circumference of the tree. Superfamilies which are too small to be written are numbered as the following: 1 = Urodoidea, 2 = Galacticoidea, 3 = Immoidea, 4 = Choreutoidea, 5 = Calliduloidea, 6= Epermenoidea, 7 = Carposinoidea, 8 = Copomorphoidea, 9 = Hyblaeoidea, 10 = Thyridoidea, 11 = Pterophoroidea, 12 = Hedyloidea, 13 = Drepanoidea, 14 = Cimelioidea, 15 = Mimallonoidea, 16 = Lasiocampoidea. Eyespots are highlighted in blue (Marginal) and yellow (Discal). The presence or absence of eyespots for each individual species is marked by blue (Marginal) or yellow (Discal) square, as well as a black square indicating the presence of the trait. Taxa which display Discal eyespots under the relaxed criteria for eyespot definition (see Methods) are marked with red squares. Ancestral state reconstruction of Discal eyespots based on these more inclusive criteria does not support a more ancestral origin of Discal eyespots, marked with a green circle and with descendants highlighted in red. This putative ancestral origin was weakly but positively supported in our model-comparison approach (see Methods; $\log_{10}$ BF under stepping stone marginal likelihood estimation = 0.145). Additional $\log_{10}$ BF values for the all relaxed data are also available in Table S5.

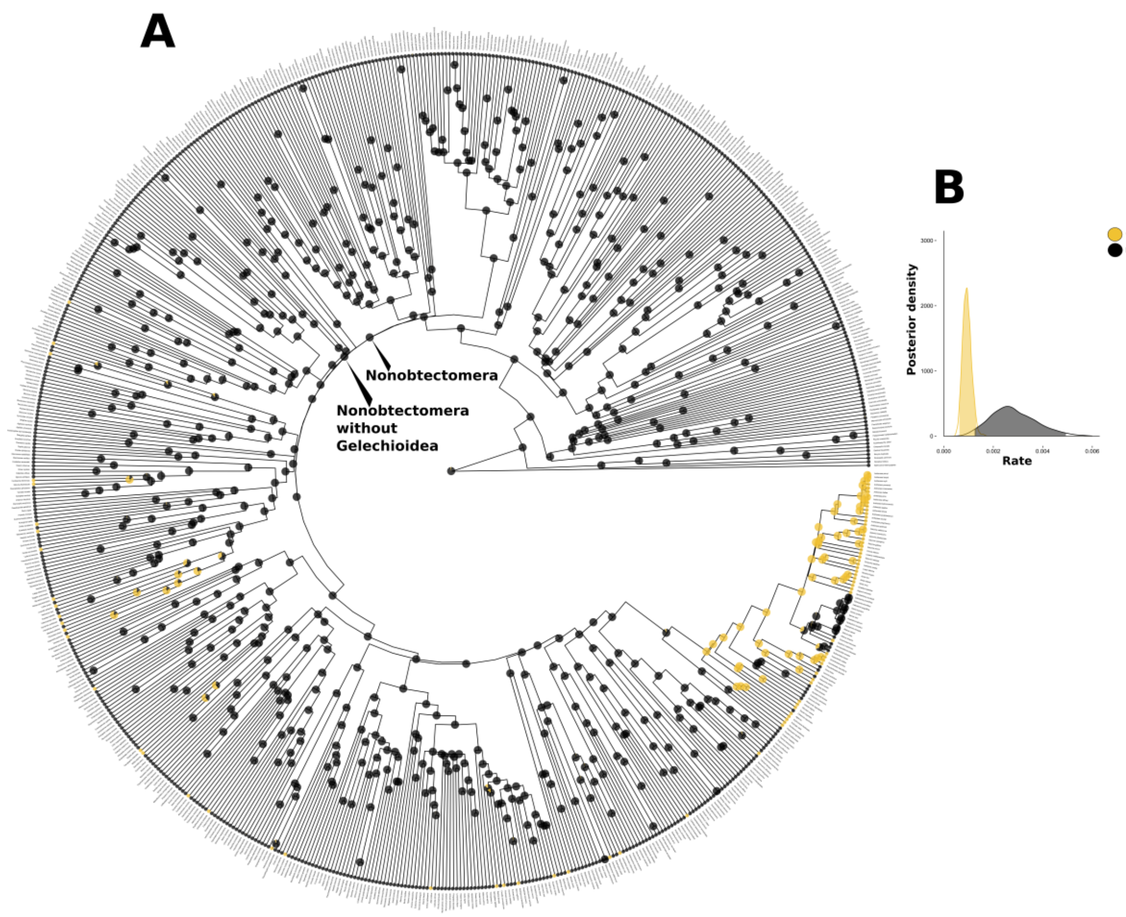


**Figure S3. Evolution of Discal eyespots in Lepidoptera.** We modelled Discal eyespot evolution as a discrete trait with two alternative states (presence or absence). **A)** Phylogeny of the Lepidoptera with ancestral state reconstructions. Pies indicate the posterior probability of Discal eyespot presence for each internal node. The posterior probability (pp) of Discal eyespot presence at the root of Lepidoptera was 0.032. The pp of a discal eyespot at the base of the Nonobtectomera was 0. The pp of a discal eyespot within the nonobtectomera excluding the Gelechioidea but including all present-day discal eyespot species was 0. **B)** Posterior densities for the rates of discal eyespot gain and loss across Lepidoptera.

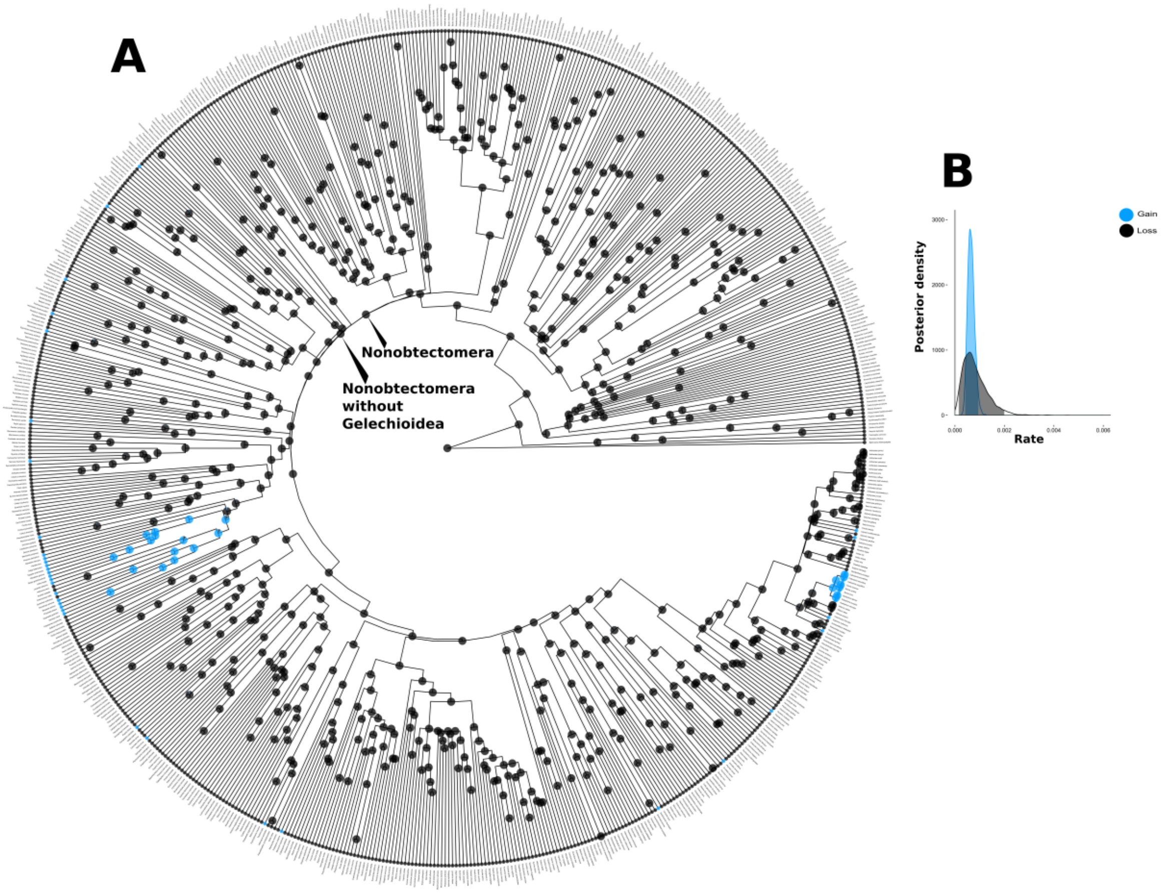


**Figure S4. Evolution of Marginal eyespots in Lepidoptera.** We modelled Marginal eyespot evolution as a discrete trait with two alternative states (presence or absence). **A)** Phylogeny of the Lepidoptera with ancestral state reconstructions. Pies indicate the posterior probability of Marginal eyespot presence for each internal node. The posterior probability (pp) of Marginal eyespot presence at the root of Lepidoptera was 0.0025. The pp of a marginal eyespot at the base of the Nonobtectomera was 0 and finally the pp of a Marginal eyespot within the nonobtectomera excluding the Gelechioidea but including all present-day Marginal eyespot species is 0. **B)** Posterior densities for the rates of marginal eyespot gain and loss across Lepidoptera.

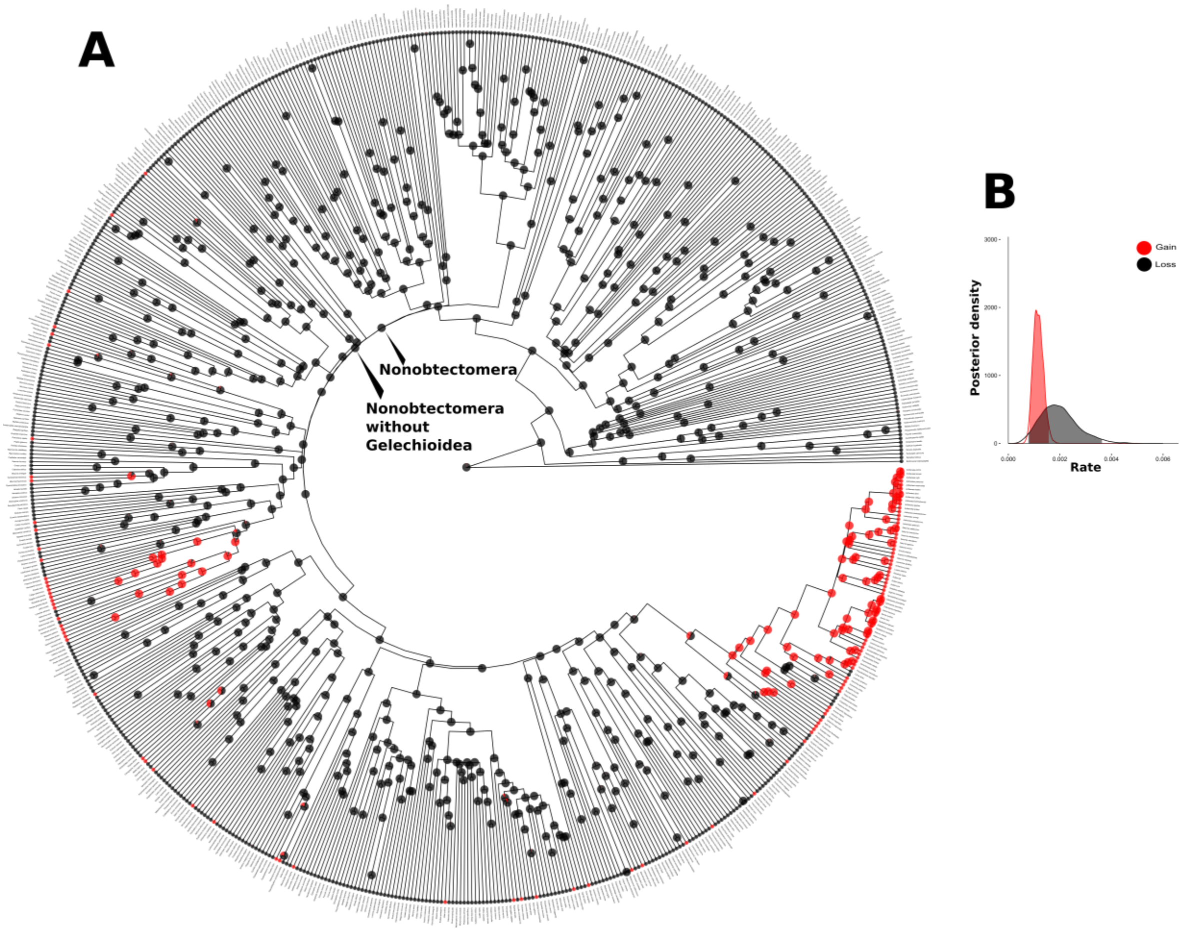


**Figure S5. Eyespot ancestral state reconstruction and transition rates in Lepidoptera for Marginal and Discal eyespots combined (relaxed).** We modelled eyespot evolution (all relaxed) as a discrete trait with two alternative states (presence or absence) for eyespots regardless of location (Discal or Marginal). **A)** Phylogeny of the Lepidoptera with ancestral state reconstructions. Pies indicate the posterior probability of eyespot (relaxed) presence for each internal node. The posterior probability (pp) of eyespot (relaxed) presence at the root of Lepidoptera was 0.019351. The pp of an eyespot (relaxed) at the base of the Nonobtectomera was 0 and finally the pp of an eyespot (relaxed) within the nonobtectomera excluding the Gelechioidea but including all present-day species with eyespots is 0. **B)** Posterior densities for the rates of eyespot gain and loss across Lepidoptera.

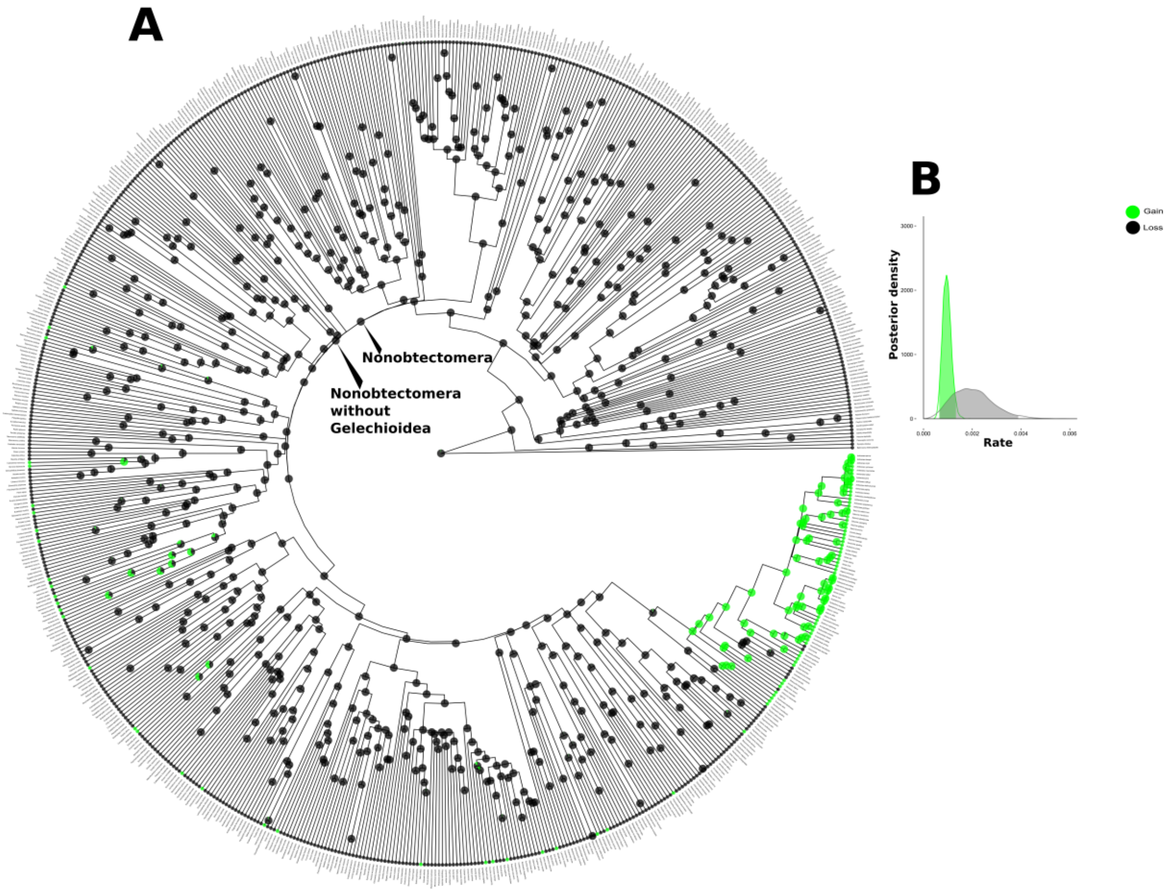


**Figure S6. Evolution of Discal (relaxed) eyespots in Lepidoptera.** We modelled Discal (relaxed) eyespot evolution as a discrete trait with two alternative states (presence or absence). **A)** Phylogeny of the Lepidoptera with ancestral state reconstructions. Pies indicate the posterior probability of Discal (relaxed) eyespot presence for each internal node. The posterior probability (pp) of Discal (relaxed) eyespot presence at the root of Lepidoptera was 0.020599. The pp of a Discal (relaxed) eyespot at the base of the Nonobtectomera was 0 and finally the pp of a Discal eyespot within the nonobtectomera excluding the Gelechioidea but including all present-day Discal eyespot species is 0. **B)** Posterior densities for the rates of Discal (relaxed) eyespot gain and loss across Lepidoptera.

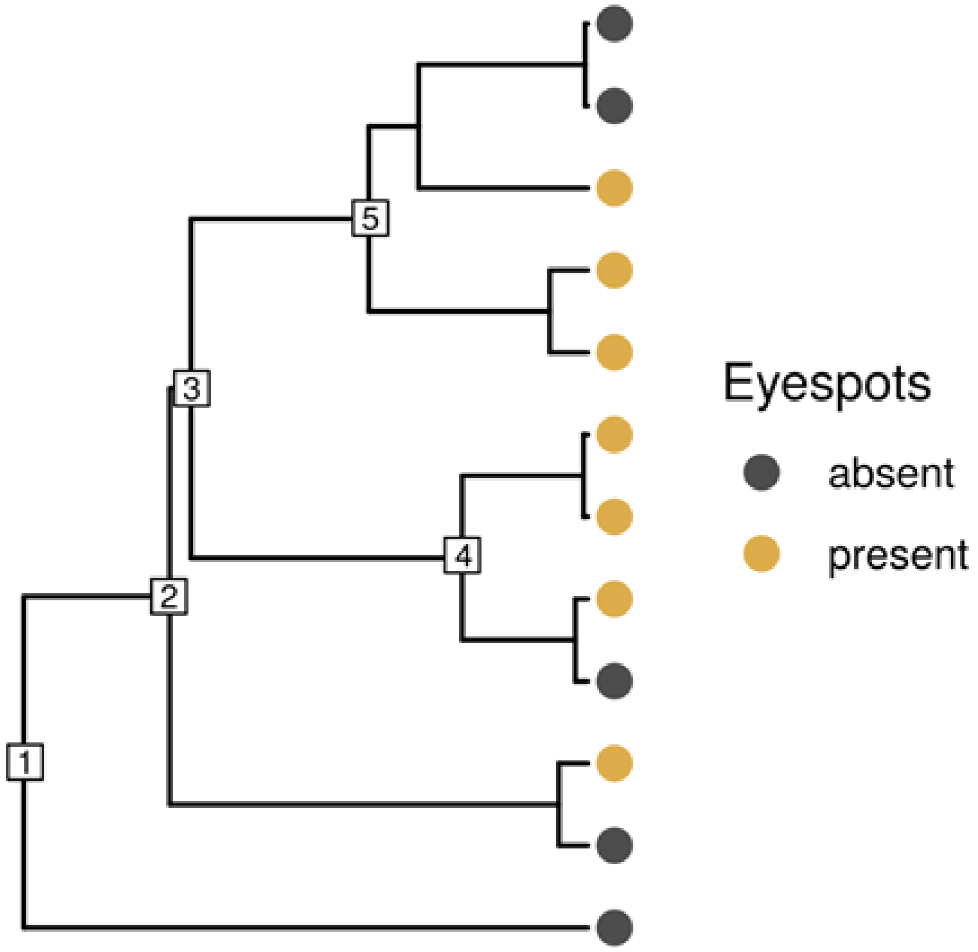


**Figure S7. Schematic representation of data partitioning for tests of eyespot homology**. We extracted all subclades with at least two eyespot-bearing taxa. In this schematic phylogeny, the root nodes of all such subclades are numbered from 2 to 5. For each of these subclades and for the entire phylogeny (with root node 1), we compared two alternative models with contrasting assumptions about the character state at the root. In the 'multiple origins' model we constrained the root node to eyespot absence and in the 'common ancestor' model we enforced eyespot presence at the root. We estimated the marginal likelihood of the two models and used Bayes factors to assess their relative fit to the data. Bayes factors thus indicated the strength of support for homology (i.e. a single ancestral evolutionary origin) or multiple independent origins of eyespots in the entire tree (1) and each relevant subclade (2-5). Subclades in which a common ancestral eyespot origin is supported are reported in Table 2.

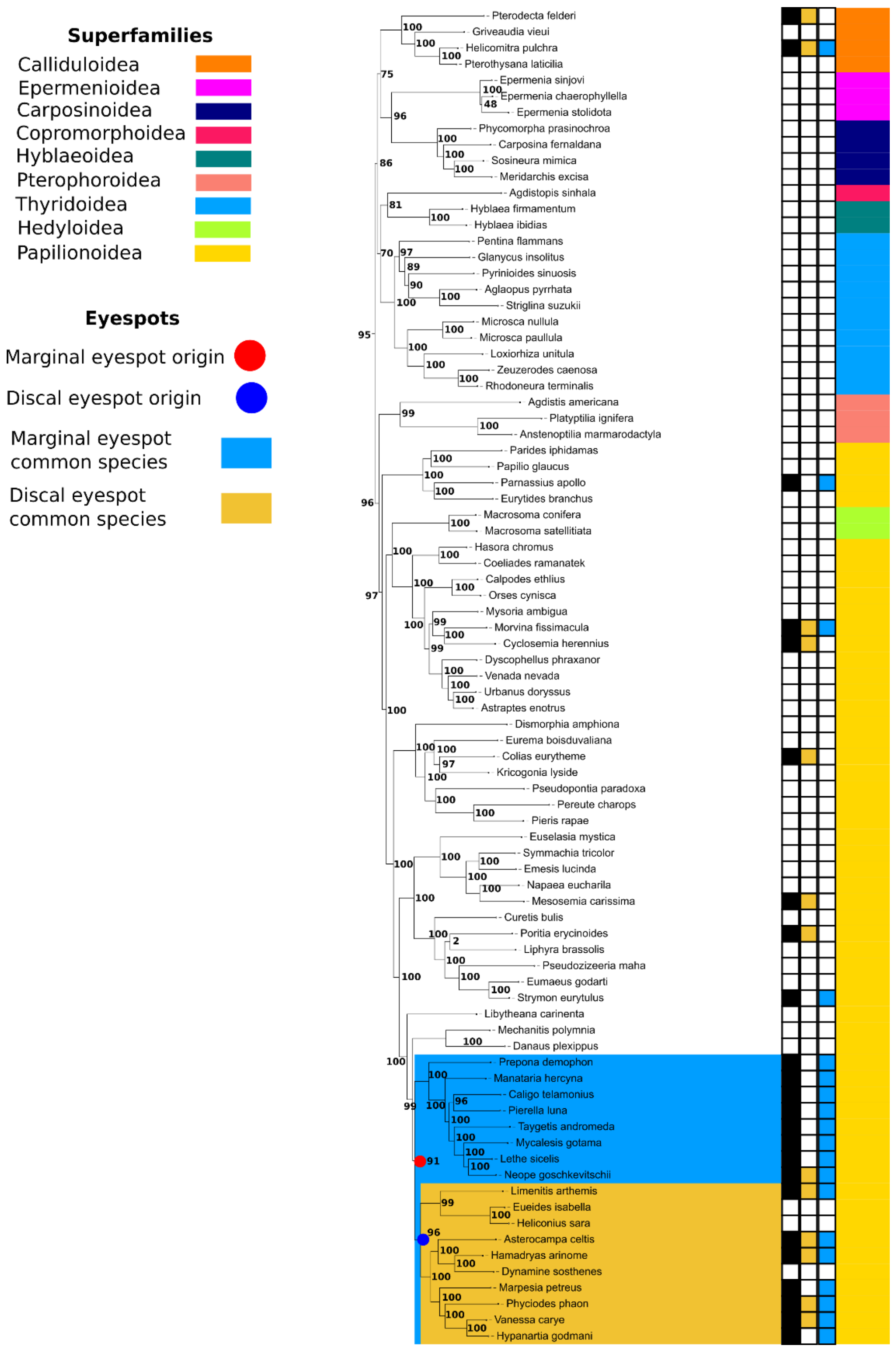


**Figure S8. View of the clade containing the Papilionoidea and related super families.** Marginal eyespot (red dot, blue square) and Discal eyespot origins (Blue dot, yellow square) origins calculated from our ancestral state reconstruction analysis as well as eyespot presence and absence are also visible.

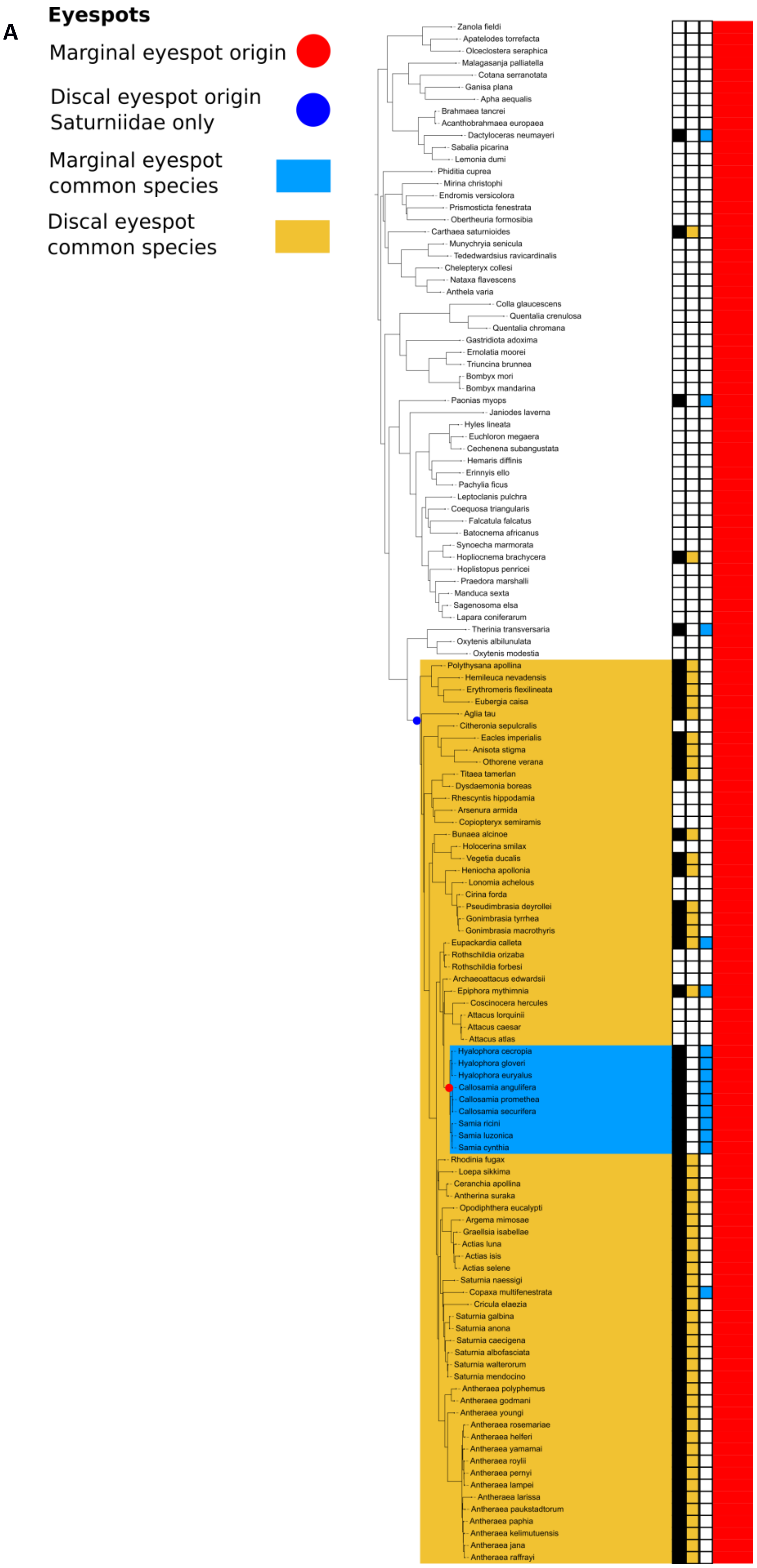
A
Eyespots
Marginal eyespot origin
Discal eyespot origin Saturniidae only
Marginal eyespot common species
Discal eyespot common species
Zanola fieldi
Apatelodes torrefacta
Olceclostera seraphica
Malagasanja palliatella
Cotana serranotata
Ganisa plana
Apha aequalis
Brahmaea tancrei
Acanthobrahmaea europaea
Dactyloceras neumayeri
Sabalia picarina
Lemonia dumi
Phiditia cuprea
Mirina christophi
Endromis versicolora
Prismosticta fenestrata
Obertheuria formosibia
Carthaea saturnioides
Munychryia senicula
Tededwardsius ravicardinalis
Chelepteryx collesi
Nataxa flavescens
Anthela varia
Colla glaucescens
Quentalia crenulosa
Quentalia chromana
Gastridiota adoxima
Ernolatia moorei
Triuncina brunnea
Bombyx mori
Bombyx mandarina
Paonias myops
Janiodes laverna
Hyles lineata
Euchloron megaera
Cechenena subangustata
Hemaris diffinis
Erinnyis ello
Pachylia ficus
Leptoclanis pulchra
Coequosa triangularis
Falcatula falcatus
Batocnema africanus
Synoecha marmorata
Hopliocnema brachycera
Hoplistopus penricei
Praedora marshalli
Manduca sexta
Sagenosoma elsa
Lapara coniferarum
Therinia transversaria
Oxytenis albilunulata
Oxytenis modestia
Polythysana apollina
Hemileuca nevadensis
Erythromeris flexilineata
Eubergia caisa
Aglia tau
Citheronia sepulcralis
Eacles imperialis
Anisota stigma
Othorene verana
Titaea tamerlan
Dysdaemonia boreas
Rhescyntis hippodamia
Arsenura armida
Copiopteryx semiramis
Bunaea alcinoe
Holocerina smilax
Vegetia ducalis
Heniocha apollonia
Lonomia achelous
Cirina forda
Pseudimbrasia deyrollei
Gonimbrasia tyrrhea
Gonimbrasia macrothyris
Eupackardia calleta
Rothschildia orizaba
Rothschildia forbesi
Archaeoattacus edwardsii
Epiphora mythimnia
Coscinocera hercules
Attacus lorquinii
Attacus caesar
Attacus atlas
Hyalophora cecropia
Hyalophora gloveri
Hyalophora euryalus
Callosamia angulifera
Callosamia promethea
Callosamia securifera
Samia ricini
Samia luzonica
Samia cynthia
Rhodinia fugax
Loepa sikkima
Ceranchia apollina
Antherina suraka
Opodiphthera eucalypti
Argema mimosae
Graellsia isabellae
Actias luna
Actias isis
Actias selene
Saturnia naessigi
Copaxa multifenestrata
Cricula elaezia
Saturnia galbina
Saturnia anona
Saturnia caecigena
Saturnia albofasciata
Saturnia walterorum
Saturnia mendocino
Antheraea polyphemus
Antheraea godmani
Antheraea youngi
Antheraea rosemariae
Antheraea helferi
Antheraea yamamai
Antheraea roylii
Antheraea pernyi
Antheraea lampei
Antheraea larissa
Antheraea paukstadtorum
Antheraea paphia
Antheraea kelimutuensis
Antheraea jana
Antheraea raffrayi

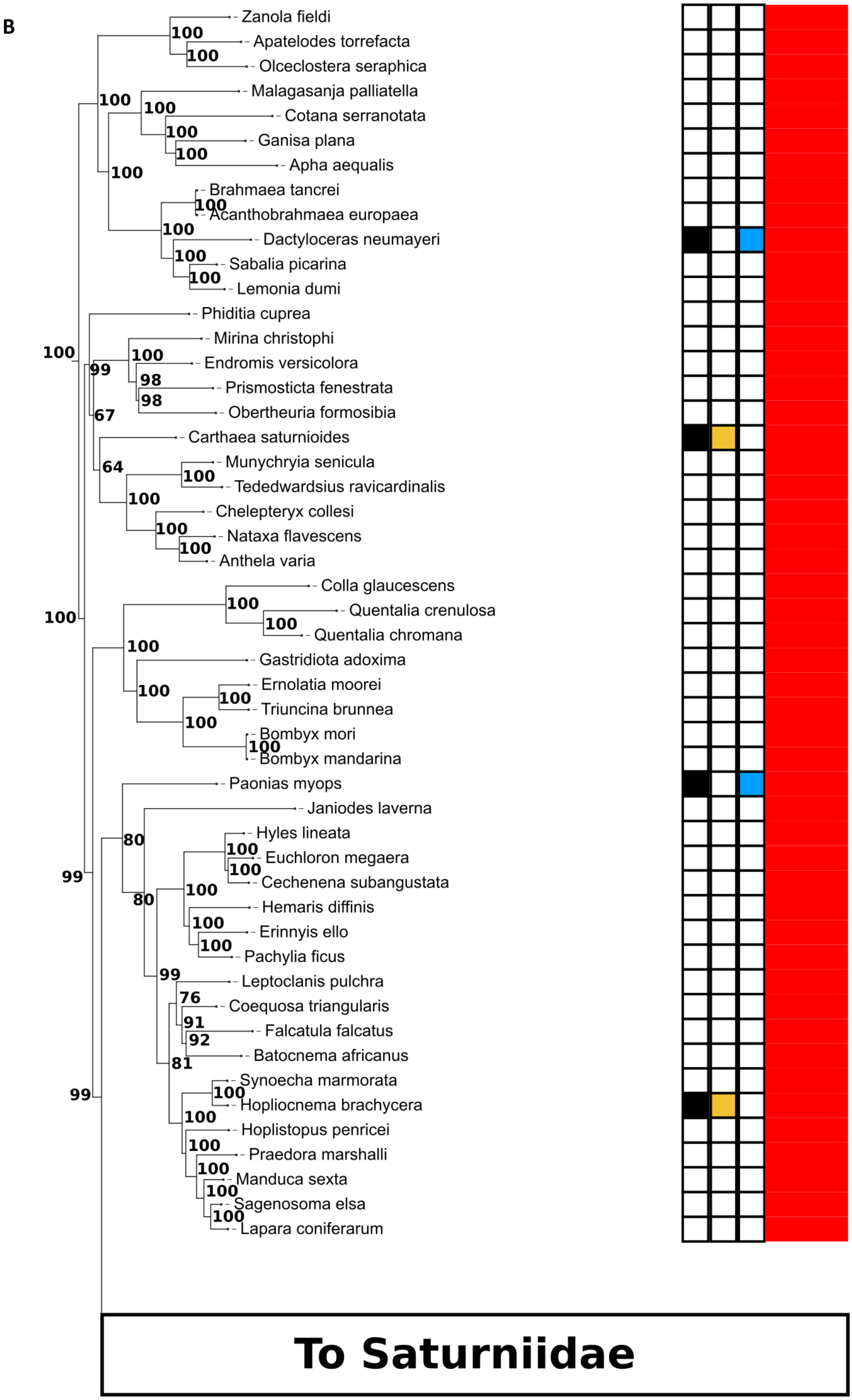

B
Zanola fieldi
Apatelodes torrefacta
Olceclostera seraphica
Malagasanja palliatella
Cotana serranotata
Ganisa plana
Apha aequalis
Brahmaea tancrei
Acanthobrahmaea europaea
Dactyloceras neumayeri
Sabalia picarina
Lemonia dumi
Phiditia cuprea
Mirina christophi
Endromis versicolora
Prismosticta fenestrata
Obertheuria formosibia
Carthaea saturnioides
Munychryia senicula
Tededwardsius ravicardinalis
Chelepteryx collesi
Nataxa flavescens
Anthela varia
Colla glaucescens
Quentalia crenulosa
Quentalia chromana
Gastridiota adoxima
Ernolatia moorei
Triuncina brunnea
Bombyx mori
Bombyx mandarina
Paonias myops
Janiodes laverna
Hyles lineata
Euchloron megaera
Cechenena subangustata
Hemaris diffinis
Erinnyis ello
Pachylia ficus
Leptoclanis pulchra
Coequosa triangularis
Falcatula falcatus
Batocnema africanus
Synoecha marmorata
Hopliocnema brachycera
Hoplistopus penricei
Praedora marshalli
Manduca sexta
Sagenosoma elsa
Lapara coniferarum
To Saturniidae

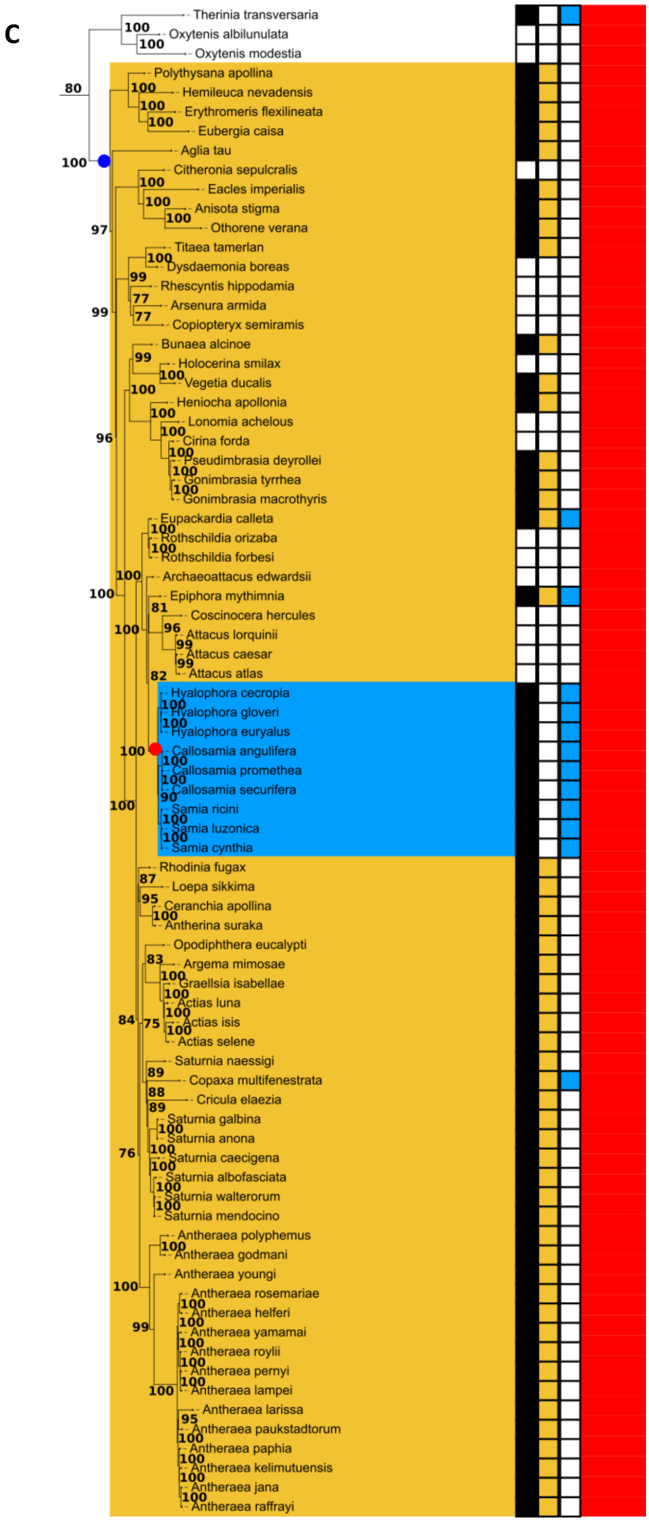


**Figure S9. Phylogenetic view of the Bombycoidea. A).** Clade containing the Bombycoidea indicating the presence of eyespot wing patterns. **B).** Subsample of the Bombycoidea showcasing the non Saturniid Bombycoidea and the presence of their eyespot patterns only. **C).** Subsample of the Bombycoidea focusing on the Saturniidae including the presence of eyespot wing patterns and their ancestral Discal eyespot (blue dot yellow square, Dark blue Triangle yellow square for Saturniidae + Sphingidae) and Marginal (red dot, blue square) eyespot origins.

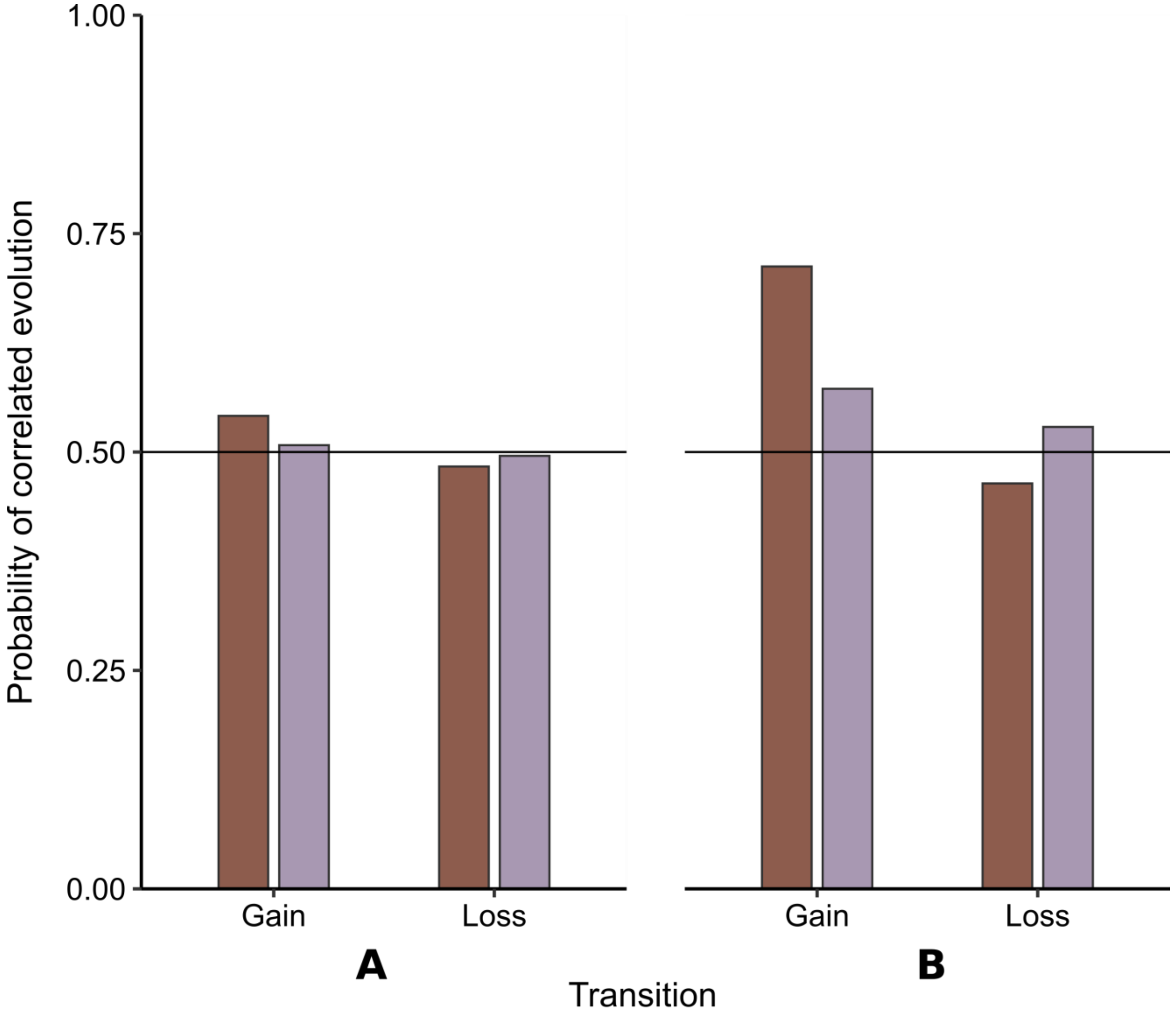


**Figure S10. Probability of correlated evolution between Discal and Marginal eyespots in Bombycoidea (A) and Papilionoidea (B).** Bar colours indicate the trait (focal eyespot type) involved in the transition: Discal eyespots are shown in maroon and Marginal eyespots in purple. Bars represent the probability that a character-state transition, an eyespot gain or loss, is conditional on the presence of the alternative eyespot type. For example, a high probability value for the gain of Discal eyespots would indicate that Discal eyespots are more likely to evolve in a lineage in which Marginal eyespots are already present. A low probability value would indicate that Discal eyespots evolve independently of Marginal eyespots, and a probability of gain of about 50% would indicate model uncertainty as to whether the origin of Discal eyespots is conditional on Marginal eyespots. Dashed lines indicate probability values corresponding to log10BF = 0.5, the threshold for substantial evidence in favour of either correlated or independent evolution of Discal and Marginal eyespots.

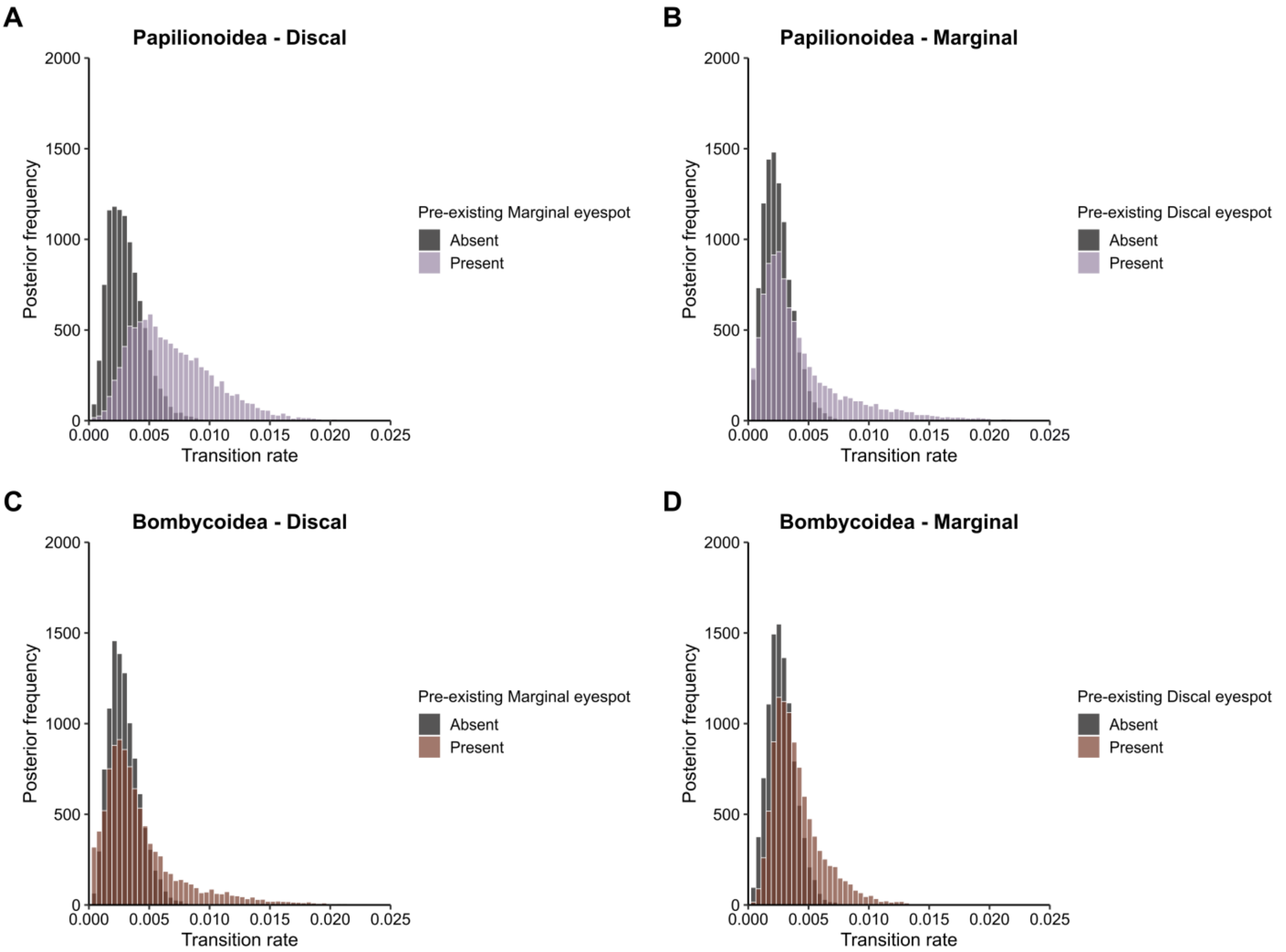


**Figure S11. Posterior distributions of the rates of eyespot gains conditional on the presence or absence of the alternative eyespot type**. (A) Gain of Discal eyespots conditional on Marginal eyespots in Papilionoidea. (B) Gain of Marginal eyespots conditional on Discal eyespots in Papilionoidea. (C) Gain of Discal eyespots conditional on Marginal eyespots in Bombycoidea. (D) Gain of Marginal eyespots conditional on Discal eyespots in Bombycoidea.

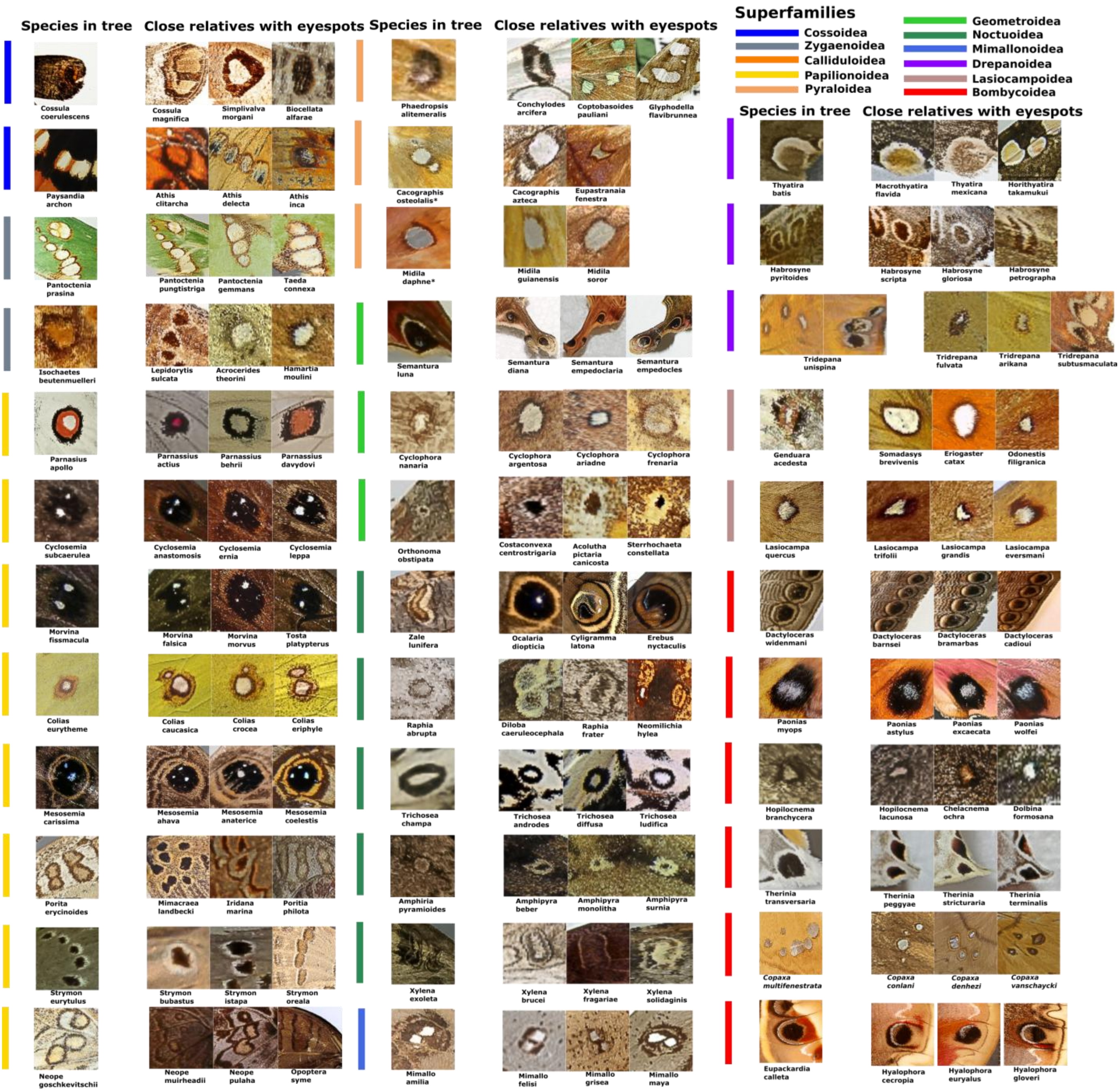


**Figure S12. Representation of individual eyespot occurrences in species not included in our phylogenetic analysis.** The super family Calliduloidea is absent because the two species with eyespots in this study *Pterodecta felderi* and *Helicomitra pulchra* have no close relatives which have photographic material available. One pair of species is marked with * (*Cacographis osteolalis* & *Midila daphne*) as they did not have three relatives available, only two relatives with eyespots are represented for each. *Carthaea saturnioides* is not represented because it is monotypic at the family level and for the purpose of this figure has no close relatives.

**Table S1.** List of species present on the tree and their close relatives (not on the tree) with eyespots. Taxa marked with a * have no images of living relatives available to determine whether an eyespot is present or absent. Taxa marked with a $ represent monotypic families and therefore have no living close relatives. Relatedness to eyespot-bearing species not included in the phylogeny is classified based on taxonomic ranks: F = family, SF = subfamily and G = genus. Type of eyespot key: M = Marginal, D = Discal and B = both types of eyespot present. Number of close relatives per level of relatedness = Genera 59%, subfamilies 32% and families, 3%.

| Superfamily | Species with eyespots in tree | Close relative 1 | Relatedness | Close relative 2 | Relatedness | Close relative 3 | Relatedness | Type of eyespot |
|---|---|---|---|---|---|---|---|---|
| Bombycoidea | *Dactyloceras neumayeri* | *Dactyloceras barnsi* | G | *Dactyloceras bramarbas* | G | *Dactyloceras cadioui* | G | M |
| Bombycoidea | *Carthaea saturnioides*$ | *N/A* | N/A | *N/A* | N/A | *N/A* | N/A | N/A |
| Bombycoidea | *Paonias myops* | *Paonias astylus* | G | *Paonias excaecata* | G | *Paonias wolfei* | G | M |
| Bombycoidea | *Hopliocnema brachycera* | *Hopilocnema lacunosa* | G | *Chelacnema ochra* | SF | *Dolbina formosana* | SF | D |
| Calliduloidea | *Pterodecta felderi** | *N/A* | N/A | *N/A* | N/A | *N/A* | N/A | N/A |
| Calliduloidea | *Helicomitra pulchra** | *N/A* | N/A | *N/A* | N/A | *N/A* | N/A | N/A |
| Cossoidea | *Cossula coerulescens* | *Cossula magnifica* | G | *Simplicivalva morgani* | SF | *Biocellata alfarae* | SF | M |
| Cossoidea | *Paysandisia archon* | *Athis clitarcha* | SF | *Athis delecta* | SF | *Athis inca* | SF | B |
| Drepanidae | *Tridrepana unispina* | *Tridrepana arikana* | G | *Tridrepana fulvata* | G | *Tridrepana subtusmaculata* | G | B |
| Drepanidae | *Habrosyne pyritoides* | *Habrosyne gloriosa* | G | *Habrosyne petrographa tapaishana* | G | *Habrosyne scripta* | G | D |
| Drepanidae | *Thyatira batis* | *Thyatira mexicana* | G | *Horithyatira takamukui* | SF | *Macrothyatira flavida* | SF | B |
| Geometroidea | *Orthonama obstipata* | *Acolutha pictaria canicosta* | SF | *Sterrhochaeta constellata* | SF | *Costaconvexa centrostrigaria* | SF | D |
| Geometroidea | *Cyclophora nanaria* | *Cyclophora argentosa* | G | *Cyclophora ariadne* | G | *Cyclophora frenaria* | G | D |
| Geometroidea | *Mania lunus* | *Mania diana* | G | *Mania empedoclaria* | G | *Mania empedocles* | G | M |
| Papilonoidea | *Cyclosemia herennius subcaerulea* | *Cyclosemia anastomosis* | G | *Cyclosemia earina* | G | *Cyclosemia leppa* | G | D |
| Papilonoidea | *Morvina fissimacula* | *Morvina falisca* | G | *Morvina morvus* | G | *Tosta platypterus* | SF | B |
| Papilonoidea | *Poritia erycinoides* | *Poritia philota* | G | *Iridana marina* | SF | *Mimacraea landbecki* | SF | D |
| Papilonoidea | *Strymon eurytulus* | *Strymon bubastus* | G | *Strymon istapa* | G | *Strymon oreala* | G | B |

| | | | | | | | | |
|---|---|---|---|---|---|---|---|---|
| Papilonoidea | *Parnassius apollo* | *Parnassius behrii* | G | *Parnassius davydovi* | G | *Parnassius actius* | G | M |
| Papilonoidea | *Colias eurytheme* | *Colias eriphyle* | G | *Colias caucasica* | G | *Colias crocea* | G | D |
| Papilonoidea | *Mesosemia carissima* | *Mesosemia ahava* | G | *Mesosemia antaerice* | G | *Mesosemia coelestis* | G | M |
| Lasiocampoidea | *Lasiocampa quercus* | *Lasiocampa trifolii* | G | *Lasiocampa grandis* | G | *Lasiocampa eversmanni* | G | D |
| Lasiocampoidea | *Genduara acedesta* | *Eriogaster catax* | SF | *Odonestis filigranica* | SF | *Somadasys brevivenis* | SF | D |
| Minimalloidea | *Mimallo amilia* | *Mimallo felisi* | G | *Mimallo grisea* | G | *Mimallo maya* | G | D |
| Noctuoidea | *Zale lunifera* | *Ocalaria dioptica* | SF | *Cyligramma latona* | SF | *Erebus nyctaculis* | SF | D |
| Noctuoidea | *Amphipyra pyramidoides* | *Amphipyra monolitha* | G | *Amphipyra berbera* | G | *Amphipyra surnia* | G | D |
| Noctuoidea | *Raphia abrupta* | *Raphia frater* | G | *Diloba caeruleocephala* | SF | *Neomilichia hylea* | F | D |
| Noctuoidea | *Xylena exsoleta* | *Xylena brucei* | G | *Xylena fragariae* | G | *Xylena solidaginis* | G | D |
| Noctuoidea | *Trichosea champa* | *Trichosea androdes* | G | *Trichosea diffusa* | G | *Trichosea ludifica* | G | D |
| Pyraloidea | *Cacographis osteolalis* | *Cacographis azteca* | G | *Eupastranaia fenestra* | SF | *N/A* | N/A | D |
| Pyraloidea | *Midila daphne* | *Midila guianensis* | G | *Midila soror* | G | *N/A* | N/A | D |
| Pyraloidea | *Phaedropsis alitemeralis* | *Coptobasoides pauliani* | SF | *Conchylodes arcifera* | SF | *Glyphodella flavibrunnea* | SF | D |
| Zygaenoidea | *Pantoctenia prasina* | *Pantoctenia gemmans* | G | *Pantoctenia punctistriga* | G | *Taeda connexa* | SF | B |
| Zygaenoidea | *Isochaetes beutenmuelleri* | *Lepidorytis sulcata* | SF | *Acrocerides theorini* | F | *Hamartia moulini* | F | D |

**Table S2.** list of all species, accession numbers and code names (original data sheet available on request) as provided by Prof Charles Mitter. Additional species added with NCBI GenBank data are at the bottom of this list. Species (n = 4) lacking an image data point of any kind are marked with an * next to their name.

| **Trichoptera (Outgroup)** | **Superfamily** | **Family** | **Accession** | **Codename** |
|---|---|---|---|---|
| *Agapetus sp* | - | Glossosomatidae | JBA-08-0111 | Agus |
| *Hydropsyche sp* | - | Hydropsychidae | JBA-08-0108-1 | Hysy |
| *Wormaldia moesta* | - | Philopotamidae | CWM-94-0226 | Wmo2 |
| *Plectrocnemia conspersa* | - | Polycentropodidae | JBA-08-0112 | Pctb |
| *Rhyacophila dorsalis* | - | Rhyacophilidae | JBA-08-0110 | Rhmd |

| | | | | |
|---|---|---|---|---|
| *Sericostoma sp* | - | Sericostomatidae | JBA-08-0116-1 | Seic |
| **Ditrysian Lepidoptera** | **Superfamily** | **Family** | **Accession number** | **Codename** |
| *Anthela varia* | Bombycoidea | Anthelidae | PZ028772 | Avar |
| *Tededwardsius ravicardinalis* | Bombycoidea | Anthelidae | AZ-07-3212 | Tedr |
| *Apatelodes torrefacta* | Bombycoidea | Bombycidae | CWM-96-0560 | Ator |
| *Olceclostera seraphica* | Bombycoidea | Bombycidae | RSP-94-1023 | Osera |
| *Zanola fieldi* | Bombycoidea | Bombycidae | 05-srnp-24117 | Zfid |
| *Bombyx mori* | Bombycoidea | Bombycidae | UNK-90-0062-3 | Bmor1 / 2 |
| *Quentalia chromana* | Bombycoidea | Bombycidae | DHJ-02-2407 | Quid |
| *Gastridiota adoxima* | Bombycoidea | Bombycidae | AZ-07-3202 | Gadx |
| *Triuncina brunnea* | Bombycoidea | Bombycidae | AYK-04-0514 | Trbr |
| *Bombyx mandarina* | Bombycoidea | Bombycidae | PZ028773 | Bmand |
| *Ernolatia moorei* | Bombycoidea | Bombycidae | PZ028774 | Emoor |
| *Colla glaucescens* | Bombycoidea | Bombycidae | PZ028775 | Cgla |
| *Quentalia crenulosa* | Bombycoidea | Bombycidae | PZ028776 | Qcren |
| *Obertheuria formosibia* | Bombycoidea | Bombycidae | AYK-04-0824-1 | Ofo |
| *Prismosticta fenestrata* | Bombycoidea | Bombycidae | AYK-04-0824-9 | Pfene |
| *Acanthobrahmaea europaea* | Bombycoidea | Brahmaeidae | RSP-95-0990 | Aeuro |
| *Dactyloceras neumayeri* | Bombycoidea | Brahmaeidae | AZ-07-3132 | Dawi |
| *Brahmaea tancrei* | Bombycoidea | Brahmaeidae | PZ028777 | Bhear |
| *Carthaea saturnioides* | Bombycoidea | Carthaeidae | TB-03-2177 | TB-03-2177 |
| *Endromis versicolora* | Bombycoidea | Endromidae | RSP-95-1008 | Eversicol |
| *Apha aequalis* | Bombycoidea | Eupterotidae | AYK-04-2506 | Aaeq |
| *Jana palliatella* | Bombycoidea | Eupterotidae | DCL-07-0005 | Jpta |
| *Cotana serranotata* | Bombycoidea | Eupterotidae | AZ-06-0059 | Cotan |
| *Ganisa plana* | Bombycoidea | Eupterotidae | AZ-07-3168 | Gpln |
| *Sabalia picarina* | Bombycoidea | Lemoniidae | AZ-06-0131 | Sabal |
| *Lemonia dumi* | Bombycoidea | Lemoniidae | RSP-96-0623 | Ldumi2 |
| *Mirina christophi* | Bombycoidea | Mirinidae | RXR-02-0511 | Mchr |
| *Phiditia cuprea* | Bombycoidea | Phiditidae | PZ028778 | Pcup |
| *Aglia tau* | Bombycoidea | Saturniidae | PZ028779 | Atau |
| *Arsenura armida* | Bombycoidea | Saturniidae | RWH-96-0876 | Aarm |
| *Janiodes laverna* | Bombycoidea | Saturniidae | KLW-03-2869 | Jcer |
| *Hemileuca nevadensis* | Bombycoidea | Saturniidae | PZ028780 | Hnev |
| *Oxytenis modestia* | Bombycoidea | Saturniidae | DHJ-02-2470 | Omod |
| *Therinia transversaria* | Bombycoidea | Saturniidae | 05-srnp-3527 | Atsv |

| | | | | |
|---|---|---|---|---|
| *Saturnia naessigi* | Bombycoidea | Saturniidae | NK-94-1026 | Snaes |
| *Antheraea paukstadtorum* | Bombycoidea | Saturniidae | PZ028781 | Apauk |
| *Antheraea polyphemus* | Bombycoidea | Saturniidae | UNK-97-3101 | Apol2 |
| *Hyles lineata* | Bombycoidea | Sphingidae | RSP-96-0929 | Hlin |
| *Cechenena subangustata* | Bombycoidea | Sphingidae | AYK-04-0214 | Ccsb1 |
| *Pachylia ficus* | Bombycoidea | Sphingidae | AYK-04-0335 | Chfc2 |
| *Euchloron megaera* | Bombycoidea | Sphingidae | IJK-03-3155 | Emga1 |
| *Erinnyis ello* | Bombycoidea | Sphingidae | RFD-96-0982 | Eys1 |
| *Hemaris diffinis* | Bombycoidea | Sphingidae | PZ028782 | Hthy |
| *Batocnema africanus* | Bombycoidea | Sphingidae | AZ-06-3209 | Baaf |
| *Paonias myops* | Bombycoidea | Sphingidae | MCN-03-1796 | Pmyo |
| *Coequosa triangularis* | Bombycoidea | Sphingidae | AZ-06-0211 | Coqu |
| *Falcatula falcatus* | Bombycoidea | Sphingidae | MF-06-0116 | Ffaa |
| *Hopliocnema brachycera* | Bombycoidea | Sphingidae | MJM-96-0232 | Hbr |
| *Leptoclanis pulchra* | Bombycoidea | Sphingidae | MF-05-0003 | Lplu |
| *Synoecha marmorata* | Bombycoidea | Sphingidae | AZ-06-0210 | Synm |
| *Manduca sexta* | Bombycoidea | Sphingidae | MM-03-2154 | Mqui |
| *Hoplistopus penricei* | Bombycoidea | Sphingidae | AZ-06-0207 | Hope |
| *Praedora marshalli* | Bombycoidea | Sphingidae | AZ-07-0208 | Prma |
| *Sagenosoma elsa* | Bombycoidea | Sphingidae | AYK-06-8000 | Sels |
| *Lapara coniferarum* | Bombycoidea | Sphingidae | PZ028783 | Lcon |
| *Pterodecta felderi* | Calliduloidea | Callidulidae | AYK 04 5717 | Ptfe |
| *Griveaudia vieui* | Calliduloidea | Callidulidae | DCL-07-0002 | Gvii |
| *Helicomitra pulchra* | Calliduloidea | Callidulidae | DCL-07-0003 | Hlpu |
| *Pterothysana laticilia* | Calliduloidea | Callidulidae | AZ-07-2027 | Pter |
| *Anthophila fabriciania* | Choreutoidea | Choreutidae | JR39-BE4 | Anfa |
| *Tebenna micalis* | Choreutoidea | Choreutidae | JCS-06-0122 | Tmic |
| *Saptha libanota* | Choreutoidea | Choreutidae | JR-08-002 | Sapth |
| *Millieria dolosalis* | Choreutoidea | Choreutidae | JR144-SL3 | Mido |
| *Anthophila felis* | Choreutoidea | Choreutidae | DRD-05-0218 | Hfel |
| *Axia margarita* | Cimelioidea | Cimeliidae | JBA-01-0051 | Axsp |
| *Carposina fernaldana* | Copromorphoidea | Carposinidae | JWB-08-0100 | Cafd |
| *Meridarchis excisa* | Copromorphoidea | Carposinidae | SWC-07-2002 | Merid |
| *Sosineura mimica* | Copromorphoidea | Carposinidae | MJM-96-0292 | Smim2 |
| *Phycomorpha prasinochroa* | Copromorphoidea | Copromorphidae | MM-08-7656 | Ppra |

| | | | | |
|---|---|---|---|---|
| *Culama crepera* | Cossoidea | Cossidae | MJM-96-0288 | Cucr |
| *Chilecomadia valdiviana* | Cossoidea | Cossidae | AYK-04-0538-02 | Cvdv |
| *Prionoxystus robiniae* | Cossoidea | Cossidae | CWM-94-0351 | Prob2 |
| *Cossula maruga* | Cossoidea | Cossidae | BL-08-0116 | Cosla |
| *Allostylus coerulescens* | Cossoidea | Cossidae | kn06-1104 | Allo |
| *Givira mucidus* | Cossoidea | Cossidae | AYK-06-7130 | Gmuc |
| *Langsdorfia lunifera* | Cossoidea | Cossidae | kn06-1132 | Llun |
| *Philiodoron frater* | Cossoidea | Cossidae | AYK-04-0542-12 | Pfoc |
| *Lebedodes cossula* | Cossoidea | Cossidae | SEM-06-2522 | Lebe |
| *Metarbela naumanni* | Cossoidea | Cossidae | WM-08-4004 | Mnau |
| *Archaeoses pentasema* | Cossoidea | Cossidae | AZ-07-2112 | Ahss |
| *Endoxyla encalypti* | Cossoidea | Cossidae | MJM-97-0302 | Eeny |
| *Psychogena personalis* | Cossoidea | Cossidae | RFD-96-0967B | Ppls |
| *Xyleutes mineus* | Cossoidea | Cossidae | AYK-04-1279-1 | Xmns |
| *Polyphagozerra coffeae* | Cossoidea | Cossidae | AYK-04-0786-2 | Zcof |
| *Cyclidia substigmaria modesta* | Drepanoidea | Drepanidae | AYK-04-0776-5 | Cysu |
| *Drepana arcuata* | Drepanoidea | Drepanidae | CWM-96-0579 | Darcu |
| *Macrauzata maxima* | Drepanoidea | Drepanidae | AYK 04 5709 | Mmax |
| *Macrocilix mysticata* | Drepanoidea | Drepanidae | AYK-04-0978-07 | Msys |
| *Oreta rosea* | Drepanoidea | Drepanidae | CWM-95-0466 | Oros |
| *Albara hollowayi* | Drepanoidea | Drepanidae | AYK-04-0824-17 | Ahwy |
| *Callidrepana palleola* | Drepanoidea | Drepanidae | AYK-06-7263 | Cpll |
| *Microblepsis accuminata* | Drepanoidea | Drepanidae | AYK-06-7266 | Miac2 |
| *Nordstromia grisearia* | Drepanoidea | Drepanidae | AYK-04-5232 | Ngri |
| *Tridrepana unispina* | Drepanoidea | Drepanidae | AYK-04-0776-02 | Tuni |
| *Habrosyne pyritoides* | Drepanoidea | Drepanidae | AYK-04-5353 | Hbpy |
| *Pseudothyatira cymatophoroides* | Drepanoidea | Drepanidae | CWM-94-0380 | Pcym |
| *Tethea consimilis* | Drepanoidea | Drepanidae | AYK-04-5374 | Ttms |
| *Euparyphasma maxima* | Drepanoidea | Drepanidae | AYK-04-5376 | Emxm |
| *Parapsestis argenteopicta* | Drepanoidea | Drepanidae | AYK-07-7617 | Parg |
| *Thyatira batis* | Drepanoidea | Drepanidae | AYK-04-2520 | Tbts3 |

| | | | | |
|---|---|---|---|---|
| *Epicopeia hainesii* | Drepanoidea | Epicopeiidae | AYK 04 5730 | Ehai |
| *Psychostrophia melangaria* | Drepanoidea | Epicopeiidae | AYK 04 5216 | Pmel |
| *Epermenia chaerophyllella* | Epermenioidea | Epermeniidae | DCL-07-0006 | Epchh |
| *Epermenia sinjovi* | Epermenioidea | Epermeniidae | JCS-08-1020 | Esji |
| *Epermenia stolidota* | Epermenioidea | Epermeniidae | JWB-07-2003 | Epst |
| *Homadaula anisocentra* | Galacticoidea | Galacticidae | JWB-05-0026 | Hani |
| *Autosticha modicella* | Gelechioidea | Autostichidae | JCS-06-0175 | Amod |
| *Deoclona yuccasella* | Gelechioidea | Autostichidae | OP-04-0131 | Deoy |
| *Galagete protozona* | Gelechioidea | Autostichidae | PS-05-2001 | Gprot2 |
| *Batrachedra pinicolella* | Gelechioidea | Batrachedridae | JCS-08-1054 | Bpin |
| *Neoblastobasis spiniharpella* | Gelechioidea | Blastobasidae | JCS-06-0126 | Nspin |
| *Diurnea fagella* | Gelechioidea | Chimabachidae | MM-08-5965 | Dfag |
| *Coleophora cratipennella* | Gelechioidea | Coleophoridae | KTP-94-0520 | Clph |
| *Coleophora artemisicolella* | Gelechioidea | Coleophoridae | SWC-06-0208 | Cpta |
| *Euclemensia bassettella* | Gelechioidea | Cosmopterigidae | LS-06-0163 | Euba |
| *Pancalia schwartzella* | Gelechioidea | Cosmopterigidae | SWC-07-2001 | Panla |
| *Pyroderces japonica* | Gelechioidea | Cosmopterigidae | AYK-04-5635 | Ajnc |
| *Hyposmochoma turdella* | Gelechioidea | Cosmopterigidae | DR-08-C1C-3 | Htur |
| *Aeolanthes semiostrina* | Gelechioidea | Elachistidae | JCS-06-0112 | Asem |
| *Agonopterix alstroemeriana* | Gelechioidea | Elachistidae | JWB-06-0048 | Agpt |
| *Bibarrambla allenella* | Gelechioidea | Elachistidae | LS-06-0168 | Ball |
| *Filinota brunniceps* | Gelechioidea | Elachistidae | KN-06-0579 | Fbrn |
| *Elachista illectella* | Gelechioidea | Elachistidae | SWC-05-0101 | Cill |
| *Elachista tengstromi* | Gelechioidea | Elachistidae | MM-08-5412 | Eten |
| *Ethmia eupostica* | Gelechioidea | Elachistidae | MJM-96-0277 | Eeu |
| *Eupselia carpocapsella* | Gelechioidea | Elachistidae | MJM-96-0291 | Epca |
| *Hypertropha tortriciformis* | Gelechioidea | Elachistidae | MJM-96-0240 | Hptr |
| *Thudaca haplonota* | Gelechioidea | Elachistidae | AZ-07-2140 | Thap |
| *Antaeotricha renselariana* | Gelechioidea | Elachistidae | 05-srnp-05145 | Aren |
| *Stenoma patens* | Gelechioidea | Elachistidae | 05-srnp-04076 | Stpa |
| *Psilocorsis reflexella* | Gelechioidea | Elachistidae | CWM-94-0267 | Prfx |
| *Aristotelia mesotenebrella* | Gelechioidea | Gelechiidae | JCS-06-0169 | Amst |

| | | | | |
|---|---|---|---|---|
| *Tenupalpa biformis* | Gelechioidea | Gelechiidae | AYK-04-5612 | Thio |
| *Monochroa cleodoroides* | Gelechioidea | Gelechiidae | SWC-07-2018 | Mcls |
| *Pectinophora gossypiella* | Gelechioidea | Gelechiidae | BT-06-3360-1 | Pgos |
| *Dichomeris punctidiscella* | Gelechioidea | Gelechiidae | CWM-94-0259 | Dpunc |
| *Encolapta tegulifera* | Gelechioidea | Gelechiidae | JCS-06-0124 | Etgf |
| *Faristenia furtumella* | Gelechioidea | Gelechiidae | JCS-06-0150 | Ffu |
| *Aroga trialbomaculella* | Gelechioidea | Gelechiidae | CWM-94-0264 | Arot |
| *Caryocolum pulatella* | Gelechioidea | Gelechiidae | SWC-07-2047 | Cpua |
| *Friseria cockerelli* | Gelechioidea | Gelechiidae | seto_dystria | seto_dystria |
| *Exoteleia pinifoliella* | Gelechioidea | Gelechiidae | KTP-94-0513 | Epif |
| *Hypatima mediofasciana* | Gelechioidea | Gelechiidae | JCS-06-0121 | Hmdf |
| *Teleiodes pekunensis* | Gelechioidea | Gelechiidae | JCS-06-0130 | Tpku |
| *Homaloxestis croceata* | Gelechioidea | Lecithoceridae | SWC-06-0243 | Htce |
| *Lecithocera cheristis* | Gelechioidea | Lecithoceridae | KTP-06-0113-1 | Leci |
| *Odites leucostola* | Gelechioidea | Lecithoceridae | AYK-04-5610 | Odle |
| *Torodora babeana* | Gelechioidea | Lecithoceridae | KTP-06-0102-1 | Tbbn |
| *Rhyzosthenes falciformis* | Gelechioidea | Lecithoceridae | JCS-06-0155 | Rfal |
| *Tisis mesozosta* | Gelechioidea | Lecithoceridae | LS-06-0056 | Tisi |
| *Lypusa maurella* | Gelechioidea | Lypusidae | DRD-07-4102 | Lmau |
| *Mompha cephalonthiella* | Gelechioidea | Momphidae | DLW-90-0014 | Mom |
| *Deuterogonia pudorina* | Gelechioidea | Oecophoridae | JCS-06-0144 | Dpd |
| *Idioglossa miraculosa* | Gelechioidea | Oecophoridae | TH-08-6213 | Imir |
| *Mimobrachyoma hilaropa* | Gelechioidea | Oecophoridae | MJM-96-0260 | Mhil |
| *Promalactis jezonica* | Gelechioidea | Oecophoridae | KTP-06-0105 | Pjez |
| *Wingia aurata* | Gelechioidea | Oecophoridae | MJM-97-0295 | Win |
| *Acria ceramitis* | Gelechioidea | Peleopodidae | AYK-04-5621 | Acer |
| *Scythris immaculatella* | Gelechioidea | Scythrididae | LS-06-0164 | Simm |
| *Hieromantis kurokoi* | Gelechioidea | Stathmopodidae | JCS-06-0157 | Hiku |
| *Stathmopoda melanochra* | Gelechioidea | Stathmopodidae | MJM-97-0301 | Stmp |
| *Leistarcha scitissimella* | Gelechioidea | Xyloryctidae | MJM-97-0314 | Lsci |
| *Tymbophora peltastis* | Gelechioidea | Xyloryctidae | MJM-97-0320 | Tymb |

| | | | | |
|---|---|---|---|---|
| *Alsophila pometaria* | Geometroidea | Geometridae | CWM-06-1013 | Ahpt |
| *Archiearis parthenias* | Geometroidea | Geometridae | NH-06-0806 | Apar |
| *Lachnocephala vellosata* | Geometroidea | Geometridae | AH-07-0001 | Lvst |
| *Biston betularia* | Geometroidea | Geometridae | CWM-94-0127 | Bbet |
| *Campaea perlata* | Geometroidea | Geometridae | JCR-94-0143 | Cper |
| *Hasodima elegans* | Geometroidea | Geometridae | AH-07-0028 | Hens |
| *Amelora megalocephala** | Geometroidea | Geometridae | AH-07-7301 | Ameg |
| *Chiasmia clathrata* | Geometroidea | Geometridae | AH-07-3740 | Ccla |
| *Plagodis fervidaria* | Geometroidea | Geometridae | CWM-94-0124 | Pfer |
| *Selenia dentaria* | Geometroidea | Geometridae | AH-07-3656 | Sbnr |
| *Chlorosea margaretaria* | Geometroidea | Geometridae | RR 98 1152 | Cmar |
| *Hypobapta xenomorpha* | Geometroidea | Geometridae | MJM 96 0284 | Hxen |
| *Earophila badiata* | Geometroidea | Geometridae | AH-07-3746 | Ebdt |
| *Eupithecia acutipennis* | Geometroidea | Geometridae | NB-06-0044 | Euac |
| *Orthonama obstipata* | Geometroidea | Geometridae | RR-98-0917 | Onst |
| *Trichopteryx carpinata* | Geometroidea | Geometridae | AH-05-3644 | Trca |
| *Aplocera efformata* | Geometroidea | Geometridae | AH-07-3780 | Aeff |
| *Cidaria fulvata* | Geometroidea | Geometridae | AH-07-3815 | Cfta |
| *Hydrelia flammeolaria* | Geometroidea | Geometridae | AH-07-3803 | Hfmm |
| *Cyclophora nanaria* | Geometroidea | Geometridae | RR 98 1138 | Cytr |
| *Haematopis grataria* | Geometroidea | Geometridae | AM-94-0358 | Hrgi |
| *Idaea dismissaria* | Geometroidea | Geometridae | RR-98-1137 | Idms |
| *Scopula limboundata* | Geometroidea | Geometridae | CWM 94 0285 | Scli |
| *Rhodometra sacraria* | Geometroidea | Geometridae | AH-07-3956 | Rsac |
| *Coronidia orithea* | Geometroidea | Sematuridae | 05-srnp-19273 | Cooh |
| *Sematura luna* | Geometroidea | Sematuridae | RWH-96-0877 | Nothsp |
| *Metorthocheilus emarginatus* | Geometroidea | Uraniidae | LS-06-0067 | Meem |
| *Syngria druidaria* | Geometroidea | Uraniidae | VOB-00-9803 | Sdru |
| *Calledapteryx dryopterata* | Geometroidea | Uraniidae | CWM-96-0576 | Cdry |
| *Erosia veninotata* | Geometroidea | Uraniidae | 06-srnp-35124 | Evnt |
| *Phazaca interrupta* | Geometroidea | Uraniidae | AZ-07-2139 | Pitr |
| *Schidax squamaria* | Geometroidea | Uraniidae | 07-srnp-57826 | Ssqu |
| *Acropteris sparsaria* | Geometroidea | Uraniidae | LS-06-0060 | Acro |
| *Lyssa zampa* | Geometroidea | Uraniidae | A-0581 | Lza |
| *Urapteroides astheniata* | Geometroidea | Uraniidae | AYK-04-1274-1 | Uptd3 |
| *Bucculatrix staintonella* | Gracillarioidea | Bucculatricidae | TH-08-6083 | Bsta |
| *Klimeschia transversella* | Gracillarioidea | Douglasiidae | DRD-01-0017 | Ktr |
| *Tinagma gaedikei* | Gracillarioidea | Douglasiidae | TH-08-6082 | Tgak |
| *Tinagma ocnerostomella* | Gracillarioidea | Douglasiidae | DRD-01-0013 | Tocn |

| | | | | |
|---|---|---|---|---|
| *Caloptilia bimaculatella* | Gracillarioidea | Gracillariidae | DRD-05-0248 | Cbim |
| *Callisto denticulella* | Gracillarioidea | Gracillariidae | AYK-08-8214 | Cdel |
| *Epicephala relictella* | Gracillarioidea | Gracillariidae | JCS-06-0172 | Epic |
| *Parectopa robiniella* | Gracillarioidea | Gracillariidae | DRD-01-0009 | Prbn |
| *Spulerina dissotoma* | Gracillarioidea | Gracillariidae | AYK-04-5630 | Sput |
| *Acrocercops albinatella* | Gracillarioidea | Gracillariidae | TH-08-6107 | Aalne |
| *Acrocercops brongniardella* | Gracillarioidea | Gracillariidae | AYK-08-8215 | Abrg |
| *Acrocercops transecta* | Gracillarioidea | Gracillariidae | AYK-08-8201 | Atran |
| *Artifodina japonica* | Gracillarioidea | Gracillariidae | AYK-08-8206 | Ajaa |
| *Caloptilia murtfeldtella* | Gracillarioidea | Gracillariidae | TH-08-6096 | Cmur |
| *Caloptilia sapporella* | Gracillarioidea | Gracillariidae | JCS-08-1033 | Csap |
| *Caloptilia stigmatella* | Gracillarioidea | Gracillariidae | JDP-08-8081 | Cstg |
| *Calybites auroguttella* | Gracillarioidea | Gracillariidae | AYK-08-8218 | Caug |
| *Conopomorpha cramerella* | Gracillarioidea | Gracillariidae | AYK-08-8231 | Ccrm |
| *Cuphodes diospyrosella* | Gracillarioidea | Gracillariidae | AYK-08-8223 | Cdio |
| *Dendrorycter marmaroides* | Gracillarioidea | Gracillariidae | PZ028784 | Dend |
| *Deoptilia heptadeta* | Gracillarioidea | Gracillariidae | PZ028785 | Deoa |
| *Acrocercops scriptulata* | Gracillarioidea | Gracillariidae | AYK-08-8224 | Dscr |
| *Eteoryctis deversa* | Gracillarioidea | Gracillariidae | AYK-08-8203 | Edev |
| *Eucalybites aureola* | Gracillarioidea | Gracillariidae | PZ028786 | Euau |
| *Gibbovalva quadrifasciata* | Gracillarioidea | Gracillariidae | PZ028787 | Gibb |
| *Gracillaria syringella* | Gracillarioidea | Gracillariidae | JDP-08-8042 | Gsyg |
| *Leucocephala schinusae* | Gracillarioidea | Gracillariidae | DRD-07-4001 | Gran |
| *Leucospilapteryx venustella* | Gracillarioidea | Gracillariidae | TH-08-6105 | Lven |
| *Liocrobyla lobata* | Gracillarioidea | Gracillariidae | PZ028788 | Lioc |
| *Marmara serotinella* | Gracillarioidea | Gracillariidae | DRD-05-0265 | Msll |
| *Micrurapteryx salicifoliella* | Gracillarioidea | Gracillariidae | TH-08-6098 | Msai |
| *Neurobathra strigifinitella* | Gracillarioidea | Gracillariidae | PZ028789 | Neur |
| *Parornix anglicella* | Gracillarioidea | Gracillariidae | AYK-08-8210 | Pxag |
| *Parornix torquillella* | Gracillarioidea | Gracillariidae | AYK-08-8213 | Pota |
| *Phodoryctis stephaniae* | Gracillarioidea | Gracillariidae | AYK-08-8204 | Pste |
| *Macrosaccus robiniella* | Gracillarioidea | Gracillariidae | JDP-08-8001 | Proi |
| *Povolnya obliquatella* | Gracillarioidea | Gracillariidae | SWC-06-0265 | Pvob |
| *Psydrocercops wisteriae* | Gracillarioidea | Gracillariidae | AYK-08-8221 | Pwis |

| | | | | |
|---|---|---|---|---|
| *Cameraria gaultheriella* | Gracillarioidea | Gracillariidae | DRD-01-0113v | Caga |
| *Afrorycter nsengai* | Gracillarioidea | Gracillariidae | AK-07-135 | Afsp |
| *Cameraria guttifinitella* | Gracillarioidea | Gracillariidae | TH-08-6116 | Cgut |
| *Cameraria ohridella* | Gracillarioidea | Gracillariidae | JDP-08-8021 | Cohd |
| *Chrysaster hagicola* | Gracillarioidea | Gracillariidae | AYK-08-8202 | Chag |
| *Chrysaster ostensackenella* | Gracillarioidea | Gracillariidae | DRD-05-0247 | Cyosk |
| *Cremastobombycia solidaginis* | Gracillarioidea | Gracillariidae | TH-08-6130 | Cmys |
| *Hyloconis luki* | Gracillarioidea | Gracillariidae | AK-07-121 | Porp |
| *Hyloconis wisteriae* | Gracillarioidea | Gracillariidae | PZ028790 | Hwis |
| *Leucanthiza amphicarpeaefoliella* | Gracillarioidea | Gracillariidae | PZ028791 | Leuz |
| *Neolithocolletis hikomonticola* | Gracillarioidea | Gracillariidae | PZ028792 | Nlik |
| *Phyllonorycter basistrigella* | Gracillarioidea | Gracillariidae | TH-08-6111 | Pbas |
| *Phyllonorycter insignitella* | Gracillarioidea | Gracillariidae | PZ028793 | Phzz |
| *Phyllonorycter lucetiella* | Gracillarioidea | Gracillariidae | TH-08-6117 | Ptea |
| *Phyllonorycter ostryaefoliella* | Gracillarioidea | Gracillariidae | TH-08-6103 | Posy |
| *Phyllonorycter symphoricarpaeella* | Gracillarioidea | Gracillariidae | TH-08-6128 | Pmicp |
| *Eumetriochroa hederae* | Gracillarioidea | Gracillariidae | AK-08-8111 | Ehdr |
| *Phyllocnistis citrella* | Gracillarioidea | Gracillariidae | JBA-06-0025 | Phcn |
| *Phyllocnistis longipalpus* | Gracillarioidea | Gracillariidae | DRD-05-0244 | Pmgl |
| *Anartioses aberrans* | Gracillarioidea | Gracillariidae | DRD-01-0010 | Anab |
| *Chilocampyla dyariella* | Gracillarioidea | Gracillariidae | EJN-06-2539 | Cdya.2 |
| *Macarostola japonica* | Gracillarioidea | Gracillariidae | AYK-08-8238 | Mjap |
| *Agriothera elaeocarpophaga* | Gracillarioidea | Roeslerstammidae | DRD-01-0143 | Agel |
| *Roeslerstammia pronubella* | Gracillarioidea | Roeslerstammidae | DRD-01-0142 | Rstm |
| *Macrosoma conifera* | Hedyloidea | Hedylidae | VOB-00-9811 | Mcon |
| *Macrosoma satellitiata satellitiata* | Hedyloidea | Hedylidae | MEE_98-0042 | Mssa |
| *Coeliades ramanatek* | Papilionoidea | Hesperiidae | DCL-07-0011 | Cram |
| *Hasora chromus* | Papilionoidea | Hesperiidae | MFB-07-003 | Hsch |
| *Astraptes enotrus* | Papilionoidea | Hesperiidae | DHJ-02-2406 | Aeno |
| *Dyscophellus phraxanor* | Papilionoidea | Hesperiidae | 05-srnp-32396 | Dpxn |
| *Venada nevada* | Papilionoidea | Hesperiidae | 05-srnp-35622 | Vnev |
| *Urbanus doryssus* | Papilionoidea | Hesperiidae | 05-srnp-46521 | Udo |
| *Calpodes ethlius* | Papilionoidea | Hesperiidae | 05-srnp-59316 | Cet |
| *Orses cynisca* | Papilionoidea | Hesperiidae | 05-srnp-07056 | Ocyn |

| | | | | |
|---|---|---|---|---|
| *Cyclosemia herennius subcaerulea* | Papilionoidea | Hesperiidae | 05-srnp-42245 | Cycs3 |
| *Morvina fissimacula* | Papilionoidea | Hesperiidae | 05-srnp-04126 | Mfmc |
| *Mysoria ambigua* | Papilionoidea | Hesperiidae | DHJ-02-2459 | Myam |
| *Hyblaea ibidias* | Hyblaeoidea | Hyblaeidae | AZ-06-0176 | Hibd |
| *Hyblaea firmamentum* | Hyblaeoidea | Hyblaeidae | LS-06-0069 | Hyfm |
| *Imma tetrascia* | Immoidea | Immidae | MJM-95-0169 | Imsp |
| *Birthana cleis* | Immoidea | Immidae | MFB-06-0101 | Bcle |
| *Imma loxoscia* | Immoidea | Immidae | AZ-07-2692 | Imma |
| *Munychryia senicula* | Bombycoidea | Anthelidae | AZ-06-0052 | Muyc |
| *Chelepteryx collesi* | Bombycoidea | Anthelidae | MJM-97-0329 | Ccol |
| *Nataxa flavescens* | Bombycoidea | Anthelidae | MJM-96-0289 | Nfla |
| *Chionopsyche montana* | Lasiocampoidea | Lasiocampidae | AZ-06-0100 | Cmtn |
| *Chondrostega vandalicia* | Lasiocampoidea | Lasiocampidae | JBA-09-3002 | Cvan |
| *Heteropacha rileyana* | Lasiocampoidea | Lasiocampidae | RSP-96-0003-1 | Hril |
| *Lasiocampa quercus* | Lasiocampoidea | Lasiocampidae | RSP-96-0862 | Lquercus |
| *Malacosoma americanum* | Lasiocampoidea | Lasiocampidae | AM-94-0145 | Mame2 |
| *Macrothylacia rubi* | Lasiocampoidea | Lasiocampidae | RSP-xx-0840 | Mrubi |
| *Eutachyptera psidii* | Lasiocampoidea | Lasiocampidae | PZ028794 | Epsidii |
| *Genduara acedesta* | Lasiocampoidea | Lasiocampidae | PZ028795 | Gace |
| *Gonometa rufobrunnea* | Lasiocampoidea | Lasiocampidae | PZ028796 | Grufo |
| *Malacosoma californicum* | Lasiocampoidea | Lasiocampidae | PZ028797 | Mcaliforn |
| *Phyllodesma americana* | Lasiocampoidea | Lasiocampidae | PZ028798 | Pamerica |
| *Tolype notialis* | Lasiocampoidea | Lasiocampidae | KRH-94-2004 | Tnot |
| *Artace cribraria* | Lasiocampoidea | Lasiocampidae | AM-94-0422 | Acri |
| *Poecilocampa populi* | Lasiocampoidea | Lasiocampidae | IJK-05-0001 | Ppop |
| *Bedosia turgida* | Mimallonoidea | Mimallonidae | VOB-01-0010 | Bedg2 |
| *Druentia alsa* | Mimallonoidea | Mimallonidae | VOB-00-8545 | Dals |
| *Euphaneta divisa* | Mimallonoidea | Mimallonidae | VOB-00-0462 | Edvs3 |
| *Lacosoma chiridota* | Mimallonoidea | Mimallonidae | JKA-95-0013 | Lch2 |
| *Mimallo amilia* | Mimallonoidea | Mimallonidae | 05-srnp-62160 | Moii |
| *Asota egens confinis* | Noctuoidea | Erebidae | AYK-04-0802-15 | Asoa |
| *Amata fortunei* | Noctuoidea | Erebidae | AYK-04-5607 | Afor |
| *Cisseps fulvicollis* | Noctuoidea | Erebidae | AM-93-0003 | Cfu2 |
| *Hypoprepia miniata* | Noctuoidea | Erebidae | RFD-93-0437 | Hymi |

| | | | | |
|---|---|---|---|---|
| *Balacra pulchra* | Noctuoidea | Erebidae | AYK-07-9197 | Bpul |
| *Mycterophora rubricans* | Noctuoidea | Erebidae | RR-97-0786 | Mrcn |
| *Eudocima salaminia* | Noctuoidea | Erebidae | AYK-07-7601 | Esal |
| *Gonodonta fulvangula* | Noctuoidea | Erebidae | 05-srnp-56095 | Gono |
| *Catocala ultronia* | Noctuoidea | Erebidae | CWM-93-0435 | Caul |
| *Diascia hayesi* | Noctuoidea | Erebidae | MF-06-0134 | Acan |
| *Erebus ephesperis* | Noctuoidea | Erebidae | AYK-04-0792-02 | Ereb |
| *Zale lunifera* | Noctuoidea | Erebidae | RFD-94-0021 | Zalu |
| *Palthis asopialis* | Noctuoidea | Erebidae | AM-93-0436 | Paang |
| *Hypena scabra* | Noctuoidea | Erebidae | RWP-93-1001 | Psc2 |
| *Lymantria dispar* | Noctuoidea | Erebidae | UNK-93-0047 | Ldi |
| *Orgyia leucostigma* | Noctuoidea | Erebidae | CWM-95-0465 | Orgy |
| *Pangrapta decoralis* | Noctuoidea | Erebidae | CWM-94-0365 | Pdrls |
| *Rivula propinqualis* | Noctuoidea | Erebidae | CWM-96-0957 | Rpro3 |
| *Phobolosia anfracta* | Noctuoidea | Erebidae | RR-95-0103 | Phfr |
| *Anomis metaxantha* | Noctuoidea | Erebidae | AYK-04-0786-16 | Anmet |
| *Micronoctua karsholti* | Noctuoidea | Micronoctuidae | MF-06-0146 | Mka |
| *Acontia aprica* | Noctuoidea | Noctuidae | JKA-97-0011 | Aaca |
| *Acronicta lobeliae* | Noctuoidea | Noctuidae | CWM-94-0230 | Albe |
| *Eudryas grata* | Noctuoidea | Noctuidae | AM-93-0448 | Egta |
| *Amphipyra pyramidoides* | Noctuoidea | Noctuidae | CWM-94-0371 | Amph |
| *Psaphida resumens* | Noctuoidea | Noctuidae | RFD-94-0035 | Puen |
| *Stiria rugifrons* | Noctuoidea | Noctuidae | EK-92-0184 | Srgf |
| *Sphragifera sigillata* | Noctuoidea | Noctuidae | AYK-04-2537 | Ssig |
| *Cryphia cuerva* | Noctuoidea | Noctuidae | RR-99-1237-2 | Cnae |
| *Condica vecors* | Noctuoidea | Noctuidae | CWM-95-0471 | Cvec |
| *Cucullia convexipennis* | Noctuoidea | Noctuidae | EK-xx-0379 | Cuvx |
| *Raphia abrupta* | Noctuoidea | Noctuidae | CWM-94-0372 | Rabp |
| *Helicoverpa zea* | Noctuoidea | Noctuidae | RWP-94-0187 | Hzea |
| *Adisura bella* | Noctuoidea | Noctuidae | MJM-88-0119 | Abel |
| *Pyrrhia cilisca* | Noctuoidea | Noctuidae | JKA-95-0002 (v.208) | Pumb |

| | | | | |
|---|---|---|---|---|
| *Agrotis ipsilon* | Noctuoidea | Noctuidae | AM-93-0451 | Agip |
| *Caradrina meralis* | Noctuoidea | Noctuidae | RR-96-0224 | Platc |
| *Mythimna unipuncta* | Noctuoidea | Noctuidae | RWP-87-0438-2 | Psdl |
| *Pseudeustrotia carneola* | Noctuoidea | Noctuidae | CWM-93-0001 | Lcart |
| *Xylena exsoleta* | Noctuoidea | Noctuidae | NH-96-0808 | Xelt |
| *Spodoptera frugiperda* | Noctuoidea | Noctuidae | JN-92-0194 | Sfr |
| *Trichosea champa* | Noctuoidea | Noctuidae | AYK-04-5234 | Tchmp |
| *Trichoplusia ni* | Noctuoidea | Noctuidae | JN-92-0192 | Tni |
| *Blenina senex* | Noctuoidea | Nolidae | AYK-04-5172 | Bsex |
| *Negeta signata* | Noctuoidea | Nolidae | AYK-04-0515 | Nsig |
| *Iscadia producta* | Noctuoidea | Nolidae | 05-srnp-07372 | Ipdt |
| *Earias roseifera* | Noctuoidea | Nolidae | AYK-04-0824-07 | Erof |
| *Meganola minuscula* | Noctuoidea | Nolidae | CWM-93-0434 | Mmi1 |
| *Scotura leucophleps* | Noctuoidea | Notodontidae | 05-srnp-22116 | Stul |
| *Crinodes besckei* | Noctuoidea | Notodontidae | DHJ-02-2285 | Cbes |
| *Hemiceras nigrescens* | Noctuoidea | Notodontidae | 05-srnp-19566 | Hngrs |
| *Dicentria violacens* | Noctuoidea | Notodontidae | 05-srnp-23178 | Dvac |
| *Heterocampa obliqua* | Noctuoidea | Notodontidae | CWM-94-0317 | Hbiud |
| *Rhuda difficilis* | Noctuoidea | Notodontidae | 05-srnp-32496 | Rdif |
| *Schizura unicornis* | Noctuoidea | Notodontidae | AM-93-0432 | Sipo |
| *Furcula cinerea* | Noctuoidea | Notodontidae | AM-93-0430 | Fci2 |
| *Gluphisia septentrionis* | Noctuoidea | Notodontidae | CWM-94-0297 | Gsep |
| *Nystalea striata* | Noctuoidea | Notodontidae | 05-srnp-03781 | Nstr |
| *Symmerista albifrons* | Noctuoidea | Notodontidae | CWM-94-0347 | Safn |
| *Datana drexelii* | Noctuoidea | Notodontidae | AM-93-0431/v.044 | Dpe2 |
| *Rosema epigena* | Noctuoidea | Notodontidae | 07-srnp-3604 | Repi |
| *Spatalia doerriesi* | Noctuoidea | Notodontidae | AYK-04-2513 | Sdoe |
| *Hapigia nodicornis* | Noctuoidea | Notodontidae | 05-srnp-05815 | Hapno |
| *Cnethodonta grisescens* | Noctuoidea | Notodontidae | AYK-04-2514 | Cgris |
| *Cerura rarata* | Noctuoidea | Notodontidae | 07-srnp-58332 | Crrt |
| *Thaumatopoea pityocampa* | Noctuoidea | Notodontidae | JBA-97-042 | Tpit |
| *Epanaphe carteri* | Noctuoidea | Notodontidae | AYK-07-9170 | Ecrt |

| | | | | |
|---|---|---|---|---|
| *Ochrogaster lunifer* | Noctuoidea | Notodontidae | AZ-07-3148-2 | Ocgs |
| *Lirimiris guatemalensis* | Noctuoidea | Notodontidae | 05-srnp-58695 | Lgua |
| *Ptilophora plumigera* | Noctuoidea | Notodontidae | NH-96-0809 | Ppga |
| *Oenosandra boisduvalii* | Noctuoidea | Oenosandridae | AZ-06-0157 | Oeno |
| *Curetis bulis stigmata* | Papilionoidea | Lycaenidae | MWT-93-A028 | Cure |
| *Pseudozizeeria maha* | Papilionoidea | Lycaenidae | AYK-04-5608 | Psma |
| *Liphyra brassolis* | Papilionoidea | Lycaenidae | KD-94-T063 | Liph |
| *Poritia erycinoides* | Papilionoidea | Lycaenidae | MWT-93-B007 | Pedy |
| *Eumaeus godarti* | Papilionoidea | Lycaenidae | 05 srnp 33799 | Egod |
| *Strymon eurytulus* | Papilionoidea | Lycaenidae | AYK-04-0543-11 | Seur2 |
| *Mechanitis polymnia* | Papilionoidea | Nympalidae | 05-srnp-32074 | Mpol |
| *Asterocampa celtis* | Papilionoidea | Nymphalidae | TPF-89-0011 | Acly |
| *Dynamine sosthenes* | Papilionoidea | Nymphalidae | 06-srnp-32108 | Dsos |
| *Hamadryas arinome* | Papilionoidea | Nymphalidae | 05-srnp-42608 | Hama |
| *Prepona demophon* | Papilionoidea | Nymphalidae | DHJ-02-2374 | Arch |
| *Marpesia petreus* | Papilionoidea | Nymphalidae | 05-srnp-56870 | Mape |
| *Danaus plexippus* | Papilionoidea | Nymphalidae | 05-srnp-05455 | Dplex |
| *Eueides isabella* | Papilionoidea | Nymphalidae | 05-srnp-06244 | Euis |
| *Heliconius sara* | Papilionoidea | Nymphalidae | 05-srnp-06959 | Heli3 |
| *Libytheana carinenta bachmanii* | Papilionoidea | Nymphalidae | AYK-01-0004 | Lica |
| *Limenitis arthemis* | Papilionoidea | Nymphalidae | LS-06-0165 | Larth |
| *Hypanartia godmani* | Papilionoidea | Nymphalidae | DHJ-02-2464 | Hgod |
| *Phyciodes phaon* | Papilionoidea | Nymphalidae | AV-91-0087 | Ptha |
| *Vanessa carye* | Papilionoidea | Nymphalidae | AYK-04-0538-03 | Vane |
| *Caligo telamonius* | Papilionoidea | Nymphalidae | 07-srnp-58684 | Ctel |
| *Lethe sicelis* | Papilionoidea | Nymphalidae | MS-08-0922 | Lsie |
| *Manataria hercyna* | Papilionoidea | Nymphalidae | 05-srnp-19327 | Mmct |
| *Mycalesis gotama* | Papilionoidea | Nymphalidae | MS-08-0923 | Mgot |
| *Neope goschkevitschii* | Papilionoidea | Nymphalidae | AYK-06-7212 | Ngos |
| *Pierella luna* | Papilionoidea | Nymphalidae | DHJ-02-2427 | Plun |
| *Taygetis andromeda* | Papilionoidea | Nymphalidae | 05-srnp-59405 | Tadm |
| *Eurytides branchus* | Papilionoidea | Papilionidae | DHJ-02-2433 | Euryt |

| | | | | |
|---|---|---|---|---|
| *Papilio glaucus* | Papilionoidea | Papilionidae | CWM-02-0138 | Pgla |
| *Parides iphidamas* | Papilionoidea | Papilionidae | 05-srnp-63425 | Piph3 |
| *Parnassius apollo* | Papilionoidea | Papilionidae | JBA-06-0019 | Pnpl |
| *Colias eurytheme* | Papilionoidea | Pieridae | AV-91-ceur | Ceur |
| *Eurema boisduvaliana* | Papilionoidea | Pieridae | 05-srnp-62336 | Exnt |
| *Kricogonia lyside* | Papilionoidea | Pieridae | SW-07-0026 | Klyd3 |
| *Dismorphia amphiona* | Papilionoidea | Pieridae | 05-srnp-05463 | Damp |
| *Pereute charops* | Papilionoidea | Pieridae | kn-06-1126 | Pcrps3 |
| *Pieris rapae* | Papilionoidea | Pieridae | JWB-06-0001 | Prap |
| *Pseudopontia paradoxa* | Papilionoidea | Pieridae | AV-92-0002 | Ptdx |
| *Euselasia mystica* | Papilionoidea | Riodinidae | 07-srnp-59426 | Emh |
| *Emesis lucinda* | Papilionoidea | Riodinidae | 05-srnp-31537 | Elu |
| *Mesosemia carissima* | Papilionoidea | Riodinidae | 06-srnp-7404 | Mcsm |
| *Napaea eucharila* | Papilionoidea | Riodinidae | 06-srnp-43842 | Neuc |
| *Symmachia tricolor* | Papilionoidea | Riodinidae | 06-srnp-44831 | Stcr |
| *Agdistis americana* | Pterophoroidea | Pterophoridae | NB-06-0144 | Adam |
| *Agdistopis sinhala* | Pterophoroidea | Pterophoridae | LS-06-0059 | Agdi |
| *Anstenoptilia marmarodactyla* | Pterophoroidea | Pterophoridae | NB-06-0002 | Anma |
| *Emmelina monodactyla* | Pterophoroidea | Pterophoridae | CWM-07-5001 | Emon |
| *Platyptilia ignifera* | Pterophoroidea | Pterophoridae | AYK-04-5601 | Plty |
| *Petrophila confusalis* | Pyraloidea | Crambidae | RR-98-1148 | Pcon |
| *Catoptria oregonicus* | Pyraloidea | Crambidae | RR-98-1176 | Caor |
| *Chilo suppressalis* | Pyraloidea | Crambidae | MAS-92-1001-1 | Csss |
| *Crambus agitatellus* | Pyraloidea | Crambidae | JWB-08-0114 | Cagt |
| *Crocidolomia luteolalis* | Pyraloidea | Crambidae | AZ-07-2685 | Ccdm |
| *Evergestis subterminalis* | Pyraloidea | Crambidae | RR-98-1157 | Esub |
| *Chalcoela iphitalis* | Pyraloidea | Crambidae | CWM-06-1016 | Ciph |
| *Cosmopterosis spatha* | Pyraloidea | Crambidae | 06-SRNP-23180 | Coth |
| *Dichogama gudmanni* | Pyraloidea | Crambidae | 05-srnp-18221 | Dcth |
| *Dicymolomia metalliferalis* | Pyraloidea | Crambidae | RR-99-1228 | Dmet |
| *Cacographis osteolalis* | Pyraloidea | Crambidae | MAS-91-0407-1 | Coste |
| *Dismidila atoca* | Pyraloidea | Crambidae | MAS-06-0202 | Dioc |
| *Midila daphne* | Pyraloidea | Crambidae | MAS-06-0201 | Miph |
| *Neurophyseta conantia* | Pyraloidea | Crambidae | MAS-91-0601 | Muso |

| | | | | |
|---|---|---|---|---|
| *Noorda blitealis* | Pyraloidea | Crambidae | WM-08-4009 | Nblt |
| *Cliniodes opalalis* | Pyraloidea | Crambidae | 05-srnp-7932 | Clop |
| *Syntonarcha iriastis* | Pyraloidea | Crambidae | AZ-07-2650 | Stna |
| *Ostrinia furnacalis* | Pyraloidea | Crambidae | MAS-92-0801 | Osfu |
| *Pyrausta nexalis* | Pyraloidea | Crambidae | RR-98-1141 | Pnex |
| *Rupela albina* | Pyraloidea | Crambidae | MAS-91-0209 | Psh29 |
| *Scirpophaga incertulas* | Pyraloidea | Crambidae | MAS 92 1003 | Sin |
| *Eudonia spenceri* | Pyraloidea | Crambidae | RR 98 1146 | Scsp |
| *Scoparia isochroalis* | Pyraloidea | Crambidae | SWC-06-0240 | Sira |
| *Diaphania indica* | Pyraloidea | Crambidae | AYK-04-5705 | Dnin |
| *Mesocondyla dardusalis* | Pyraloidea | Crambidae | MAS-91-0224 | Ppy224 |
| *Phaeodropsis alitemeralis* | Pyraloidea | Crambidae | MAS-91-0122 | Ppy122 |
| *Niphopyralis chionesis* | Pyraloidea | Crambidae | AZ-07-2642 | Niph |
| *Monoloxis flavicinctalis* | Pyraloidea | Pyralidae | 05-srnp-24832 | Mfla |
| *Polyterpnes polyrrhoda* | Pyraloidea | Pyralidae | AZ-07-2643 | Aphy |
| *Accinctapubes albifasciata* | Pyraloidea | Pyralidae | 05-srnp-46779 | Acal |
| *Salma pyrastis* | Pyraloidea | Pyralidae | MJM-97-0297 | Spis |
| *Galleria melonella* | Pyraloidea | Pyralidae | CWM-08-2532 | Gmel |
| *Ambesa laetella* | Pyraloidea | Pyralidae | RR-98-1191 | Ambe |
| *Dioryctria auranticella* | Pyraloidea | Pyralidae | RR-98-1142 | Daur |
| *Plodia interpunctella* | Pyraloidea | Pyralidae | RFD-96-1255 | Pin |
| *Gauna aegulsalis* | Pyraloidea | Pyralidae | MJM-97-0311 | Gaeg |
| *Dolichomia olinalis* | Pyraloidea | Pyralidae | AM-93-0021 | Heol |
| *Orthopygia glaucinalis* | Pyraloidea | Pyralidae | AYK-04-0881-10 | Ogla |
| *Pyralis farinalis* | Pyraloidea | Pyralidae | CWM-08-2331 | Fnls2 |
| *Pseudocossus boisduvalii* | Sesioidea | Brachodidae | DCL-07-0004 | Pbod |
| *Synechodes coniophora* | Sesioidea | Brachodidae | AxK-08-0602 | Syco |
| *Phycodes toulgoetalla** | Sesioidea | Brachodidae | DCL-07-0007 | Nggi |
| *Amauta cacica* | Sesioidea | Castniidae | DHJ-02-2377 | Amca |
| *Paysandisia archon* | Sesioidea | Castniidae | JBA-05-0069 | Pays |
| *Synemon plana* | Sesioidea | Castniidae | MJM-97-0322 | Spla |
| *Telchin licus pauperata* | Sesioidea | Castniidae | BL-08-0101 | Tlic |
| *Xanthocastnia evalthe* | Sesioidea | Castniidae | JYM-09-0104 | Xeva |

| | | | | |
|---|---|---|---|---|
| *Zegara polymorpha* | Sesioidea | Castniidae | JYM-09-0102 | Zpol |
| *Vitacea polistiformis* | Sesioidea | Sesiidae | CB-07-1005 | Vipo |
| *Melittia cucurbitae* | Sesioidea | Sesiidae | TPF-94-1106 | Mcuc2 |
| *Podosesia syringae* | Sesioidea | Sesiidae | TPF-94-1103 | Psy2 |
| *Synanthedon exitiosa* | Sesioidea | Sesiidae | JWB-06-0047 | Syex |
| *Ichneumenoptera chrysophanes* | Sesioidea | Sesiidae | AxK-08-189 | Ichr |
| *Pennisetia hylaeiformis* | Sesioidea | Sesiidae | MM-08-0001 | Phms |
| *Lophocnema eusphyra* | Sesioidea | Sesiidae | AK-11 | AK-11 |
| *Rhodoneura terminalis* | Thyridoidea | Thyrididae | 05-srnp-30764 | Lte |
| *Microsca paullula* | Thyridoidea | Thyrididae | 05-srnp-07402 | Mpll3 |
| *Pyrinioides sinuosis* | Thyridoidea | Thyrididae | AYK-04-0776-1 | Pnuo |
| *Loxiorhiza unitula* | Thyridoidea | Thyrididae | 06-srnp-41325 | TJan2 |
| *Microsca nullula* | Thyridoidea | Thyrididae | VOB-01-0008 | Tros2 |
| *Pentina flammans* | Thyridoidea | Thyrididae | 05-srnp-03426 | Pfla |
| *Striglina suzukii* | Thyridoidea | Thyrididae | AYK-06-7230 | Sski |
| *Aglaopus pyrrhata* | Thyridoidea | Thyrididae | AZ-07-2033 | Apta |
| *Glanycus insolitus* | Thyridoidea | Thyrididae | AYK-04-1279-6 | Gilt |
| *Zeuzerodes caenosa* | Thyridoidea | Thyrididae | 06-srnp-6866 | Zcae |
| *Exoncotis umbraticella* | Tineoidea | Acrolophidae | DRD-06-1317 | Exum |
| *Amydria brevipennella* | Tineoidea | Acrolophidae | DRD-01-0005 | Abre |
| *Acrolophus panamae* | Tineoidea | Acrolophidae | JWB-08-0102 | Usae |
| *Dysoptus bilobus* | Tineoidea | Arrhenophanidae | DRD-06-1320 | Dbil |
| *Peloponnesia haettenschwileri* | Tineoidea | Psychidae | DRD-01-0112 | Plha |
| *Rebelia thomanni* | Tineoidea | Psychidae | DRD-01-0110 | Rtnn |
| *Narycia duplicella* | Tineoidea | Psychidae | DRD-01-0035 | Nard |
| *Dahlica triquetrella* | Tineoidea | Psychidae | DRD-01-0041 | Dahl |
| *Oreopsyche tenella* | Tineoidea | Psychidae | DRD-01-0057 | Oreo |
| *Acanthopsyche zelleri* | Tineoidea | Psychidae | DRD-01-0043 | Azri |
| *Psyche crassiorella* | Tineoidea | Psychidae | DRD-06-1334 | Pcra |
| *Perisceptis carnivora* | Tineoidea | Psychidae | DRD-07-1301 | Carni |
| *Opogona thiadelia* | Tineoidea | Tineidae | DRD-01-0147 | Othi |

| | | | | |
|---|---|---|---|---|
| *Psecadiodes aspersus* | Tineoidea | Tineidae | DRD-01-0149 | Psasp |
| *Pyloetis mimosae* | Tineoidea | Tineidae | DRD-01-0148 | Pymi |
| *Paraptica concinerata** | Tineoidea | Tineidae | WM-08-4002 | Pcoci |
| *Dyotopasta yumaella* | Tineoidea | Tineidae | CWM-08-2534 | Dyum |
| *Oenoe hybromella* | Tineoidea | Tineidae | LS-06-0172 | Oehy |
| *Xylesthia pruniramiella* | Tineoidea | Tineidae | DRD-07-0288 | Xylp |
| *Hybroma servulella* | Tineoidea | Tineidae | DRD-07-0289 | Hybs |
| *Bathroxena heteropalpella* | Tineoidea | Tineidae | DRD-07-0290-2 | Brxh |
| *Doleromorpha porphyria* | Tineoidea | Tineidae | TH-08-6176 | Dpor |
| *Diachorisia velatella* | Tineoidea | Tineidae | TH-08-6177 | Dvel |
| *Leucomele miriamella* | Tineoidea | Tineidae | TH-08-6193 | Lmir |
| *Cephimallota chasanica* | Tineoidea | Tineidae | JCS-06-0149 | Cosa |
| *Myrmecozela ochracella* | Tineoidea | Tineidae | DRD-07-4103 | Moch |
| *Moerarchis inconcisella* | Tineoidea | Tineidae | MJM-97-0307 | Moin |
| *Xystrologa wielgusi* | Tineoidea | Tineidae | DRD-05-0263-1 | Xwi |
| *Nemapogon cloacella* | Tineoidea | Tineidae | DRD-01-0015 | Nclo |
| *Moraphaga bucephala* | Tineoidea | Tineidae | DRD-01-0144 | Mobu |
| *Scardiella approximatella* | Tineoidea | Tineidae | DRD-01-0003 | Sapp |
| *Phereoeca uterella* | Tineoidea | Tineidae | DRD-05-0252 | Putr |
| *Tinea columbariella* | Tineoidea | Tineidae | EN-91-0006 | Tco2 |
| *Trichophaga tapetzella* | Tineoidea | Tineidae | DRD-01-0131-1 | Tpgt |
| *Monopis pavlovskii* | Tineoidea | Tineidae | DRD-05-0267 | Mmon |
| *Praeacodes atomosella* | Tineoidea | Tineidae | DRD-05-0251 | Prato |
| *Tineovertex melanochryseus* | Tineoidea | Tineidae | TH-07-4001 | Tmel |
| *Heliocosma melanotypa* | Tortricoidea | Heliocosmidae | MJM-96-0274 | Hmnt |
| *Heliocosma argyroleuca* | Tortricoidea | Heliocosmidae | AZ-07-2755 | Hear |
| *Auratonota dispersans* | Tortricoidea | Tortricidae | KN-06-3410 | Audi |
| *Heppnerographa tricesimana* | Tortricoidea | Tortricidae | KN-06-1076 | Htmn |
| *Pseudatteria volcanica* | Tortricoidea | Tortricidae | 06-SRNP-2203 | Pvol |
| *Bactra furfurana* | Tortricoidea | Tortricidae | JWB-08-0103 | Bffn |
| *Bactra maiorina* | Tortricoidea | Tortricidae | NB-06-0152 | Bmra |
| *Ancylis sparulana* | Tortricoidea | Tortricidae | JBA-06-0014 | Ancy |
| *Episimus tyrius* | Tortricoidea | Tortricidae | JWB-05-0011 | Etyr |
| *Endothenia hebesana* | Tortricoidea | Tortricidae | JWB-05-0009 | Eheb |

| | | | | |
|---|---|---|---|---|
| *Epiblema abruptana* | Tortricoidea | Tortricidae | KTP-94-0521 | Basp |
| *Eucosma picrodelta* | Tortricoidea | Tortricidae | MF-06-3012 | Eucm |
| *Pelochrista zomonana* | Tortricoidea | Tortricidae | JWB-05-0037 | Eusp |
| *Spilonota eremitana* | Tortricoidea | Tortricidae | SWC-06-0227 | Serm |
| *Cryptophlebia illepida* | Tortricoidea | Tortricidae | JWB-08-0109-1 | Cppd |
| *Cydia pomonella* | Tortricoidea | Tortricidae | JG-96-0004 | Cpo |
| *Dichrorampha cancellatana** | Tortricoidea | Tortricidae | SWC-06-0226 | Dicma |
| *Grapholita delineana* | Tortricoidea | Tortricidae | AYK-04-5609 | Gdel |
| *Grapholita packardi* | Tortricoidea | Tortricidae | JWB-05-0028 | Lsna |
| *Multiquaestia purana* | Tortricoidea | Tortricidae | MF-06-3013 | Ctph |
| *Cryptaspasma querula* | Tortricoidea | Tortricidae | JBA-08-0101 | Cqur |
| *Afroploce karsholti* | Tortricoidea | Tortricidae | MF-06-3002 | Akar |
| *Hedya dimidiana* | Tortricoidea | Tortricidae | KTP-06-0125 | Hdda |
| *Lobesia aeolopa* | Tortricoidea | Tortricidae | KTP-06-0120-1 | Laelp |
| *Olethreutes fasciatana* | Tortricoidea | Tortricidae | JWB-05-0003 | Olfa |
| *Argyrotaenia alliselana* | Tortricoidea | Tortricidae | CWM-94-0262 | Arga |
| *Choristoneura rosaceana* | Tortricoidea | Tortricidae | JWB-05-0024 | Cros |
| *Clepsis melaleucanus* | Tortricoidea | Tortricidae | KTP-94-0503 | Crs |
| *Dichelia cosmopis* | Tortricoidea | Tortricidae | AZ-07-2136 | Dcsp |
| *Pandemis limitata* | Tortricoidea | Tortricidae | KTP-94-0506 | Cnsp |
| *Anacrusis nephrodes* | Tortricoidea | Tortricidae | 05-srnp-03861 | Anne |
| *Cnephasia alfacarana* | Tortricoidea | Tortricidae | JBA-08-0044 | Calfa |
| *Decodes asapheus* | Tortricoidea | Tortricidae | NB-06-0003 | Dasa |
| *Aethes promptana* | Tortricoidea | Tortricidae | JWB-05-0018 | Aesp |
| *Eugnosta sartana* | Tortricoidea | Tortricidae | JWB-06-0043 | Cstn |
| *Eugnosta busckana* | Tortricoidea | Tortricidae | NB-06-0041 | Etbk |
| *Eulia ministrana* | Tortricoidea | Tortricidae | JWB-08-0111 | Eutr |
| *Phricanthes asperana* | Tortricoidea | Tortricidae | RS-06-0101 | Pasp |
| *Amorbia humerosana* | Tortricoidea | Tortricidae | JWB-08-0095 | Ahum |
| *Platynota ideausalis* | Tortricoidea | Tortricidae | JWB-05-0029 | Pida2 |
| *Sparganothis reticulatana* | Tortricoidea | Tortricidae | JWB-06-0020 | Srtc |
| *Acleris semipurpurana* | Tortricoidea | Tortricidae | JWB-06-0031 | Cler |

| | | | | |
|---|---|---|---|---|
| *Acleris affinatana* | Tortricoidea | Tortricidae | JWB-08-0108 | Aaff |
| *Urodus decens* | Urodoidea | Urodidae | DA-03-3349 | Ursp |
| *Wockia asperipunctella* | Urodoidea | Urodidae | MM-08-0022 | Wasp |
| *Urodus parvula* | Urodoidea | Urodidae | JWB-06-0033 | Urdu |
| *Acrolepiopsis sapporensis* | Yponomeutoidea | Acrolepiidae | JCS-06-0177 | Asap |
| *Acrolepia hemiglypha* | Yponomeutoidea | Acrolepiidae | SWC-06-0204 | Dhem |
| *Acrolepia xylophragma* | Yponomeutoidea | Acrolepiidae | KN-06-1146 | Axpg |
| *Bedellia somnulentella* | Yponomeutoidea | Bedelliidae | DRD-01-0150 | Bsmu |
| *Diploschizia impigritella* | Yponomeutoidea | Glyphipterigidae | JWB-05-0010 | Dimp |
| *Doxophyrtis hydrocosma* | Yponomeutoidea | Glyphipterigidae | MM-09-1002 | Dhyd |
| *Orthotelia sparganella* | Yponomeutoidea | Glyphipterigidae | JCS-09-0112 | Ospa |
| *Proditrix gahniae* | Yponomeutoidea | Glyphipterigidae | MM-09-1003 | Pgah |
| *Lepidotarphius perornatellus* | Yponomeutoidea | Glyphipterygidae | SWC-07-2003 | Lpts |
| *Aetole tripunctella* | Yponomeutoidea | Heliodinidae | TH-08-6088 | Aetr |
| *Epicroesa metallifera* | Yponomeutoidea | Heliodinidae | LS-06-0061 | Emet |
| *Cycloplasis panicifoliella* | Yponomeutoidea | Heliodinidae | TH-08-6110 | Cpan |
| *Embola ionis* | Yponomeutoidea | Heliodinidae | TH-08-6090 | Eion |
| *Neoheliodines nyctaginella* | Yponomeutoidea | Heliodinidae | TH-08-6089 | Nnyc |
| *Leucoptera coffeella* | Yponomeutoidea | Lyonetiidae | CWM-07-2322 | Leuco |
| *Lyonetia prunifoliella* | Yponomeutoidea | Lyonetiidae | DRD-05-0253 | Lpfe |
| *Lyonetia ledi* | Yponomeutoidea | Lyonetiidae | JCS-08-1021 | Lled |
| *Plutella xylostella* | Yponomeutoidea | Plutellidae | JWB-05-0013 | Pxy |
| *Eidophasia messingiella* | Yponomeutoidea | Plutellidae | MM-08-6879 | Emnl |
| *Rhigognostis schmaltzella* | Yponomeutoidea | Plutellidae | MM-08-8908 | Rsze |
| *Argyresthia austerella* | Yponomeutoidea | Yponomeutidae | TH-08-6095 | Aase |
| *Argyresthia brockeella* | Yponomeutoidea | Yponomeutidae | JCS-08-3014 | Abck |
| *Atteva punctella* | Yponomeutoidea | Yponomeutidae | JWB-05-0041 | Atpu2 |
| *Prays fraxinella* | Yponomeutoidea | Yponomeutidae | JBA-06-0101 | Pryo |
| *Saridoscelis kodamai* | Yponomeutoidea | Yponomeutidae | SJC-08-4039 | Skod |
| *Scythropia crataegella* | Yponomeutoidea | Yponomeutidae | MM-08-6730 | Sytg |
| *Cedestis subfasciella* | Yponomeutoidea | Yponomeutidae | MM-08-5451 | Opin1 |
| *Thecobathra anas* | Yponomeutoidea | Yponomeutidae | AYK-04-5640 | Tan |
| *Yponomeuta multipunctella* | Yponomeutoidea | Yponomeutidae | JWB-06-0026 | Ymul |

| | | | | |
|---|---|---|---|---|
| *Cedestis exiguata* | Yponomeutoidea | Yponomeutidae | JCS-08-1025 | Cexi |
| *Klausius minor* | Yponomeutoidea | Yponomeutidae | JCS-08-1008 | Kmin |
| *Swammerdamia glaucella* | Yponomeutoidea | Yponomeutidae | SJC-08-3046 | Swgl |
| *Xyrosaris lichneuta* | Yponomeutoidea | Yponomeutidae | JCS-08-1006 | Xlic |
| *Yponomeuta anatolicus* | Yponomeutoidea | Yponomeutidae | JCS-08-1045 | Yana |
| *Yponomeuta kanaiellus* | Yponomeutoidea | Yponomeutidae | JCS-08-1047 | Ykan |
| *Yponomeuta myriosema* | Yponomeutoidea | Yponomeutidae | AZ-08-0503 | Yrsm |
| *Zelleria celastrusella* | Yponomeutoidea | Yponomeutidae | TH-08-6085-1 | Zcll |
| *Atemelia torquatella* | Yponomeutoidea | Yponomeutidae | MM-08-6266 | Aotl |
| *Euhyponomeutoides ribesiellus* | Yponomeutoidea | Yponomeutidae | MM-08-6137 | Eryt |
| *Paraswammerdamia conspersella* | Yponomeutoidea | Yponomeutidae | MM-08-2617 | Pcpa |
| *Ochsenheimeria urella* | Yponomeutoidea | Ypsolophidae | MM-08-0008 | Oure |
| *Ypsolopha nigrimaculata* | Yponomeutoidea | Ypsolophidae | JCS-06-0102 | Yni |
| *Bhadorcosma lonicerae* | Yponomeutoidea | Ypsolophidae | SJC-08-4041 | Bhlo |
| *Ypsolopha yasudai* | Yponomeutoidea | Ypsolophidae | JCS-08-1048 | Yyas |
| *Brachycodilla osorius* | Zygaenoidea | Aididae | MEE-98-0043 | Aios |
| *Acraga coa* | Zygaenoidea | Dalceridae | MEE-98-0040 | Acoa |
| *Dalcerides ingenita* | Zygaenoidea | Dalceridae | NM-93-0001 | Ding2 |
| *Acraga philetera* | Zygaenoidea | Dalceridae | DRD-05-0220 | Dsp |
| *Minacraga plata* | Zygaenoidea | Dalceridae | 06-srnp-22724 | Mata |
| *Epipomponia nawai* | Zygaenoidea | Epipyropidae | AYK-04-5615 | Enaw |
| *Fulgoraecia exigua* | Zygaenoidea | Epipyropidae | DM-xx-0832 | Esp2 |
| *Himantopterus fuscinervus* | Zygaenoidea | Himantopteridae | CWM-08-2535 | Laos |
| *Anticrates phaedima* | Zygaenoidea | Lacturidae | AZ-07-2775 | Anph |
| *Lactura subfervens* | Zygaenoidea | Lacturidae | 05-srnp-63415 | Lsub |
| *Eustixis aglaodora* | Zygaenoidea | Lacturidae | AZ-07-2761 | Eust |
| *Strigivenifera venata* | Zygaenoidea | Limacodidae | SEM-06-5643 | Svta |
| *Apoda biguttata* | Zygaenoidea | Limacodidae | UNK-95-1002 | Abig2 |
| *Euclea delphinii* | Zygaenoidea | Limacodidae | AM-94-0275 | Edel3 |
| *Prolimacodes badia* | Zygaenoidea | Limacodidae | AM-93-0028 | Pdbi |
| *Phobetron hipparchia* | Zygaenoidea | Limacodidae | 05-srnp-61560 | Phih |

| | | | | |
|---|---|---|---|---|
| *Pantoctenia prasina* | Zygaenoidea | Limacodidae | MF-05-0084 | Pntp |
| *Belippa horrida* | Zygaenoidea | Limacodidae | AYK-04-0989-13 | Behd |
| *Oxyplax pallivitta* | Zygaenoidea | Limacodidae | MEE-01-002 | Dmpa |
| *Doratifera quadriguttata* | Zygaenoidea | Limacodidae | MJM-97-0303 | Dqiu |
| *Hadraphe aprica* | Zygaenoidea | Limacodidae | SEM-06-5637 | Hapri |
| *Isochaetes beutenmuelleri* | Zygaenoidea | Limacodidae | CWM-94-0298 | Isoc |
| *Perola murina* | Zygaenoidea | Limacodidae | MEE-98-0078 | Pmur |
| *Megalopyge crispata* | Zygaenoidea | Megalopygidae | UNK-95-1001 | Lcr2 |
| *Mesoscia dyari* | Zygaenoidea | Megalopygidae | MEE-05-0050 | Mdyi |
| *Megalopyge lapena* | Zygaenoidea | Megalopygidae | MEE-05-0004 | Mglp |
| *Norape tener* | Zygaenoidea | Megalopygidae | MEE-05-0020 | Nora |
| *Podalia contigua* | Zygaenoidea | Megalopygidae | 06-SRNP-43001 | Octg |
| *Eterusia aedea* | Zygaenoidea | Zygaenidae | AYK-04-0788-04 | Etsp |
| *Pidorus glaucopis* | Zygaenoidea | Zygaenidae | AYK-04-0843-03 | Pigl |
| *Acoloithusfalsarius* | Zygaenoidea | Zygaenidae | AZ-06-0001 | Acfa |
| *Malthaca dimidiata* | Zygaenoidea | Zygaenidae | JWB-08-0099 | Pydd |
| *Pryeria sinica* | Zygaenoidea | Zygaenidae | JWB-05-0045 | Psin |
| *Zygaena fausta* | Zygaenoidea | Zygaenidae | JBA-06-0024 | Zgfa |
| *Actias isis* | Bombycoidea | Saturniidae | Gen bank number: EU032775.1 | |
| *Actias luna* | Bombycoidea | Saturniidae | Gen bank number: AF015048.1 | |
| *Actias selene* | Bombycoidea | Saturniidae | Gen bank number: AF373954.1 | |
| *Anisota stigma* | Bombycoidea | Saturniidae | Gen bank number: EU032774.1 | |
| *Antheraea godmani* | Bombycoidea | Saturniidae | Gen bank number: AY301299.1 | |
| *Antheraea helferi* | Bombycoidea | Saturniidae | Gen bank number: AAY301300.1 | |
| *Antheraea jana* | Bombycoidea | Saturniidae | Gen bank number: AY301304.1 | |
| *Antheraea kelimutuensis* | Bombycoidea | Saturniidae | Gen bank number: AY301303.2 | |
| *Antheraea lampei* | Bombycoidea | Saturniidae | Gen bank number: AY301301.1 | |
| *Antheraea larissa* | Bombycoidea | Saturniidae | Gen bank number: AY301302.1 | |
| *Antheraea paphia* | Bombycoidea | Saturniidae | Gen bank number: AF373952.2 | |
| *Antheraea pernyi* | Bombycoidea | Saturniidae | Gen bank number: AY301310.2 | |

| | | | |
|---|---|---|---|
| *Antheraea raffrayi* | Bombycoidea | Saturniidae | Gen bank number: AY301307.1 |
| *Antheraea rosemariae* | Bombycoidea | Saturniidae | Gen bank number: AY301308.1 |
| *Antheraea roylii* | Bombycoidea | Saturniidae | Gen bank number: AY301309.1 |
| *Antheraea yamamai* | Bombycoidea | Saturniidae | Gen bank number: MW677190.1 |
| *Antheraea youngi* | Bombycoidea | Saturniidae | Gen bank number: AY301311.1 |
| *Antherina suraka* | Bombycoidea | Saturniidae | Gen bank number: AF373955.2 |
| *Archaeoattacus edwardsii* | Bombycoidea | Saturniidae | Gen bank number: AF015046.1 |
| *Argema mimosae* | Bombycoidea | Saturniidae | Gen bank number: AF373951.1 |
| *Attacus atlas* | Bombycoidea | Saturniidae | Gen bank number: EU032768.1 |
| *Attacus caesar* | Bombycoidea | Saturniidae | Gen bank number: AF373948.1 |
| *Attacus lorquinii* | Bombycoidea | Saturniidae | Gen bank number: AF373950.1 |
| *Bunaea alcinoe* | Bombycoidea | Saturniidae | Gen bank number: EU032888.1 |
| *Callosamia angulifera* | Bombycoidea | Saturniidae | Gen bank number: AF015050.1 |
| *Callosamia promethea* | Bombycoidea | Saturniidae | Gen bank number: AF015052.1 |
| *Callosamia securifera* | Bombycoidea | Saturniidae | Gen bank number: AF015053.1 |
| *Ceranchia apollina* | Bombycoidea | Saturniidae | Gen bank number: AF373958.1 |
| *Cirina forda* | Bombycoidea | Saturniidae | Gen bank number: AF373959.1 |
| *Citheronia sepulcralis* | Bombycoidea | Saturniidae | Gen bank number: EU032803.1 |
| *Copaxa multifenestrata* | Bombycoidea | Saturniidae | Gen bank number: EU032799.1 |
| *Copiopteryx semiramis* | Bombycoidea | Saturniidae | Gen bank number: EU032802.1 |
| *Coscinocera hercules* | Bombycoidea | Saturniidae | Gen bank number: AF015051.1 |
| *Cricula elaezia* | Bombycoidea | Saturniidae | Gen bank number: EU032795.1 |
| *Dysdaemonia boreas* | Bombycoidea | Saturniidae | Gen bank number: EU032806.1 |
| *Eacles imperialis* | Bombycoidea | Saturniidae | Gen bank number: AF015058.1 |
| *Epiphora mythimnia* | Bombycoidea | Saturniidae | Gen bank number: AF015055.1 |
| *Erythromeris flexilineata* | Bombycoidea | Saturniidae | Gen bank number: EU032808.1 |
| *Eubergia caisa* | Bombycoidea | Saturniidae | Gen bank number: EU032807.1 |
| *Eupackardia calleta* | Bombycoidea | Saturniidae | Gen bank number: AF015054.1 |
| *Graellsia isabellae* | Bombycoidea | Saturniidae | Gen bank number: AF015047.1 |
| *Heniocha apollonia* | Bombycoidea | Saturniidae | Gen bank number: EU032816.1 |

| | | | |
|---|---|---|---|
| *Holocerina smilax* | Bombycoidea | Saturniidae | Gen bank number: AF373963.1 |
| *Hyalophora cecropia* | Bombycoidea | Saturniidae | Gen bank number: EU032818.1 |
| *Hyalophora euryalus* | Bombycoidea | Saturniidae | Gen bank number: AF015057.1 |
| *Hyalophora gloveri* | Bombycoidea | Saturniidae | Gen bank number: EU032809.2 |
| *Gonimbrasia macrothyris* | Bombycoidea | Saturniidae | Gen bank number: AF373965.1 |
| *Gonimbrasia tyrrhea* | Bombycoidea | Saturniidae | Gen bank number: AF373966.1 |
| *Loepa sikkima* | Bombycoidea | Saturniidae | Gen bank number: AF015059.1 |
| *Lonomia achelous* | Bombycoidea | Saturniidae | Gen bank number: EU032831.1 |
| *Opodiphthera eucalypti* | Bombycoidea | Saturniidae | Gen bank number: EU032847.1 |
| *Othorene verana* | Bombycoidea | Saturniidae | Gen bank number: EU032852.1 |
| *Oxytenis albilunulata* | Bombycoidea | Saturniidae | Gen bank number: EU032845.1 |
| *Polythysana apollina* | Bombycoidea | Saturniidae | Gen bank number: EU032854.1 |
| *Pseudimbrasia deyrollei* | Bombycoidea | Saturniidae | Gen bank number: EU032860.1 |
| *Rhescyntis hippodamia* | Bombycoidea | Saturniidae | Gen bank number: EU032871.1 |
| *Rhodinia fugax* | Bombycoidea | Saturniidae | Gen bank number: AF015061.1 |
| *Rothschildia forbesi* | Bombycoidea | Saturniidae | Gen bank number: AF015060.1 |
| *Rothschildia orizaba* | Bombycoidea | Saturniidae | Gen bank number: AF015062.1 |
| *Samia cynthia* | Bombycoidea | Saturniidae | Gen bank number: AF015063.1 |
| *Samia luzonica* | Bombycoidea | Saturniidae | Gen bank number: AF373972.1 |
| *Samia ricini* | Bombycoidea | Saturniidae | Gen bank number: AF015065.1 |
| *Saturnia albofasciata* | Bombycoidea | Saturniidae | Gen bank number: AF373968.1 |
| *Saturnia anona* | Bombycoidea | Saturniidae | Gen bank number: AF373969.1 |
| *Saturnia caecigena* | Bombycoidea | Saturniidae | Gen bank number: AF373970.1 |
| *Saturnia galbina* | Bombycoidea | Saturniidae | Gen bank number: AF373971.1 |
| *Saturnia mendocino* | Bombycoidea | Saturniidae | Gen bank number: AF373973.1 |
| *Saturnia walterorum* | Bombycoidea | Saturniidae | Gen bank number: AF373975.1 |
| *Titaea tamerlan* | Bombycoidea | Saturniidae | Gen bank number: EU032887.1 |
| *Vegetia ducalis* | Bombycoidea | Saturniidae | Gen bank number: EU032889.1 |

| **Table S3.** list of 27 genes used for phylogenetic inference in this study. Fragment length (bp) are also provided. | | |
|---|---|---|
| Gene number | mRNA coding gene name | Number of base pairs |
| 1 | Syntaxin 1A | 488 |
| 2 | Phosphogluconate degydrogenase | 733 |
| 3 | GTP binding protein | 797 |
| 4 | Glucosamine phosphate isomerase | 557 |
| 5 | Clathrin coat assembly protein | 626 |
| 6 | Gelosolin | 650 |
| 7 | Glycogen synthase | 974 |
| 8 | Glu - + pro-tRNA synthase | 401 |
| 9 | Triose phosphate isomerase | 458 |
| 10 | Proteaosome subunit | 545 |
| 11 | his tRNA synthase | 455 |
| 12 | Deaminase | 770 |
| 13 | hypothetical protein* (unknown) | 455 |
| 14 | Dynamin | 221 |
| 15 | Phosphate dehydrogenase | 623 |
| 16 | Tetrahydropholate synthase | 593 |
| 17 | Methyletransferase | 737 |
| 18 | ala-tRNA synthase | 710 |
| 19 | Nucleolar cistine rich protein | 323 |
| 20 | Glucose phosphate isomerase | 665 |
| 21 | Acetyl-CoA carboxylase | 506 |
| 22 | Spectrin alpha chain | 593 |
| 23 | Carbamolyphosphate synthase | 3056 |
| 24 | Dopa decarboxylase | 1340 |
| 25 | Enoclase | 1139 |
| 26 | Period | 1433 |
| 27 | Wingless | 455 |

| **Table S4.** Overview and links to the image databases used for this study. | | |
|---|---|---|
| **Database name** | **URL** | **Description** |
| Afromoths | http://www.afromoths.net | African moth database |
| Bold V3 & Bold V4 | V3: https://v3.boldsystems.org/<br>V4: https://v4.boldsystems.org/ | Multispecies database |
| Butterflies of America | http://www.butterfliesofamerica.com | Database on the butterflies of America (Mostly USA) |
| Cornell University insect collection | https://cuic.entomology.cornell.edu/ | University collection Lepidoptera database |
| CSIRO moths of Australia | https://moths.csiro.au/ | Australian Lepidoptera image database |
| GBIF (Global Biodiversity Image Facility) | https://www.gbif.org/ | Multi species database |
| Gracilliaridae database | http://www.gracillariidae.net | Database specifically for the Superfamily Gracillarioidea |
| Insecta Pro | http://insecta.pro/ | Insect image database |
| Japanese moths online | http://www.jpmoth.org | Image database of the moths of Japan |
| Lepi forum | https://lepiforum.org/ | A German Lepidoptera forum and database with images |
| Moths of india | https://www.mothsofindia.org/ | Image database of the moths of India |
| Moth photographers of America | https://mothphotographersgroup.msstate.edu/ | Image database of the moths of North America (Mostly USA). A database part of Mississippi state University by the Mississippi entomological museum |

| | | |
|---|---|---|
| Museum of Shimane | http://museum-database.shimane-u.ac.jp/specimen/detail/502420180331122106 | Museum of Shimane (Japan) moth image database |
| Scan bugs | http//:scan-bugs.org/portal/index.php. | Insect image database |
| Swedish museum of natural history | http://www2.nrm.se/en/ | Insect database of the Swedish museum of natural history |
| Smithsonian tropical institute institute | Smithsonian Tropical Research Institute \| (si.edu) | Insect database of the Smithsonian institute USA |
| Taiwan encyclopedia of life | https://taieol.tw/ | Taiwanese biodiversity encyclopedia website |
| Project Biodiversity Belgium | Projectbiodiversity.be | Biodiversity recording project in Belgium |

**Inference of ancestral eyespot origins: validating the model-comparison approach**

We used a model comparison approach to evaluate support for ancestral eyespot origin in multiple subclades across our Ditrysian phylogeny. In this approach, we specified two alternative models for each subclade, which only differed from each other in the priors assigned to the root state frequencies – i.e. the presence or absence of eyespots in the common ancestor of the subclade. In the 'common ancestor' model, we enforced a 100% probability of eyespot presence at the root of the tree, whereas in the 'multiple origins' model, we enforced the ancestral absence of eyespots, such that, eyespots in the most ancestrally diverged lineages of the subclade must have originated independently. For each subclade, we then compared the absolute fit of these alternative models to the data by estimating their marginal likelihood and computing the $\log_{10}$ Bayes Factor (BF) between the models. Following convention, we interpret $\log_{10}$(BF) values between 0-0.5 as barely worth mentioning, values 0.5-1 as substantial evidence, values between 1-2 as strong evidence, and values > 2 as decisive evidence in favour of the 'common ancestor' model, if positive, and in favour of the 'multiple origins' model if negative.

We used simulations to evaluate the performance of this model-comparison approach, specifically to estimate the rate of false positives (Type-I error), where a subclade without ancestral eyespots would receive a positive $\log_{10}$(BF), and the rate of false negatives (Type-II error), where we would compute a negative $\log_{10}$(BF) for a subclade which actually did have an eyespot-bearing common ancestor. We simulated 500 trees, with 10, 50, and 100 terminal nodes, using a birth-death process with speciation ($\lambda = 0.01$) and extinction ($\mu = 0.05$) rates within the range for Lepidoptera (Wahlberg et al. 2013). We simulated trees using the *TreeSim* package (Stadler, 2019) in R (R core team, 2023). For each tree, we then simulated two-character histories for a discrete trait, one where the trait was present at the root of the tree and one where it was absent. Character histories were simulated using the *simulate_mk_model* function in the R package *castor* (Louca and Doebeli, 2017). Transition rates were based on the posterior means of our clade-wide analysis of eyespot evolution (see "*Modelling the origin and loss of eyespots in Lepidoptera*" in the main Methods), with values of 0.002 for the gain of the trait and 0.015 for the loss of the simulated trait. Thus, like eyespots across Ditrysia, our simulated traits are lost at a rate almost an order of magnitude higher than they are gained.

For each of the 3,000 tree and character simulated datasets (two-character datasets per each of 1,500 trees), we applied the model-comparison approach used for empirical datasets and outlined above. We expected $\log_{10}$(BF) to be positive for character histories with ancestral trait presence and negative for character histories simulated with trait absence at the root. We also expected that the distribution of $\log_{10}$(BF) values and their false positive and false negative error rates would depend on the size of the tree.

Overall, we found our model-comparison approach to be conservative (Fig S10). False positives, where the true ancestral state was trait absence but the $\log_{10}$(BF) was positive, occurred in 2.2%, 1.4%, and 1.2% of simulations on large (100 tips), medium-size (50 tips), and small (10 tips) trees, respectively. False positives incorrectly supported by $\log_{10}$(BF) > 0.5 (substantial, strong, or decisive evidence) accounted for 0.8% of simulations on large trees and no simulations on medium-size or small trees.

False negatives, where the true ancestral state is trait presence but the $\log_{10}$(BF) is negative, were slightly more common: 4.4% in large trees, 4.4% in medium-size trees and 5.2% in small trees. This discrepancy is owed to the unequal rates of character gain and loss. A relatively high rate of losses erodes the phylogenetic signal that otherwise enables reconstructing the ancestral presence of the trait. Additional simulations where both trait gains and losses had slow rates, comparable to the estimated rate of eyespot gains (0.002) resulted in comparable rates of false positives and false negatives.

Despite the error rates being near or below what is typically considered acceptable (i.e. 5%), we noted that the $\log_{10}$(BF) tended to be modest (Fig. S26). For small trees with 10 terminal taxa and ancestral eyespots, the strength of evidence supporting the correct inference was unlikely to surpass the category of "barely worth mentioning" (mean $\log_{10}$(BF) = 0.19, 95% HPD interval = -0.03 – 0.42). For medium and large trees, the expected support was "substantial" (mean $\log_{10}$(BF) = 0.74, 95% HPD interval = -0.02 – 1.53) and "strong" (mean $\log_{10}$(BF) = 1.18, 95% HPD interval = -0.03 – 2.26) respectively.

Taken together, our simulation analysis suggests that inferred ancestral eyespots in Saturniidae and Nymphalidae, even if supported by modest $\log_{10}$(BF) values, are likely correct, based on the estimated error rate of ~5% in large trees and medium-size trees. However, there might be clades within Ditrysia where the phylogenetic signature of ancestral eyespot could have been erased by a relatively rapid evolutionary losses of the trait. We thus consider the results of our model-comparison a conservative estimate of the evolutionary depth of ancestral eyespots currently found in saturniids, nymphalids and their relatives.

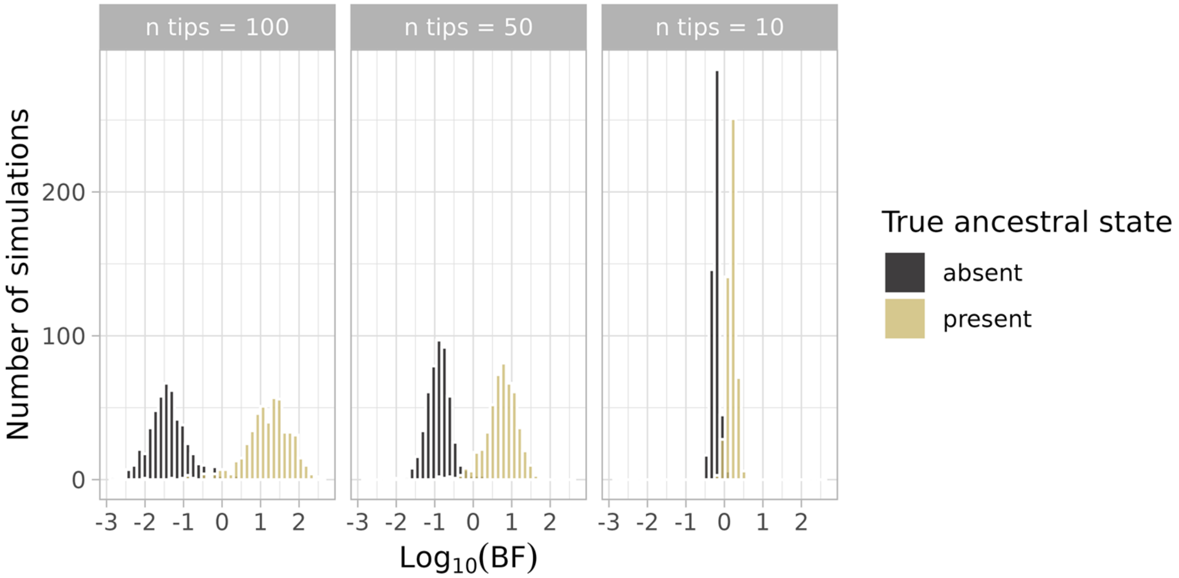


**Figure S13.** Support for the 'common ancestor' model in simulated character histories in large (n tips = 100), medium-size (n tips = 50) and small (n tips = 10) trees. Positive $\log_{10}$(BF) values correctly indicate support for the 'common ancestor' model when the true ancestral state is the presence of the trait. Negative $\log_{10}$(BF) values are expected when the true ancestral state is absence, thus favouring the 'multiple origins' model. Character histories were simulated under ancestral eyespot presence and ancestral eyespot absence for 500 trees of each size.

**Table S5.** Log 10 of the raw Bayes factors ($\log_{10}$ BF) of comparisons between the 'common ancestor' and the 'multiple-origins' models of eyespot evolution for all eyespots relaxed only. Values can vary from -1 to +1. Values can vary from -1 to +1. For each subclade with two or more eyespot-bearing taxa, we estimated the marginal likelihood of each of these alternative models and compared them via $\log_{10}$ BF. We report log10 BF values for comparisons between the 'common ancestor' and 'multiple origins' models. Following our estimation of run-to-run uncertainty in marginal likelihoods (see Methods), we treat log10 BF values < 0.5 as weak or inconclusive support and refrain from strong biological interpretation of such values. Here, we show subclades with weak but positive evidence for the 'common ancestor' model ($\log_{10}$ BF between 0.1 – 0.5), and Bayes Factors based on marginal likelihood estimates using the stepping stone algorithm. All other subclades examined, either had support for the 'multiple origins' model ($\log_{10}$ BF < -0.1) or showed no positive evidence for either model ($\log_{10}$ BF between -0.1 – 0.1).

| Clade | Eyespot type | $\log_{10}$ BF for common ancestor |
|---|---|---|
| Saturniidae | All relaxed | 0.150 |
| Nymphalidae | All relaxed | 0.520 |

Tree topology

Tineoidea (Latreille, 1810)

Tineoidea is a superfamily under debate and if all species assigned to the Tineoidea are included, then this superfamily is polyphyletic (Mitter et al. 2017, Figs. 3, S14). Previous works have demonstrated that if certain taxa are removed (Mutanen et al. 2010; Regier et al. 2015A) the Tineoidea become a monophyletic group (Mutanen et al. 2010; Regier et al. 2015A; Mitter et al. 2017).

The Tineoidea in this study are the most basal Ditrysian group, this is confirmed by previous works and our own (Heikkilä et al. 2015; Mitter et al. 2017; Kawahara et al. 2019; Figs. 3, S14). This work does not attempt to resolve the Tineoidea. We did not attempt to remove any species and as such this superfamily remains polyphyletic in this work (Fig. S14). The potential reclassification of this superfamily into various taxonomic rankings/groups is worthy of investigation in potential future works. Many new discoveries are likely to be found in this basal group.

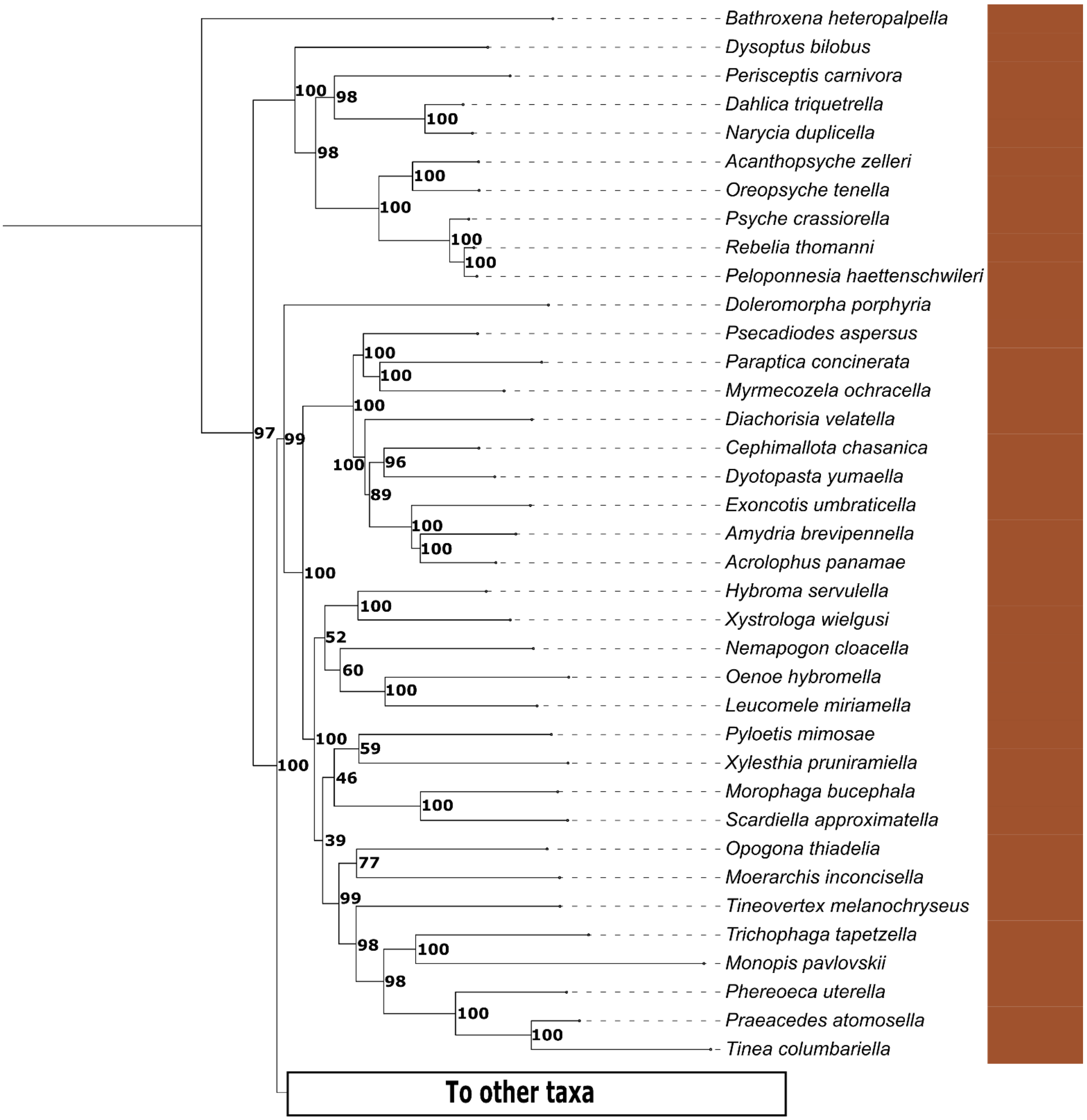

**Figure S14. Phylogenetic view of the Tineoidea only.**

Yponomeutoidea (Stephens, 1829)

Historically, the superfamily Yponomeutoidea continues to be under debate (Mitter et al. 2017) with certain species not clustering together. However, one molecular study did demonstrate support for the group's monophyletic status (Sohn et al. 2013). The present study provides some support for the monophyly of Yponomeutoidea (Figs. 3, S15). However, this monophyly is only achieved if we exclude taxa from the genus *Leucoptera* (Yponomeutoidea, Lyonetiidae, Lyonetiinae) and the genus *Cycloplasis* (Yponomeutoidea, Heliodinidae), which in this study is nested within a clade of Gracillarioidea (*Leucoptera*, Fig. S16) or are seen clustering with Urodoidea (*Cycloplasis*, Fig. S17). Further investigations are needed to determine the phylogenetic placement of these taxa as our findings are not as in previous works such as Sohn et al. (2013).

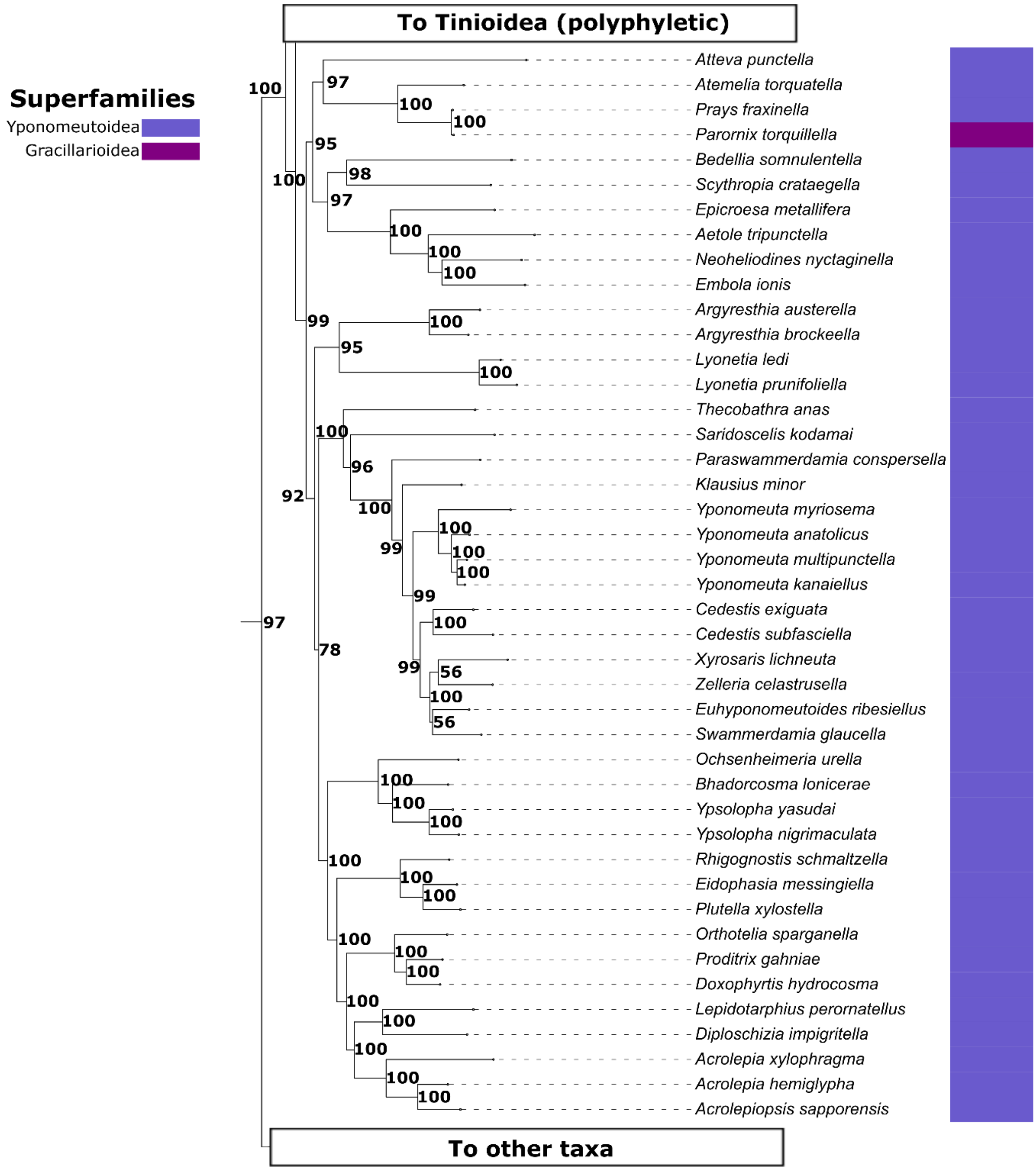


**Figure S15. Phylogenetic view of the Yponomeutoidea only.** The species *Parornix torquillella*, (Gracillariidae, Gracillariinae) is also observed herein.

Gracillarioidea (Stainton, 1854)

Previous works placed the subfamily Douglasiidae in the superfamily Gracillarioidea (Kristensen and Schmidt-Rhaesa, 1998). However, more recent phylogenetic research has changed this placement. It was found that the Douglassidae does not belong within the Gracillarioidea (Regier et al. 2009; Mutanen et al. 2010; Kawahara et al. 2011; Regier et al. 2013, Figs. 3, S16) and are covered in more detail below.

The species (*Parornix torquillella*, Gracillariidae, Gracillariinae) in this study is moved basal within the Yponomeutoidea (Figs. 3, S15, S16). This finding is also further explored below. As suggested by Mitter et al. (2017), we find that the Gracillarioidea and the Yponomeutoidea do not form a monophyletic group but instead are adjacent to each other (Figs. 3, S15, S16).

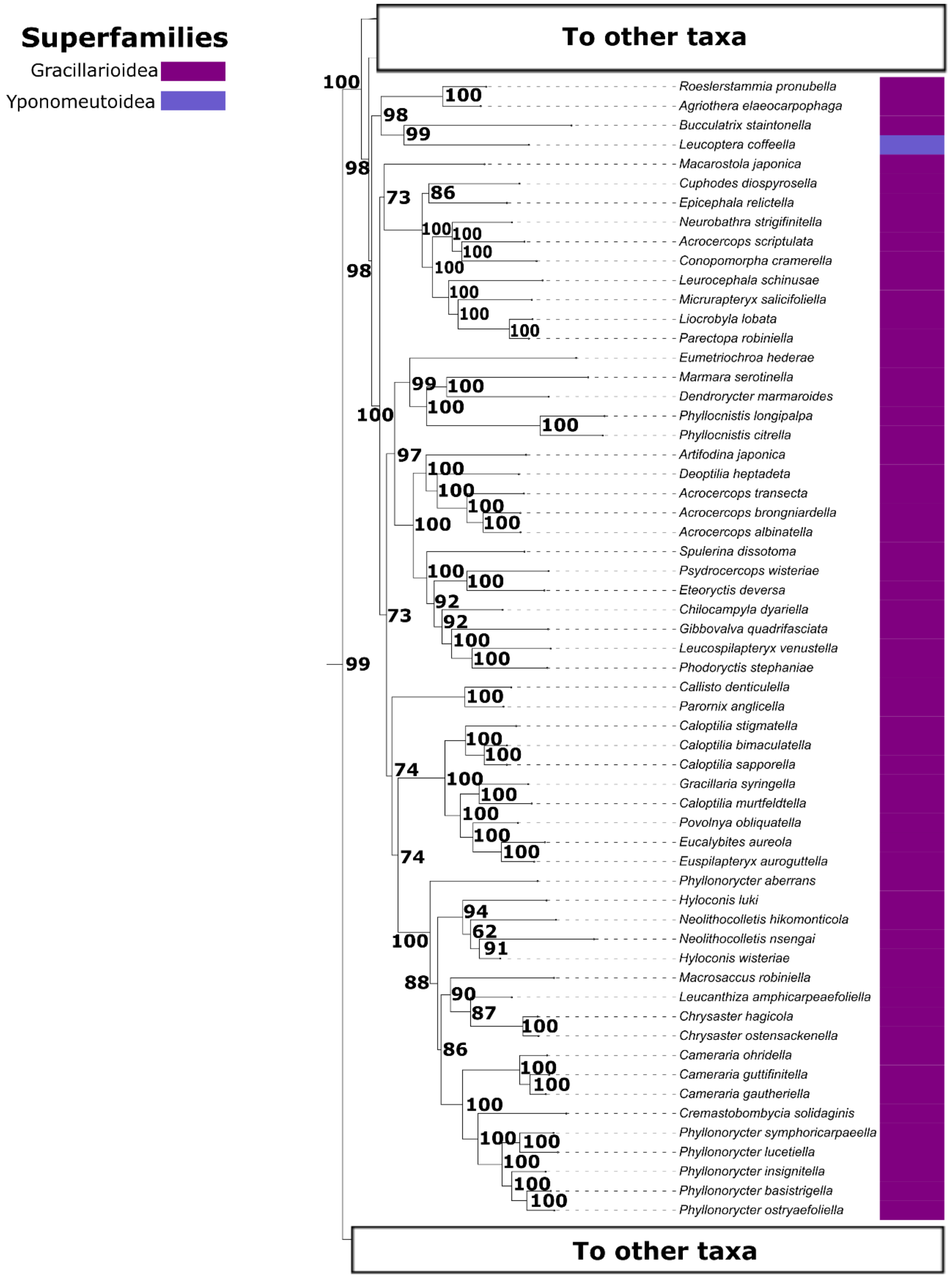


**Figure S16. Phylogenetic view of the Gracillarioidea only.** The species *Leucoptera coffeella* is also present.

Choreutoidea (Stainton, 1858), Pterophoroidea (Latreille, 1802) and Urodoidea (Kyrki, 1988)
These three superfamilies are represented by very few individuals in this work (Figs. 3, S17). However, despite the small number of species, we find that the Urodoidea are monophyletic. The Pterophoroid species *Emmelina monodactyla* sister to the Douglassidae (Figs. 3, S17). We note that one species of Choreutoidea (*Milleria dolosalis*) clusters with the Yupometoid *Cycloplasis panicifoliella* and the Urodoidea (Figs. 3, S17). The superfamilies Urodoidea and Choreutoidea have been recovered as sister before by Rota et al. (2022) however this was without Pterophoroidea, the Douglassidae or *C. panicifoliella*. Further research is required before any conclusions can be made from this grouping (Figs. 3, S17).

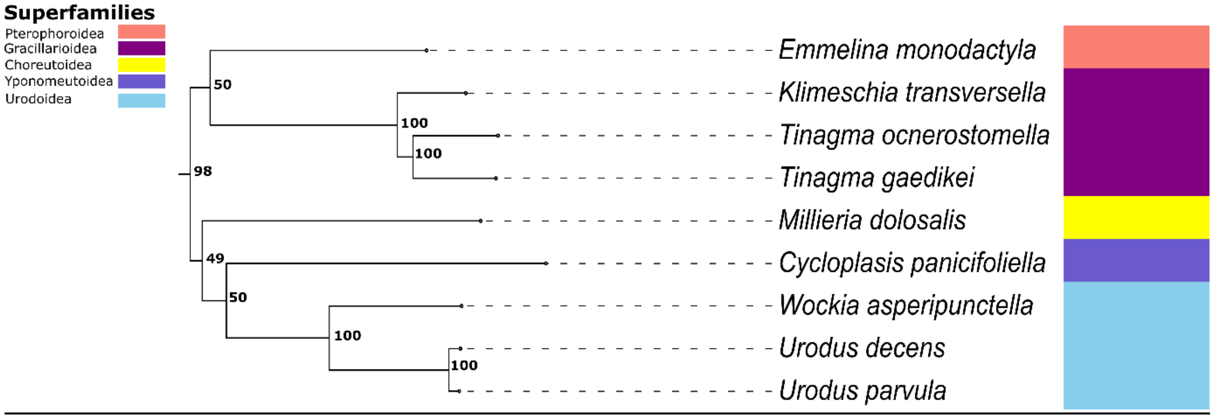


**Figure S17. Phylogenetic view of the Choreutoidea, Pterophoroidea and Urodoidea.** The Douglassidae and the yponomeutid moth *C. panicifoliella* are also present.

Tortricoidea (Latreille, 1802)
In comparison to other groups of Lepidoptera, there has been comparatively little work conducted on the Tortricoidea (Mitter et al. 2017). In this study, we demonstrate that the Tortricoidea could potentially be a monophyletic superfamily pending future research. However, the reclassification of two species from the genus *Heliocosma* is required. In our study the genus *Heliocosma* is found forming a small, separate group alongside one species from the superfamily Galacticoidea and one species of Cossioidea (Figs. 3, S18). Within the remaining Tortricoidea, we observe that our findings backed by strong bootstrap values (77-100 Fig. S19) are supported by a multigene study (Reiger et al. 2012A).

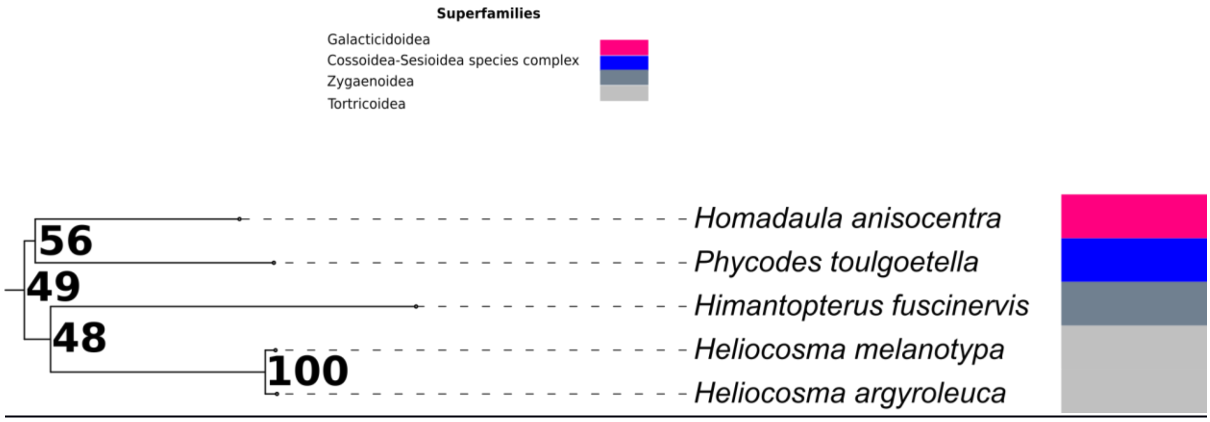


**Figure S18. *Heliocosma sp.,* the Galactid *Homadaula anisocentra* and the Cossid *Phycodes toulgoetella* only.**

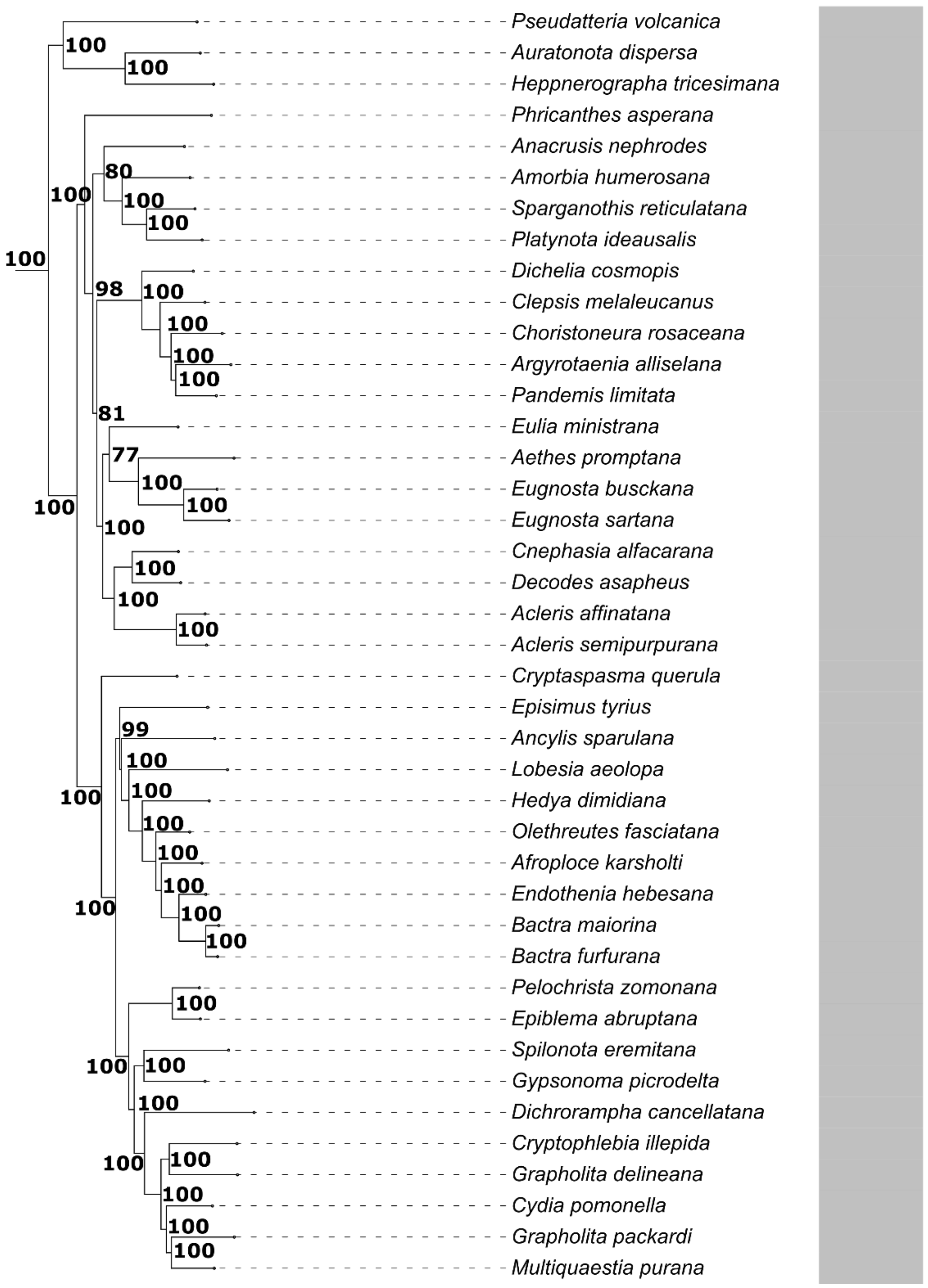


**Figure S19. Phylogenetic view of the Tortricoidea only.**

<u>Gelechioidea (Stainton, 1854)</u>

The Gelechioidea in this study are considered a monophyletic group. This is demonstrated by previous studies (Heikkilä et al. 2014; Heikkilä et al. 2015; Sohn et al. 2015; Fig. 3, Fig. S20). We observe the Batrachedridae, Blastobasidae, Coleophoridae, and Elachistidae (restricted to the subfamily Elachistinae), together with representatives of Gelechiidae (subfamilies Dichomeridinae and Gelechiinae, including Aristotelia, Dichomeris, Encolapta, Faristenia, Friseria, Hypatima, Monochroa, Pectinophora, Teleiodes, Aroga, and Caryocolum). The smaller families Momphidae, Scythrididae, and Stathmopodidae also belong to this first assemblage.

The Oecophorid-line comprises the families Autostichidae, Chimabachidae, Cosmopterigidae (Euclemensia, Pancalia, Pyroderces, Hyposmocoma), Depressariidae (Agonopterix, Bibarrambla, Psilocorsis, Thudaca, Aeolanthes), and Elachistidae (subfamilies Ethmiinae, Hypertrophinae, and Stenomatinae), together with Lecithoceridae, Lypusidae, Oecophoridae, Peleopodidae, and Xyloryctidae (69 – 100, Fig. S20). Globally, it appears that the majority of the

Gelechioidea as a group remains unresolved (Mitter et al. 2017) and may yet be subject to systematic changes in the future.

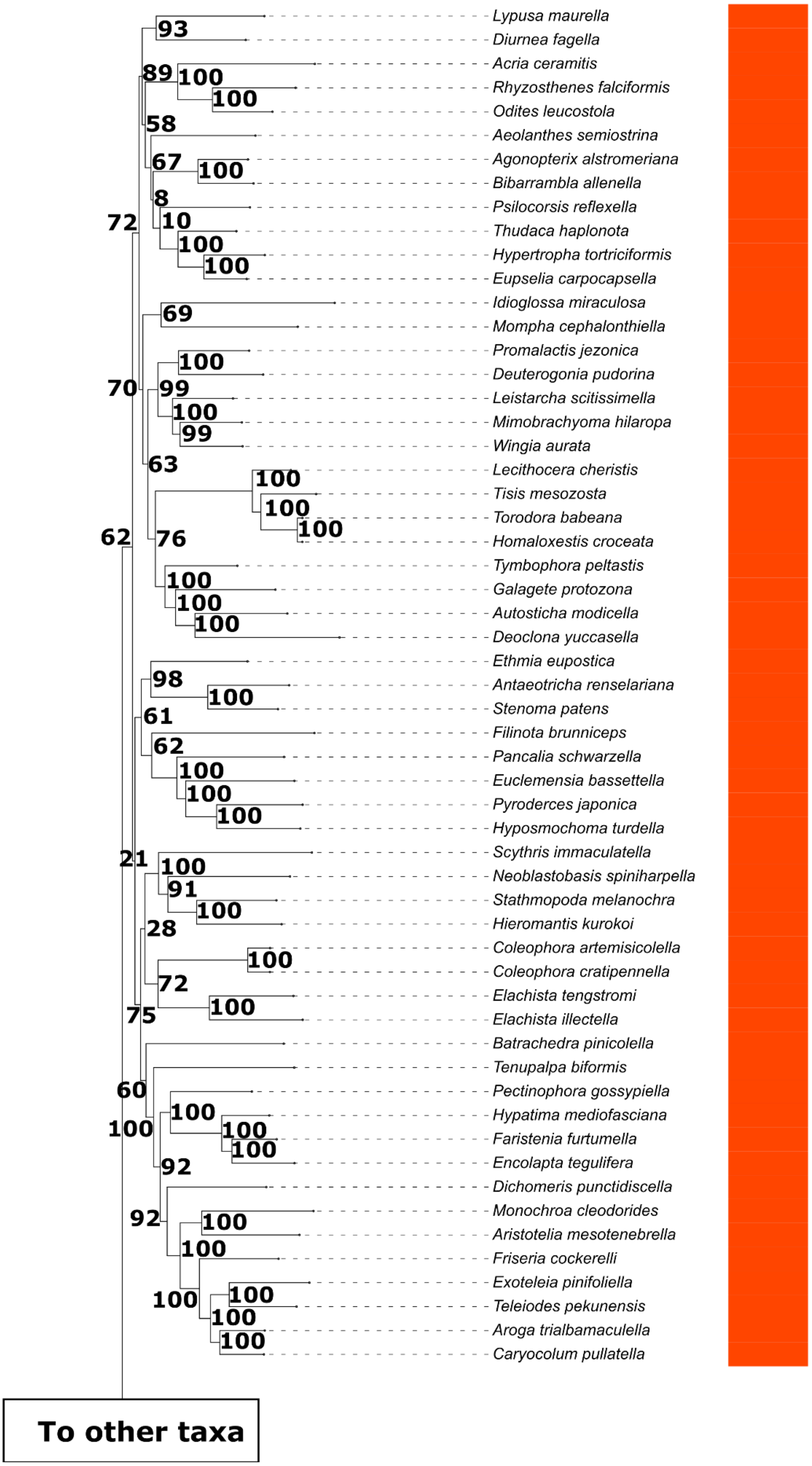


**Figure S20. Phylogenetic view of the Gelechioidea only.**

Cossioidea (Leach, 1815), Sesioidea (Boisduval, 1828), Zyganoidea (Latreille, 1809) and Choreutoidea (Stainton, 1858):

This group (Cossioidea, Sesioidea, Zyganoidea and Choreutoidea) of superfamilies have been the subject of intense debate for many years. Certain taxa from all four superfamilies appear distributed across the Lepidoptera meaning that until a revision is conducted, these groups remain paraphyletic, an observation previously mentioned by multiple authors (Mutanen et al. 2010; Bazinet et al. 2013; Regier et al. 2013; Heikkilä et al. 2015; Mitter et al. 2017).

Firstly, in this study we find that the Cossioidea and Sesioidea superfamilies are highly paraphyletic, forming two assemblages. The main assemblage of species from both superfamilies is formed (sometimes referred to as a species complex, Fig. S21, Cossioidea in green, Sesioidea in red, also see Heikkilä et al. 2015; Mitter et al. 2017). Secondly, in this study we find a separate second assemblage of only Sesioidea, this second assemblage splits off earlier and is found as sister to the majority of the Choreutoidea (Fig. S21).

Unlike some previous studies (Heikkilä et al. 2015; Mitter et al. 2017) we do not find any taxa associated with the Zyganoidea to be included within the Cossioidea-Sesioidea species complex (S21, S22) and instead we find that the Zyganoidea are placed next to the main Cossioidea-Sesioidea group but as their own entity (S21, S22). Based on our analysis, we have moderate to strong support (bootstrap values of 48 - 100) for the exclusion of the genera *Himanopterus, Fulgoraecia and Epipomponia* (currently Zyganoidea) from the Cossioidea-Sesioidea species complex. Instead, the species *Fulgoraecia & Epipomponia* alongside the Immoidea are placed as sister to all Cossioidea-Sesioidea, Choreutoidea and the remaining bulk of the Zyganoidea (Fig. S21). While *Himanopterus,* clusters with the Genus *Helicosoma*, the Cossid *Phycodes toulgoetella* and the Galacticoidea*,* Fig. S18). More work remains to determine the phylogenetic placement between the different species which comprise the Cossioidea-Sesioidea species complex. This could be considered a minor improvement as we demonstrate (in support of Mitter et al. 2017) that there is good reason to reclassify the Cossioidea-Sesioidea at the superfamily level.

As mentioned above, the large majority of Choreutoidea (excluding *Milleria dolosalis*, Figs. S17, S21) are also found here as sister to the smaller group of Sesioidea which split off before all Cossioidea, the remaining Sesioidea and the Zyganoidea (Fig. S22). The placement of the Choreutoidea in this phylogenetic position is not entirely novel. Both Brock (1971) as well as Heppner and Duckworth (1981) had originally theorised based on morphology that Choreutoidea were part of the Sesioidea. Here we maintain that Choreutoidea remain separate taxonomically at the level of the superfamily however the placement of the bulk of the Choreutoidea as sister to a group of Sesioidea is intriguing and warrants further study.

Finally, the superfamily Zyganoidea in this study (Figs. 3, S22) is composed of two subclades, one including the families Limacodidae, Aididae and Megalopygidae, and the other including most taxa in the families Lacturidae, Dalceridae, and Zygaenidae (Epstein, 1996; Mutanen et al. 2010; Bazinet et al. 2013; Regier et al. 2013; Mitter et al. 2017 Figs. 3, S22). As mentioned above, species from the families Epipyropidae and Himantopteridae cluster outside the Zyganoidea, instead clustering with either Immoidea (Fig. S21) or with the small group compromising the Helicosoma, Galacticoidea and the Cossid *Phycodes toulgoetella* (Himantopteridae, Fig. S18). We suggest that increased genetic and taxonomic sampling of the Immoidea and Galacticoidea clades coupled with more studies looking at the Zyganoidea.

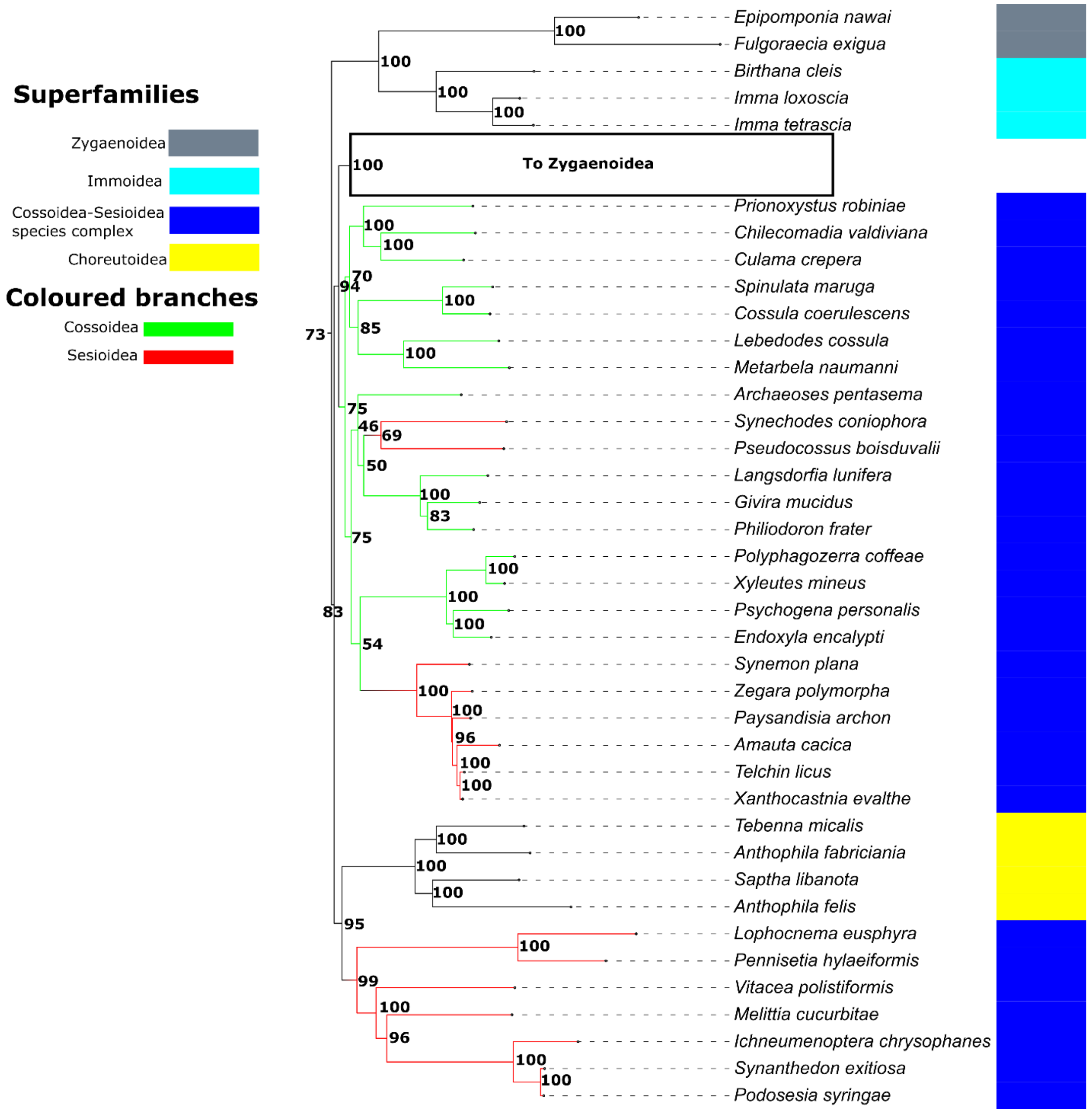


**Figure S21. Phylogenetic view of the Cossioidea and Sesioidea.** Branches of this sub tree have been colour coded Green (Cossioidea) and Red (Sesioidea). The bulk of the Choreutoidea as well as the Immoidea and the Zyganids *Epipomponia nawai* and *Fulgoraecia exigua* are also observed here.

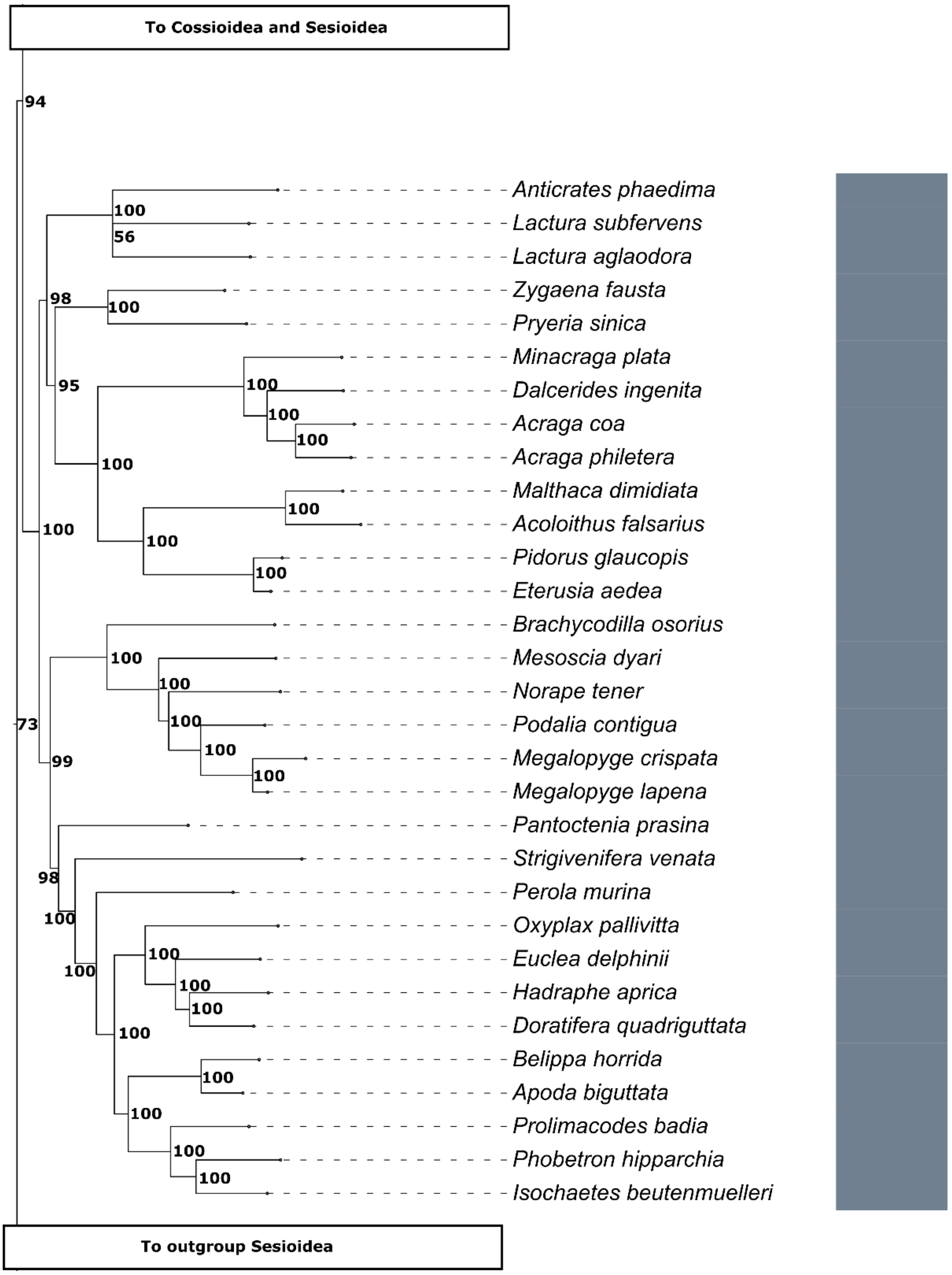


**Figure S22. Phylogenetic view of the Zyganoidea only.**

<u>Papilionoidea (Latreille, 1802) + Hedyloidea (Scoble, 1986)</u>

The Hedyloidea have been shown to be nested within the Papilionoidea since the early 2010's, with increasingly strong evidence demonstrating that Papilionoidea are sister to all other butterflies (Mutanen et al. 2010; Heikkilä et al. 2012; Regier et al. 2013; Kawahara & Breinholt, 2014; Mitter et al. 2017; Kawahara et al. 2023). In our study, these findings are also confirmed with strong support (Figs. 3, S8). We demonstrate the already established phylogenetic relationships between Papilionoidea and Hedyloidea, and within Hedyloidea (Mitter et al. 2017; Fig. S8).

Pyraloidea (Latreille, 1809)

The monophyletic status of Pyraloidea is strongly supported by both morphological (Heikkilä et al. 2015) and molecular studies (Regier et al. 2009; Mutanen et al. 2010; Regier et al. 2012B; Kawahara et al. 2019; Fig. 3). This study agrees with the previous findings of Minet (1982, 1985) that the superfamily Pyraloidea can be split into two monophyletic families, the Crambridae and the Pyralidae (Fig. S23). We find mostly very strong bootstrap support in our results (68 – 100, Figs. 3, S23) for the taxonomic placement of the Pyraloidea in this work. We also find multiple previous studies which support our findings (Minet 1982 & 1985; Regier et al. 2009; Mutanen et al. 2010; Regier et al. 2012B; Kawahara et al. 2019).

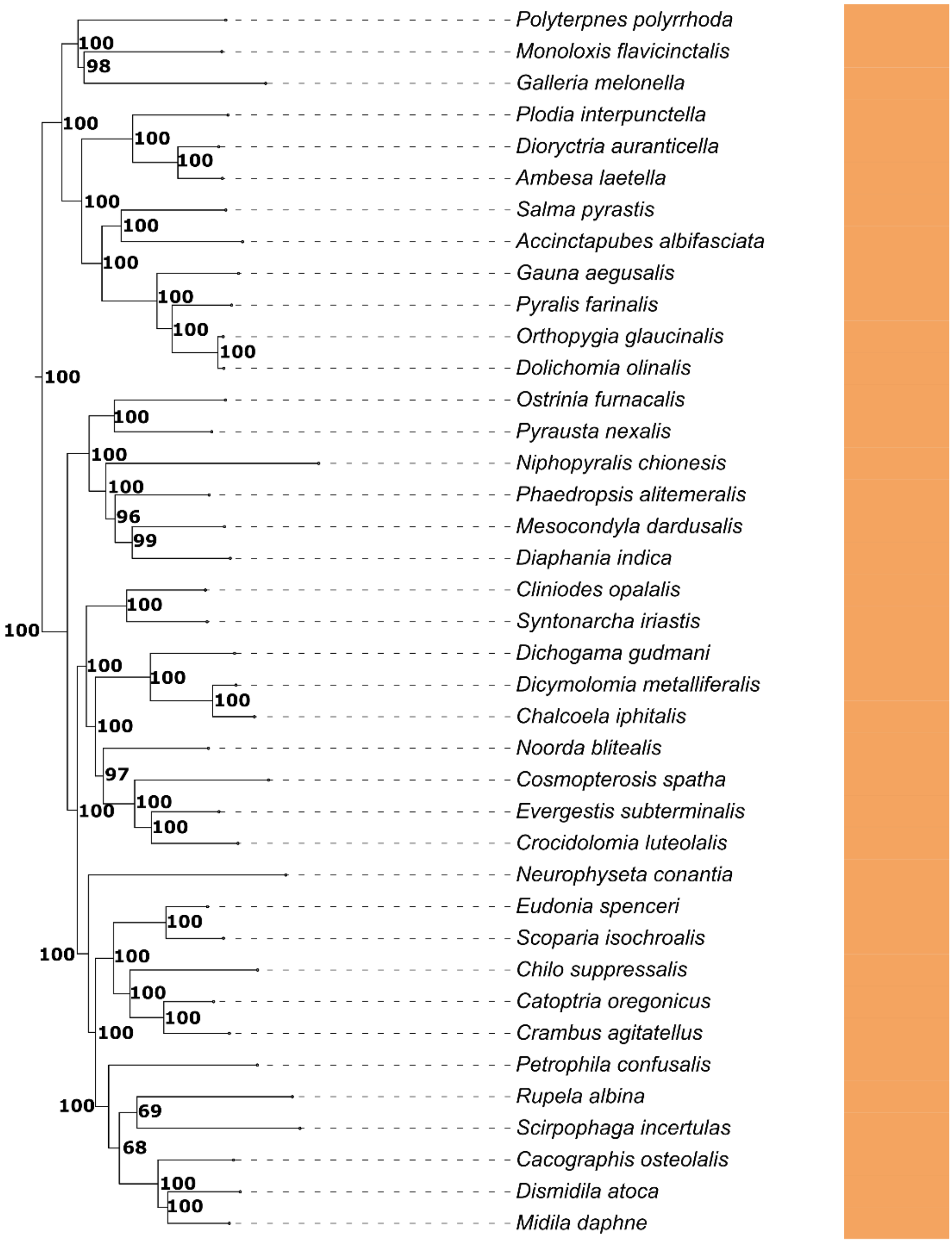


**Figure S23. Phylogenetic view of the Pyraloidea only.**

Geometroidea (Leach, 1815)

This superfamily is often placed as close to the Bombycoidea + Lasiocampoidea complex (Mitter et al. 2017; Kawahara et al. 2019). Alternatively, it has been placed at the base of the Macroheterocena with the closest relatives being the Drepanoidea (Heikkilä et al. 2015). In this study, we do not show the Noctuoidea and Geometroidea as sister clades, basal to the Macroheterocena with Drepanoidea found between the two (Figs 3, S24-26). Instead, our results show an intermediary result between Heikkilä et al. (2015) and others (Mitter et al. 2017; Kawahara et al. 2019).

Within the Geometroidea, we also find evidence supporting the placement of the family Semanturidae (Figs. 3, S24). Our results also support the previously demonstrated inclusion of the Epicopeiidae in Geometroidea (94-100, Fig. S24). The phylogenetic placement of this family has oscillated between the Drepanoidea and the Geometroidea over time (Kristensen & Schmidt-Rhaesa, 1998; Bazinet et al. 2013; Regier et al. 2013; Mitter et al. 2017; Figs. 3, S24). Our study supports multiple previous studies indicating that Geometroidea, including Epicopeiidae, is monophyletic (Kristensen & Schmidt-Rhaesa, 1998; Bazinet et al. 2013; Regier et al. 2013; Mitter et al. 2017; Kawahara et al. 2019. Figs. 3, S24). More research is recommended with regards to the placement of the Epicopeiidae.

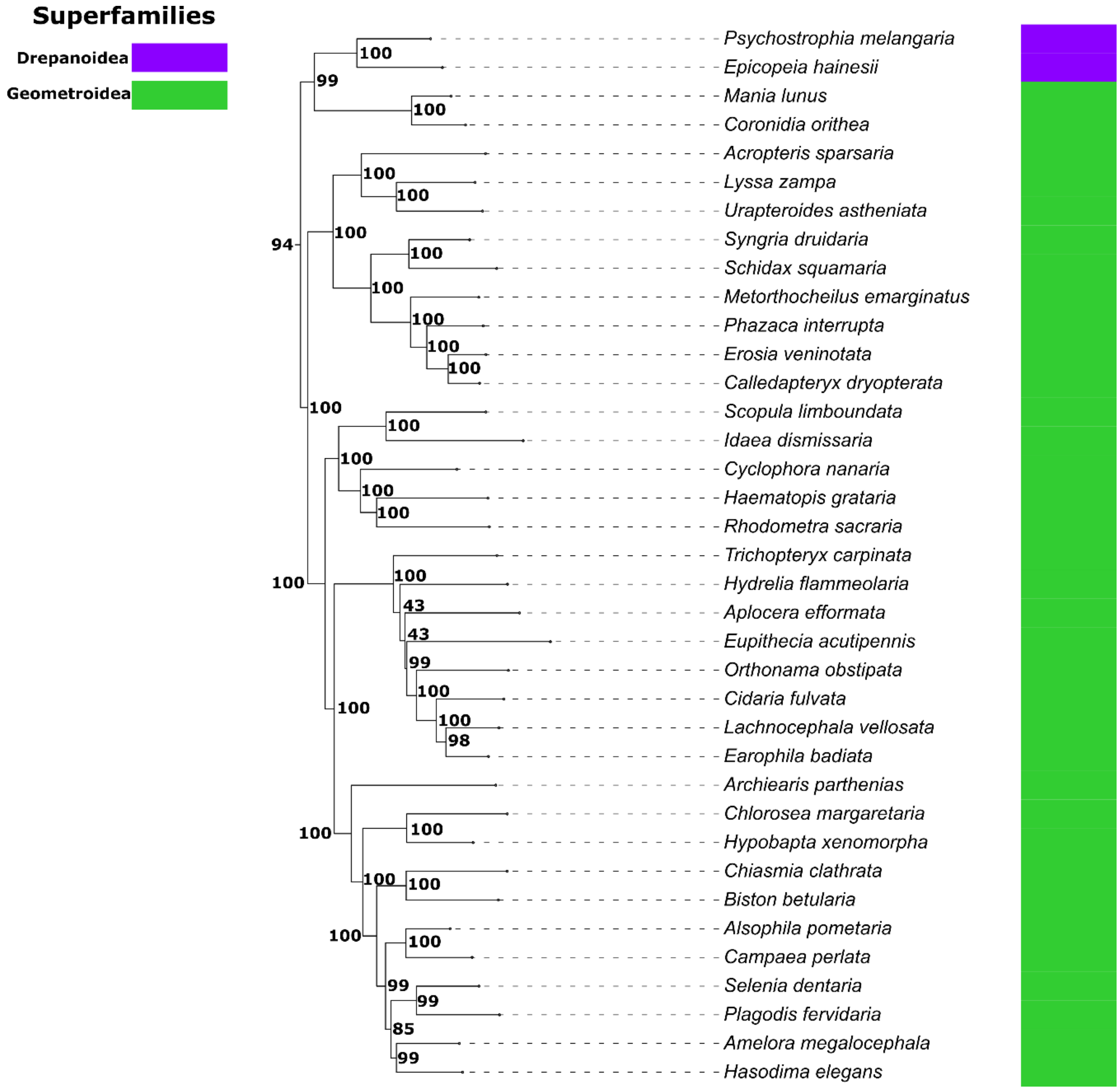


**Figure S24. Phylogenetic view of the Geometroidea.** The Drepanid moths *Psychostrophia melangaria* and *Epicopeia hainessi* are also observed, clustering with the Semanturidae and basal to all other Geometroidea.

Drepanoidea (Boisduval, 1828)
In this study we found that the Drepanoidea are (with the exception of two species clustering with the Geometroidea, Figs, S24, S25) monophyletic and placed within the Macroheterocerna (Fig. 3), as demonstrated in previous studies (Heikkilä et al. 2015, Mitter et al. 2017, and Kawahara et al. 2019).

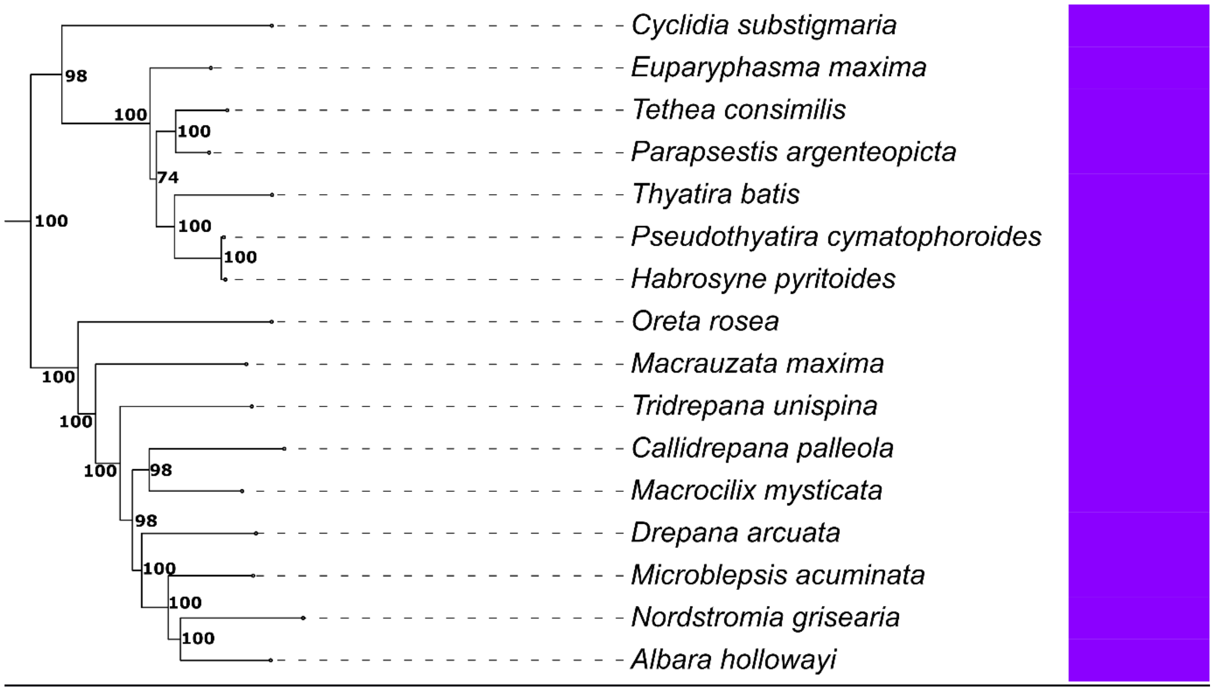


**Figure S25. Phylogenetic view of the Drepanoidea only.**

Noctuoidea (Latreille, 1809)
In many recent phylogenies, the placement of the Noctuoidea varies, the superfamily is sometimes placed next to the Bombycoidea although other placements have also been proposed (Heikkilä et al. 2015; Mitter et al. 2017; Kawahara et al. 2019). In our study Noctuoidea (Fig. S26) are found in the middle of the Macroheterocerna, between the Lasiocampoidea-Mimallonoidea group and the Drepanoidea/Geometroidea (Figs. 3, S24-S27).

We find that Noctuoidea is strongly supported as monophyletic, as in multiple previous studies (Regier et al. 2009; Mutanen et al. 2010; Zahiri et al. 2011; Regier et al. 2013; Figs. 3, S26). Our taxonomic sampling of Noctuoidea was composed primarily by three large families (Noctuidea, Notodontidae, and Eribidae), with only a handful of taxonomic representatives from other families, poorly sampled in this work (Micronoctua (Micronoctuidea)- 1sp, Oreonsandra (Oenosandridae)- 1sp and *Earias, Belenina, Maganola, Iscadia* & *Negeta* – 5sp (Nolidae), Table S2; Fig. S26).

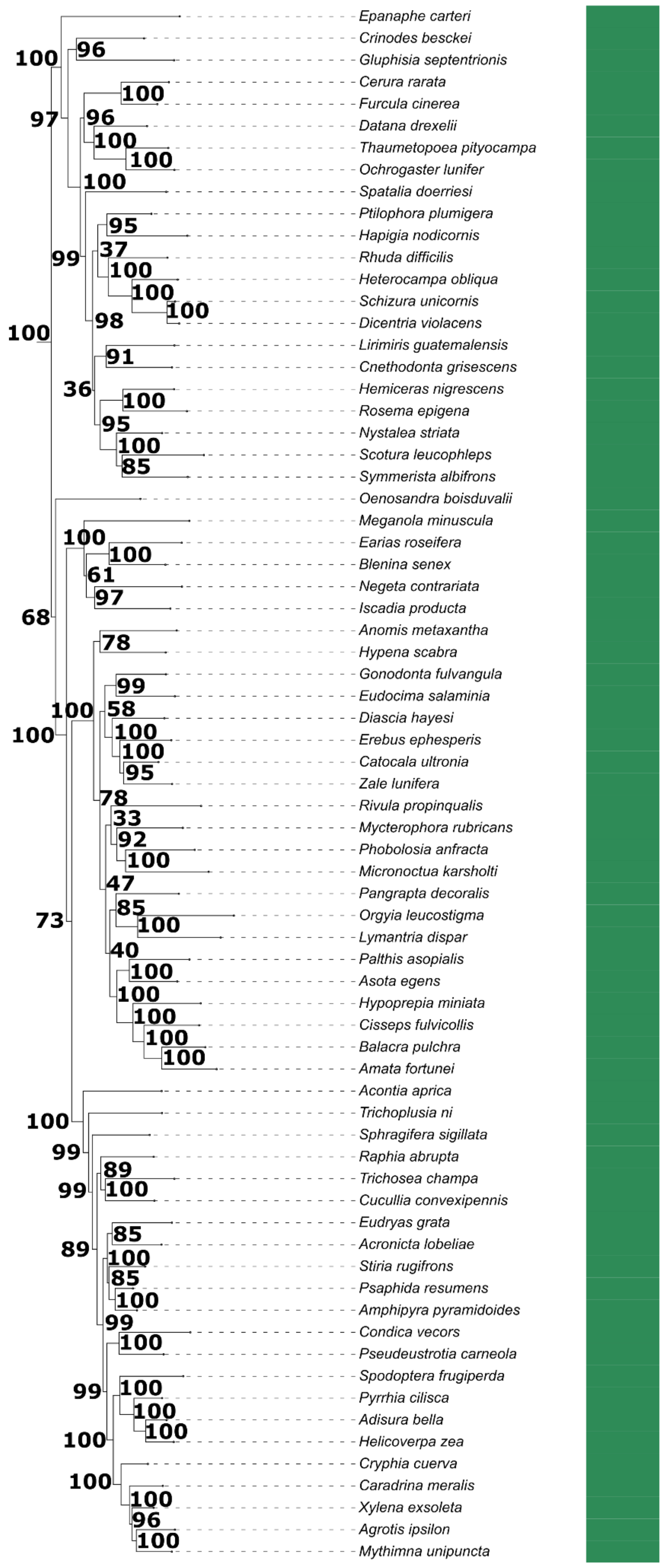


**Figure S26. Phylogenetic view of the Noctuoidea only.**

Mimallonoidea (Minet, 1983)
We find that the superfamily Mimallonoidea and the Cimeliidae form a clade within the Macroheterocerna (Fig. S27). Previous studies have argued for the inclusion of Cimeliidae within the Drepanoidea (Van Nieukerken et al. 2011). However, in this study our sole Cimeliidae species (*Axia margarita*) clusters with the Mimallonoidea and not the Drepanoidea with strong bootstrap support (94, Fig. S27). We recommend further research regarding the placement of *A. margarita* and the Cimeliidae as a whole.

The Mimallonoidea in previous research are placed variably. Sometimes this superfamily is placed outside the Macroheterocerna (Heikkilä et al. 2015; Kawahara et al. 2019) and sometimes within (Mitter et al. 2017; Figs. 3, S27). In this study, we have strong support for the placement of Mimallonoidea as a monophyletic group within the Macroheterocerna (Fig. 3).

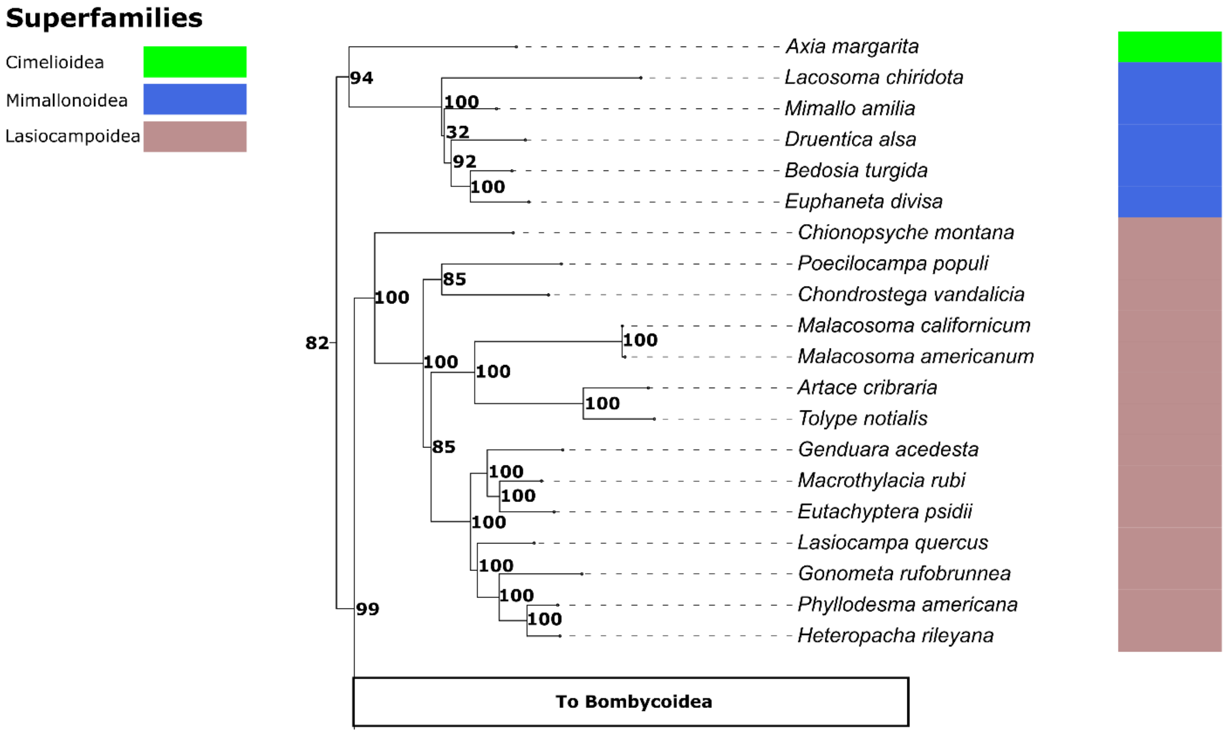


**Figure S27. Phylogenetic view of the Lasiocampoidea, Mimallonoidea and Cimelioidea only.**

Bombycoidea (Gravenhorst, 1843) complex and Lasiocampoidea (Latreille, 1802)
The placement of the Bombycoidea is consistent across phylogenetic studies including our own. Datasets which span multiple superfamilies regularly place the Bombycoidea as the terminal taxa (Heikkilä et al. 2015; Mitter et al. 2017; Figs. 3, S9A-C) although, this is not always the case (Kawahara et al. 2019). Bombycoidea is consistently placed as sister to the Lasiocampoidea (Bazinet et al. 2013; Regier et al. 2013; Heikkilä et al. 2015; Mitter et al. 2017; Kawahara et al. 2019; Figs. 3, S9A-C, S27). We recovered both Lasiocampoidea and Bombycoidea as individually monophyletic superfamilies with the Bombycoidea sister to Lasiocampoidea (Fig. 3, S9A, S27).

Within the Bombycoidea, the family Bombycidae and its subfamilies may not be monophyletic. Taxa currently assigned the subfamily Apatelodinae clustered with taxa from the superfamilies Eupterotidae, Lemoniidae, Brahmaeidae, with strong support (100, Fig. S9A-C). Similarly, some taxa in the Bombycidae subfamily Prismostictinae were recovered as more closely related to Carthaeidae, Endromidae, Mirinidae, Phiditidae, Anthelidae than to other Bombycidae (Fig. S9A-C). As in previous studies (Heikkilä et al. 2015; Kawahara et al. 2019), the Sphingidae and the Saturniidae were recovered as sister clades (Fig. S9A-C).

### Smaller superfamilies and their phylogenetic affinities

Epermenoidea (Minet, 1983)
We only sampled three species of this superfamily (Fig. S8). The Epermenoidea are small tuft moths which were previously placed in either the Copomorphoidea or the Yponomeutoidea, based on physical characteristics or molecular phylogenetics (Dugdale et al. 1999, Sohn et al. 2013), but more recently have been elevated to the level of

superfamily with the aid of molecular phylogenetics (Heikkilä et al. 2015; Mitter et al. 2017; Fig. 3). Our results support of Mitter et al. (2017) and Heikkilä et al. (2015), indicating that the Epermenoidea are Nonobtectomeran Apoditrysia (Figs. 3, S9), and support the idea that Epermenioidea are their own taxonomic unit at the level of the superfamily. This phylogenetic placement warrants further investigation.

Galacticoidea (Minet, 1986)
The superfamily Galacticoidea is sparsely sampled in this work (one species), forming a group with the inclusion of the Douglassid Gracillarioidea, the Cossid *Phycodes toulgoetella* and the Zyganid *Himanopterus fuscinervis* (Figs. 3, S18). The Galacticoidea clustering with taxa regarded as being *incertae cedis* is potentially due to their low sample size and thorough investigations into this clustering of species is advised before any meaningful conclusions can be drawn from this grouping (Figs. 3, S18).

Immoidea (Common, 1979)
The Immoidea sparsely sampled in this work formed a group (sister to the main Cossioidea, Sesioidea and Zyganoidea) with the inclusion of the Parasitic Zyganoidea with strong support (100, Figs. 3, S21). The clustering of the Immoidea with taxa regarded as being *incertae cedis* (parasitic Zyganoidea of the genera *Fulgoraecia* and *Epipomponia*, see above) is potentially due to their low sample size and thorough investigations into this clustering of species is advised before any meaningful conclusions can be drawn from this grouping (Figs. 3, S21).

Calliduloidea (Moore, 1877), Copomorphoidea (Hampson, 1918), Hyblaeoidea (Hapmson, 1893) and Thyridoidea (Swinhoe, 1892)
These four superfamilies are represented by a handful of species in this work (Fig. 3, Table S2) and do not appear frequently in molecular studies. However, their placement on the tree is an interesting result, most notably because they have clustered together with medium to strong node support (48-100, Figs. 3, S8). Previous findings by Kawahara et al. (2019), demonstrate evidence of some relatedness between these superfamilies, notably Kawahara et al. (2019) observe that the Calliduloidea and Thyridoidea clustered together. Other studies have however found no evidence for the phylogenetic relatedness of these superfamilies (Heikkilä et al. 2015). Finally, one Pterophorid (*Emmelina monodactyla*) clusters separately with the Choreutidae, Pterophoridae and Urodidae + Douglassid Gracillarioidea (Figs. 3, S18) and not with its counterparts (Figs. 3, S8). The placement of all these species/superfamilies requires further investigation.

**Taxonomic summary of the Major superfamilies**

In summary, we find strong evidence for the monophyly of the following superfamilies: Noctuoidea, Geometroidea, Thyridoidea, Pyraloidea, Lasiocampoidea, Yponomeutoidea, Gelechioidea, Mimallonoidea, Papilionoidea + Hedyloidea and Bombycoidea (Fig. 3), consistent with multiple previous studies (Heikkilä et al. 2015; Mitter et al. 2017; Kawahara et al. 2019). We also include (albeit small) several representative species of Lepidopteran superfamilies which rarely (if at all) appear in large scale multi-superfamily phylogenetic trees (Hyblaeoidea, Pterophoroidea, Urodoidea, Choreutoidea, Immoidea, Galacticoidea, Epermenoidea, Calliduloidea, Copomorphoidea & Cimelioidea. Figs. 3, S8, S18, S21, S27). The placement of these rarely included taxonomic groups is important if we are to build the full picture of Lepidopteran evolution. When it comes to the phylogenetic placement of otherwise rarely included superfamilies, we offer suggestive avenues for future research through the support we obtained in our tree analyses. We urge that further research be conducted to include more species from these taxonomic groups.

Finally, we provide strong evidence and further support for the revision of two distinct superfamilies: the Cossioidea-Sesioidea complex and the Tineoidea complex (Figs. 3, S14, S21). Our findings contribute to ongoing efforts to resolve the phylogenetic relationships of these poorly known groups (Mitter et al. 2017).

**Novel placements of individual species and small taxonomic groups, considered *incertae cedis***

Species assigned to the Zyganoidea
The ectoparasitic species *Epipomponia nawai*, *Fulgoraecia exigua (Epipyropidae)* currently classed as Zyganoidea were found as sister to an unexpected superfamily, the Immoidea (Figs. 3, S21). Another species currently placed in

the Zyganoidea (*Himantopterus fuscinervus*) clustered with the Galacticoidea + Heliocosma and the Cossid *Phycodes toulgoetella* (Fig. S18). However, in works which include morphological data, ectoparasitic species of Zyganoidea were placed with varying support as sister to the Limacodidae (Heikkilä et al. 2015; Mitter et al. 2017). Further investigation is required to resolve the phylogenetic placement of these taxa.

*Phycodes toulgoetella*

*Phycodes toulgoetella* placed alongside the Galacticoidea is a novel placement which to our knowledge is not reported before (Fig. S18). We report this placement with some support (49-56) but a specific taxonomic investigation at the genetic level is recommended.

*Cycloplasis panicifoliella*

*C. panicifoliella* was moved out of Heliodinidae, but still within the Yponomeutoidea by Hsu and Powell (2005). Then this genus was then removed entirely from the Yponomeutoidea by Sohn et al. (2013), who instead placed this genus as *incertae cedis* in clade composed of Apoditrysia + Gelechioidea. We find evidence to support the core finding made by Sohn et al. (2013), that *C. panicifoliella* does belong in the Apoditrysia but not within the Gelechioidea group. Additionally, unlike Sohn et al. (2013), this work, places *C. panicifoliella* as sister to the small superfamily Urodoidea (Figs. 3, S17) whereas Sohn et al. (2013) did not recover the same relationship and instead, they found *C. panicifoliella* as sister to a member of the superfamily Galacticoidea. We recommend that additional phylogenetic studies include larger numbers Urodoidea and Galacticoidea to ascertain their relationships and the phylogenetic placement of C. *panicifoliella.*

*Heliocosma* and allies

Our study (Figs. 3, S17) recovered found the genus *Heliocosma* within their own group. This is not consistent with Mutanen et al. (2010) who found these species to be within the Cossioidea-Sesioidea complex (S21 in this study). The *Heliocosma* in this study (two species) cluster with a small group of species (Figs. 3, S18). This small group appears to be sister to all other nonobtectomera and is composed of the Galacticoidea (1spp), the Cossid *Phycodes toulgoetella* and the Zyganid *Himantopterus fuscinervis*. Our evidence for the inclusion of the *Heliocosma* within this tiny group adds further evidence for a review of the Cossioidea-Sesioidea complex and allies/associated species including the Galacticoidea.

Douglassidae (currently *incertae cedis*)

Placement of the Douglassidae varies across works (Regier et al. 2009; Mutanen et al. 2010; Kawahara et al. 2011; Regier et al. 2013, Table S2, Figs. 3) and in our case, the Douglassidae are placed with the Pterophorid *Emmelina monodactyla* (Fig. S17) as sister to the Urodoidea, the Choreuthid *Milleria dolosalis* and the Yupometoid/*incertae sedis* species *Cycloplasis Panicifoliella*. The clustering of the Douglassidae with these other species is novel (Fig. S17). Further work is required to determine the phylogenetic affinities and validity of this grouping.

*Parornix torquillella*

We sampled two species of *Parornix* (*P. torquillella* & *P. anglicella*) which were not recovered as a clade in our phylogenetic analysis (Figs. 3, S15, S16). *P. anglicella* remains in its currently accepted taxonomic position (within the Gracillarioidea), while *P. torquillella* is placed in the Yponomeutoidea, both with strong bootstrap support for their positioning on the tree (74-100, Figs. S15, S16). This result is unexpected and calls for further investigations with a larger sample of *Parornix,* Yponomeutoidea, and Gracillarioidea.

*Milleria dolosalis*

This species, originally assigned to the superfamily Choreutoidea (Heppner, 1982) finds itself clustering with species otherwise considered to be in *incertae cedis* or novel placement i.e. the Douglassidae, the Pterophorid *Emmelina monodactyla*, *Cycloplasis Panicifoliella* and the Urodoidea (Figs. 3, S17). We suggest larger sampling and further investigations to further understand what is occurring with these taxa.

*Agdistopis sinhala*

*Agdistopis sinhala* (originally Pterophoroidea, Pterophoridae, Macropiratinae) is placed in this study within a group consisting of the superfamilies Hyblaeoidea and Thyridoidea (Fig. 3, S8). Previously, Kawahara et al. (2018) found evidence for the placement of the genus *Agdistopsis* with the following superfamilies: Thyridoidea + Calliduloidea, Alucitoidea, Carposinoidea (in this work the Junior synonym of Copomorphoidea is used), Epermenoidea,

Hyblaeoidea. In this work while we recover the Calliduloidea, Alucitoidea, Carposinoidea and Epermenoidea in proximity but separate, we find strong support for *A. sinhala* as sister to the Hyblaeoidea, the two forming a group sister again to the superfamily Thyridoidea with strong bootstrap support (70-100, Figs. 3, S8).
